# Novel antiviral interferon sensitive genes unveiled by correlation-driven gene selection and integrated systems biology approaches

**DOI:** 10.1101/2021.04.05.438481

**Authors:** Cristina Cheroni, Lara Manganaro, Lorena Donnici, Valeria Bevilacqua, Raoul J.P. Bonnal, Riccardo L. Rossi, Raffaele De Francesco

## Abstract

Interferons (IFNs) are key cytokines involved in alerting the immune system to viral infection. After IFN stimulation, cellular transcriptional profile critically changes, leading to the expression of several IFN stimulated genes (ISGs) that exert a wide variety of antiviral activities. Despite many ISGs have been already identified, a comprehensive network of coding and non-coding genes with a central role in IFN-response still needs to be elucidated. We performed a global RNA-Seq transcriptome profile of the HCV permissive human hepatoma cell line Huh7.5 and its parental cell line Huh7, upon IFN treatment, to define a network of genes whose coordinated modulation plays a central role in IFN-response. Our study adds molecular actors, coding and non-coding genes, to the complex molecular network underlying IFN-response and shows how systems biology approaches, such as correlation networks, network’s topology and gene ontology analyses can be leveraged to this aim.

## Introduction

The hepatitis C virus (HCV) is a positive-sense single-stranded RNA virus belonging to the family of *Flaviviridae*. HCV is the causative agent of hepatitis C and the major cause of hepatocellular carcinoma. HCV infection potently stimulates the production of endogenous IFNs, which leads to ISG up-regulation in infected hepatocytes (Heim & Thimme, 2014).

The study of HCV in the laboratory has been particularly challenging over the years because of the narrow host range (it infects only humans and chimpanzees) and due to the paucity of cell lines permissive to HCV replication. Most of the cell systems tested over the years, in fact, did not support viral replication. A Huh7 derived clone termed Huh7.5 was found to be susceptible to HCV infection and highly permissive for HCV replication. This cell line was rapidly established as a robust and widely used in vitro infection model for HCV. Subsequent studies revealed that Huh7.5 permissiveness to HCV replication could possibly be a consequence of a single point mutation in the dsRNA sensor retinoic acid-inducible gene-I (RIG-I) (reviewed in Bartenschlager & Pietschmann, 2005). RIG-I is a pattern recognition receptor that senses non-self viral nucleic acids. Specifically RIG-I recognizes pathogen-associated molecular patterns (PAMPs) characterized by a short 5′ triphosphate uncapped double stranded or single stranded RNA. After binding to its PAMP, RIG-I starts a signaling cascade that leads to the activation of transcription factors IRF3 and NF-κB. These two factors activate the transcription of type I and III interferons as well as other antiviral genes that induce a general antiviral state of the infected cells (Kell & Gale, 2015). Notwithstanding the possible molecular defects in the RIG-I sensor, however, treatment of Huh7.5 with type I IFNs potently inhibit replication of sub genomic RNA HCV replicons (reviewed in https://pubmed.ncbi.nlm.nih.gov/14638404/).

Interferon is a class of primordial cytokines that plays a crucial role in the cellular response to the presence of viral pathogens. There are three classes of IFN, type I, II and III and each class comprises different members. Type I IFNs are constituted by IFN-β, IFN-ε, IFN-ω and 13 different subtypes of IFN-α, type II IFN group is composed by only one member named IFN-γ and type III IFN family comprises IFN-λ1, IFN-λ2 IFN-λ3 and IFN-λ4. All IFNs signal through the JAK–STAT pathway, type I and type III IFN share similar antiviral functions, while the type II IFN functions as a pro-inflammatory and immunomodulatory molecule. Type I and III IFN engage two different heterodimeric receptors: the type I IFN receptor (IFNAR) is composed of subunits IFNAR1 and IFNAR2 while type III IFNs receptor (IFNLR), comprises of IFNLR1 (also termed IL28Rα) and IL10Rβ. Upon interaction with its receptor, both type I and III IFN activate the JAK-STAT signaling pathway, leading to the phosphorylation of STAT1 and STAT2 and interaction with IRF9 protein to form a trimeric complex named transcription factor complex ISGF3 (Stanifer *et al*, 2019). The complex migrates into the nucleus and binds to the IFN-stimulated response elements (ISREs) in the promoter regions of a set of genes, overall defined as interferon-stimulated genes (ISGs) that are responsible for the antiviral state of the cell. The binding of ISGF3 complex to ISREs elements induces the transcription of ISGs and initiates the cellular antiviral response.

Despite binding to different receptors, type I and III IFN trigger the same cascade of intracellular events, therefore having a similar effect on cell physiology (Stanifer *et al*, 2019).

However, the potency and the kinetics of the antiviral response are different. Indeed, type III IFN response is delayed and less potent compared to type I IFN induced, but more sustained over time. In addition, the expression pattern of IFNAR and IFNLR is distinct, with IFNAR being expressed on the surface of all cell types, while IFNLR has been observed to be expressed mainly by epithelial cells and neutrophils. IFNLR has also been shown to be expressed by human hepatocytes and that HCV seems to preferentially stimulate IFN-λ production rather than type I IFNs. Of note, IFNL locus polymorphisms are associated with different clinical outcomes during hepatitis C virus (HCV) infection suggesting the importance of this molecule in the response and pathogenesis related to HCV infection (Lazear *et al*, 2019; Mesev *et al*, 2019; Lazear *et al*, 2015). Long non-coding RNAs are a class of RNA molecules longer than 200 nucleotides that are transcribed, but lack the ability to code for a protein product. Given their increasingly described role in the transcriptional and post-transcriptional regulation of gene expression, long non-coding RNAs have been emerging as a new layer of biological complexity (Mathy & Chen, 2017). While a canonical set of protein-coding genes acting as interferon stimulated genes is well-known (Schoggins, 2019), a comprehensive view on the network of protein coding and long non-coding genes with a central role in IFN response still needs to be elucidated.

Although recently developed therapy regimens have strongly increased the cure rate, infection by hepatitis C is still a major cause of chronic liver disease (Maucort-Boulch *et al*, 2018). In hepatocytes, the infection by HCV induces an interferon response that is aimed at suppressing viral replication. In fact, in cellular models, treatment with both class I and class III interferon molecules is able to suppress HCV replication and several protein-coding genes have been reported to mediate this phenomenon, thus acting as HCV restriction factors (Marcello *et al*, 2006). We chose to use the Huh7.5 cell line along with the parental Huh7 because of the higher permissiveness to HCV infection of the first, and in order to focus also on downstream RIG-1 mediated signaling. So we performed RNAseq on Huh7.5 in the presence or in the absence of IFN treatment including also the parental clone Huh7 in the same conditions.

Each gene is estimated to interact with at least four other genes in the actualization of regulatory networks (Arnone & Davidson, 1997) and to have functional relevance in many biological functions (Miklos & Rubin, 1996). Uncovering correlations among genes is instrumental in deciphering the regulatory rationales of signal transduction pathways. Adopting higher level approaches able to analyze correlations as a whole can bypass the intrinsic limitations of the classical differential expression relying on arbitrary significance or expression cut-offs (Schadt *et al*, 2005). Weighted Gene Correlation Network Analysis (WGCNA) provides a system perspective of gene expression by grouping genes into modules selected according to correlation with a relevant phenotypic or molecular trait; this allows to find and integrate transcriptional patterns without using arbitrary selection cutoffs (Langfelder & Horvath, 2008). Using WGCNA it is possible to correlate traditional gene expression with any external measurement obtained from the same samples set: this cross-domain data integration leverages the discovery of otherwise hidden modules of highly correlated genes that can be used as new candidates or to unveil subtle molecular or biochemical mechanisms (Stuart *et al,* 2003; Langfelder & Horvath, 2007).

In order to obtain a genome-wide view of the network of coding and non-coding RNA genes in IFN treated hepatoma cells and to identify possible novel effectors of IFN antiviral activity we applied to our set of RNA sequencing data a systems biology approach modeled on independent phenotypical pattern of the two well-known IFN sensitive genes (Schneider *et al*, 2014) individually measured in the populations of IFN-treated and untreated hepatoma cell lines. After the identification of differentially expressed (DE) genes of IFN treated and untreated Huh7 and Huh7.5 cells, we constructed a weighted gene co-expression network including both coding and non-coding molecules, gene modules and sub-networks were identified as significantly related to IFN-treatment as a whole. Integrating topological network metrics and gene ontology terms we identified 42 novel hub genes, including 7 long noncoding RNAs (lncRNAs) and 7 pseudogenes, in the modules associated with IFN-response. Notably, putative binding sites for transcription factors involved in the IFN-response (such as IRF9 and STAT1) were also predicted for most of the genes included in the selected modules.

## Results

### IFN treated cells show a robust over expression of hundreds of genes

In order to decipher the genome-wide landscape of the network of protein coding and long-non coding genes mediating interferon antiviral activity in hepatoma cell lines, we profiled by RNAseq the transcriptome of Huh7 and Huh7.5 cells treated with interferon alpha (IFNα), beta (IFNβ) or lambda (IFNλ) for 8 hours. Each treatment was compared to respective mock-treated cells in 6 pairwise comparisons, for a total of 43 samples classified in 12 experimental conditions (**Table 1**). After gene expression quantification in read counts, dimensionality reduction by principal component analysis (PCA) was applied on the complete dataset: samples nicely segregated according to cell line identity and treatment with no outliers, as shown by PCA whose first two components are able to explain more than 90% of the entire dataset variability (**Supplementary Figure 1**).

**Table 1.** Experimental design. The 6 biological comparisons (3 treatments in each cell line) are described with the number of biological replicates (from 3 to 4) for each condition in parenthesis.

| Cell line | Treatment (# of samples) | Control (# of samples) |
| --- | --- | --- |
| Huh7 | IFN alpha (4) | Mock (3) |
|  | IFN beta (3) | Mock (3) |
|  | IFN gamma (4) | Mock (4) |
| Huh7.5 | IFN alpha (4) | Mock (4) |
|  | IFN beta (3) | Mock (3) |
|  | IFN gamma (4) | Mock (4) |

Gene expression was robustly modulated by IFNα and IFNβ treatments, with IFNλ inducing a milder perturbation (Figure 1a) as expected. Differential expression analysis using 0.05 false discovery rate (FDR) and a twofold change (FC) cut-offs showed hundreds of differentially expressed genes, mostly up-regulated by the treatment (**Figure 1a** and **Supplementary Figure 1**). Specifically, the following number of genes were retrieved as significantly differentially expressed (DE) in treated compared to mock-treated cells: 320 for Huh7-IFNα; 263 for Huh7-IFNβ; 79 for Huh7-IFNλ; 252 for Huh7.5-IFNα; 302 for Huh7.5-IFNβ; 78 for Huh7.5-IFNλ. (**Figure 1a**).

**Figure 1.**
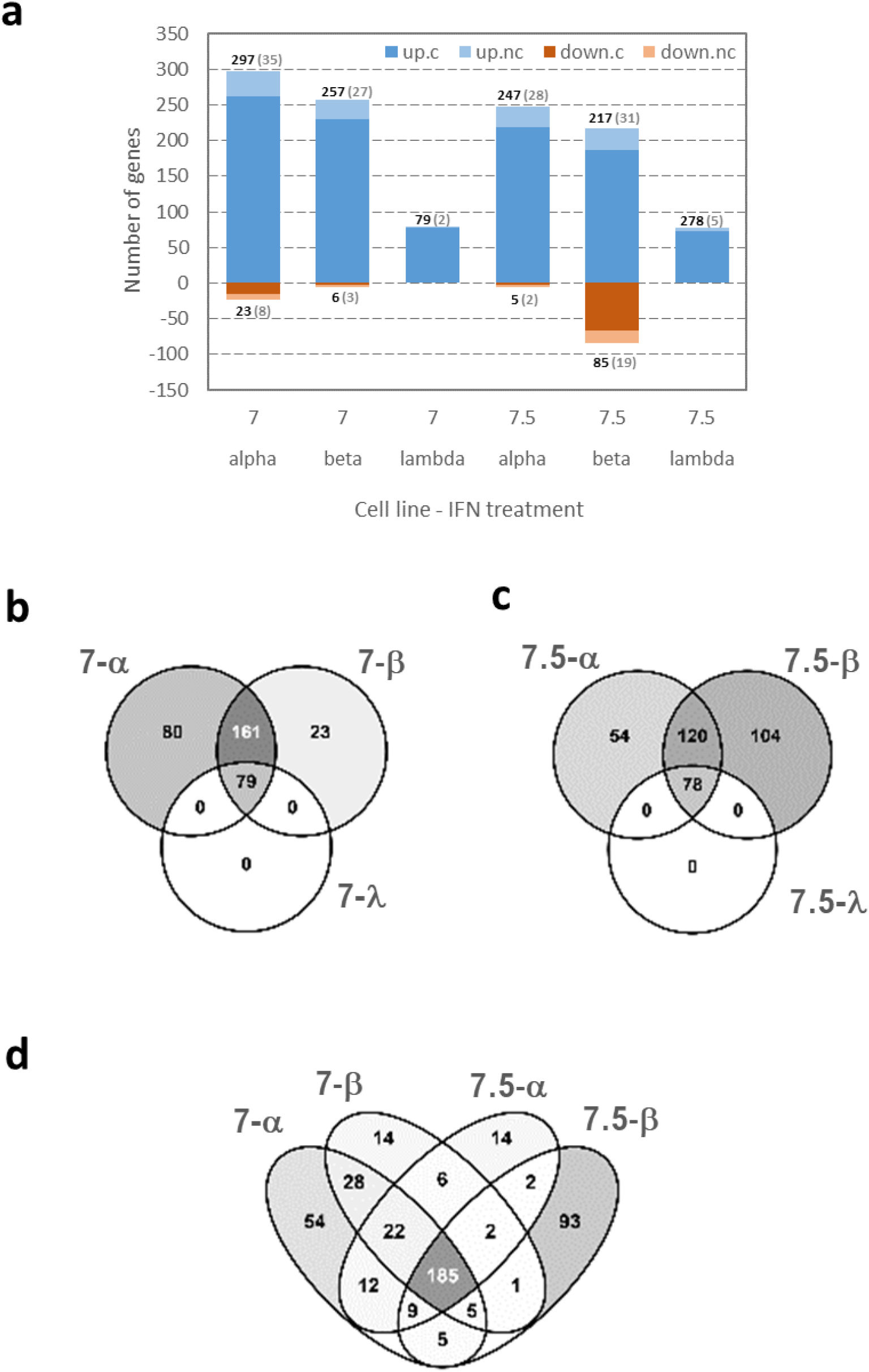
Differential expression analysis on RNASeq transcriptome profiling. **a**. Number of differentially expressed genes that are significantly modulated in each treatment compared to mock-treated cells after DESeq2 analysis with FDR < 0.05 and absolute Log2FC > 1. Over expressed (blue) and downregulated genes (rust red): dark colours for protein coding genes (actual numbers in bold) and light colours for non-coding genes (actual numbers in brackets). **b-d**. Venn diagrams illustrating the overlap of modulated genes across treatments. Panel **b** shows the overlap between IFNα and IFNβ treatments in the Huh7 line, while panel **c** shows the same results for Huh7.5. In both lines, the genes modulated by IFNλ are entirely included in the overlapping region of other two Interferons. Panel **d** shows how all genes in **b** and **c** overlap, showing a group of 185 genes modulated in all cell lines and by both type I IFN perturbations.

Comparing results (numbers of DE genes) of treatments with different IFNs, the fraction of DE genes shared by all contrasts (i.e. observed in all cases) was about 20% (23 and 21.9 in Huh7 and Huh7.5 cells respectively) with IFNλ-modulated genes completely included in the overlapping region for both Huh7 and Huh7.5 (**Figure 1b,c**). A group of 185 genes was consistently modulated by type I IFNs and in both cell lines (**Figure 1d**): these genes are all redundantly involved in IFN-related functions, as confirmed by a scan of the biological processes with Gene Ontology, where mostly high level terms such as “immune system process”, “response to stimulus”, “defense to response”, “innate immune response” and “cytokine production” were overrepresented (not shown).

Taking into account the differentially expressed genes after IFN treatments (Figure 1a), we assessed the functional relevance of cell lines specific subsets of these genes. Comparing genes perturbed by the IFN treatment in Huh7 and Huh7.5, we found that genes that were differentially expressed in both Huh7 and Huh7.5 cell lines (the Venn overlaps and middle panels in Supplementary Figure 2a and b) were the only ones triggering relevant ontologies and pathway enrichments, for both IFNα and IFNβ. Genes that were perturbed exclusively in Huh7 cells enriched a few general terms, while genes perturbed exclusively in Huh7.5 cells did not trigger any significant enrichment. This is an indication that the two cell lines appeared functionally homogeneous according to ontologies and pathways’ enrichment analyses after IFN treatments. Most DE genes were protein-coding, but also some non-coding RNAs (lncRNA) were identified (**Figure 1a** and **Supplementary Figure 1**). Specifically, lncRNA represented the 13.4% of all IFNα induced genes in Huh7 and the 11.9% in Huh7.5. Analysis of IFNβ stimulation dataset revealed that the lncRNA constituted the 11.4% and 16.5% of induced genes in Huh7 and Huh7.5 respectively.

On the other end, IFNλ stimulated lncRNA represented only the 2.5% and 6.4% of induced genes in Huh7 and Huh7.5 respectively. All differentially expressed gene lists at the set cut-off values can be found in **Supplementary File 1**.

### Two gene expression modules among the IFN-related co-expression network are specifically associated with IFN response

Conventional differential expression analysis has several shortcomings when applied to a complex experimental design composed by multiple comparisons, mostly related to experimental noise masking lower intensity signals that collectively can be biologically relevant in terms of regulation. Since we wanted to select genes associated to a continuous effect such as IFN response and at the same time overcome the limitations of a conventional pairwise comparison, we applied weighted gene co-expression network analysis (WGCNA) (Langfelder & Horvath, 2008) that is able to group genes into modules with a similar expression pattern and then classifying modules according to their correlation to a continuous independent external variable, usually connected to the phenotypic perturbation under investigation. This method allowed us to summarize the large number of profiled genes in a small set of gene modules exemplifying the transcriptional behaviors in response to all IFN treatments. Instead of analyzing the treated-untreated contrasts separately in the two cell lines and for the three IFN treatments, as it is usually done in differential expression, we applied the WGCNA framework on normalized and appropriately transformed read count values of 6168 genes (selected on the basis of the coefficient of variation) across all 43 samples in our experimental design. By employing a gene clustering strategy on the Topological Overlap Dissimilarity Measure, we identified 12 gene modules with size ranging from 65 genes in the Tan module to 1335 genes in the Turquoise module (**Figure 2**). 1267 genes remained unassigned (**Figure 2a**, grey color). Multidimensional scaling confirmed a clear clustering in the three-dimensional space of the genes assigned to the same module (**Supplementary Figure 3a**).

**Figure 2.**
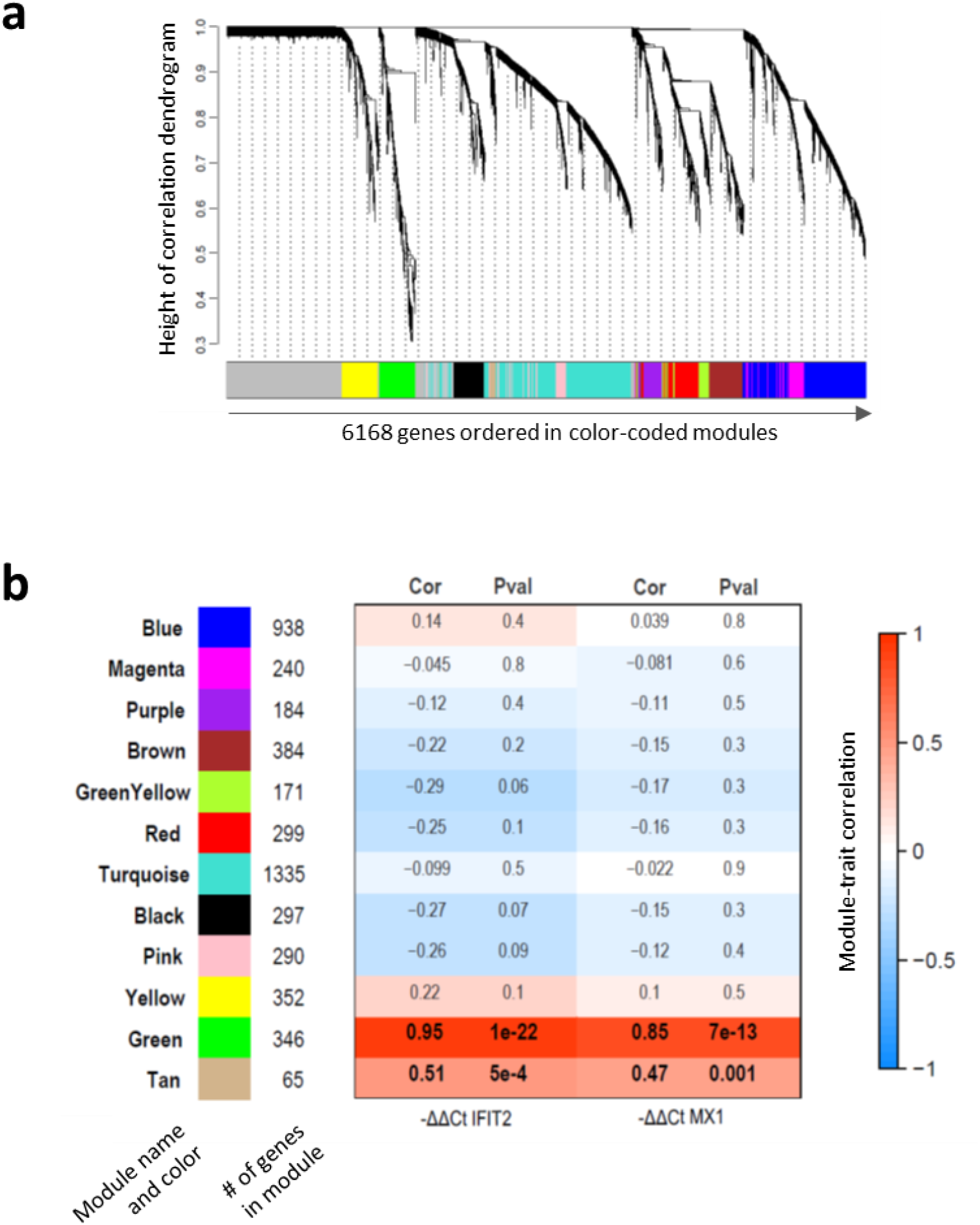
Generation of the gene co-expression network and selection of modules associated with IFN treatment. **a**. Gene dendrogram generated on the Topological Overlap Dissimilarity matrix: each branch of the dendrogram corresponds to a module of highly interconnected genes, for a total of 12 modules identified. The color row below the dendrogram represents the module assignment. **b**. Identified modules with relative colour “names”: numbers beside each colour block indicates the number of genes assigned to each of them. The heat map shows the correlation values (Pearson correlation degree “Cor” and statistical significance “Pval”) between each module eigengene and the induction of IFIT2 and MX1 transcripts measured by real-time PCR. Negative correlations are in blue, positive correlations are in red. Only Green and Tan modules show a significant positive correlation with the ISGs expression.

In order to pinpoint the modules related to IFN response we used the expression pattern of two induced well known IFN sensitive genes (ISGs) (Schneider *et al*, 2014), MX1 and IFIT2, as independent experimental variables to scan the summarized expression profile of each module. To this aim, expression of MX1 and IFIT2 was measured by RT qPCR on the same sample set (**Supplementary Figure 3b**,**c**), and after identification of the expression profile of each module with the WGCNA summarizing metric (its first principal component or module eigengene) we related it to the induction of the two ISGs and selected the modules with a similar expression profile. Two modules (Green and Tan) out of twelve identified were selected due to their strong positive correlation to both IFN response proxies (MX1 and IFIT2 expression), with statistically significant p-Values (**Figure 2b**).

The Green module is composed of 346 genes, that overall shows a remarkable pattern of induction by IFNα and IFNβ (and weaker induction by IFNλ) in both Huh7 and Huh7.5 cells (**Supplementary Figure 4**). The Tan module groups 65 genes that are induced by IFN treatment, but also show differential baseline levels in Huh7 compared to Huh7.5 (basal higher expression in the latter) (**Figure 2b** and **Supplementary File 2**).

The remaining modules, Black, Brown, GreenYellow, Purple, Red and Yellow do not show a clear change of gene expression pattern according to treatment or cell identity. Blue and Magenta modules group genes whose expression levels are lower in Huh7.5 compared to Huh7, while Turquoise and Pink modules are composed by genes expressed at higher levels in Huh7.5 compared to Huh7 cells (data not shown). None of them shows a statistically significant correlation with the induction of the two IFN sensitive genes used (MX1 and IFIT2) (**Figure 2b**).

### Gene modules associated with IFN-response share functional roles and display clear regulatory motifs

Green and Tan modules together count 411 genes, and they do contain almost all the 185 initially selected DE genes while adding many more, witnessing the added value of a selection method not strictly based on hard cutoffs imposed by differential expression. To characterize the two modules, the most informative genes belonging to them can be found using significance measures or topological properties such as high neighborhood connectivity. A functional neighborhood is composed of nodes (genes) that are highly connected (i.e. significantly co-expressed) to a given set of nodes and its analysis facilitates a guilt-by-association screening strategy to prioritize nodes for their potential ability to interact in the biological context under investigation (Stuart *et al*, 2003; Schadt *et al*, 2005). In order to prioritize most relevant genes and to limit noise in subsequent functional analyses potentially triggered by peripheral ones (Yip & Horvath, 2007; Ravasz *et al*, 2002) we applied this procedure and defined modules’ subnetwork using the adjacency matrix (the matrix of co-expression measures, or weights, between all gene pairs) obtained by the WGCNA method: we imported nodes and edges of this matrix into Cytoscape and imposed a threshold of 0.1 as minimum edge weight. Sub-networks of highly connected nodes were thus selected from both the green and tan modules: 282 nodes (255 protein-coding and 27 non-coding genes) and 18085 edges were extracted from the Green module, while 54 nodes (50 protein-coding and 4 non-coding) and 601 edges from the Tan module (**Supplementary File 3**).

Once the modules’ sub-networks of highly-connected (i.e. highly co-expressed) genes were defined, we checked their functional composition with gene ontology (GO) enrichment analysis for biological processes (BP). For Green genes, a remarkable enrichment was observed for categories associated to IFN-response and defense to virus (**Figure 3a**). A similar outcome, although less prominent, was also obtained for the Tan genes (**Figure 3b**). GO enrichment analysis thus confirmed that Green and Tan sub-networks are indeed functionally enriched for genes associated with innate immunity phenomena.

**Figure 3.**
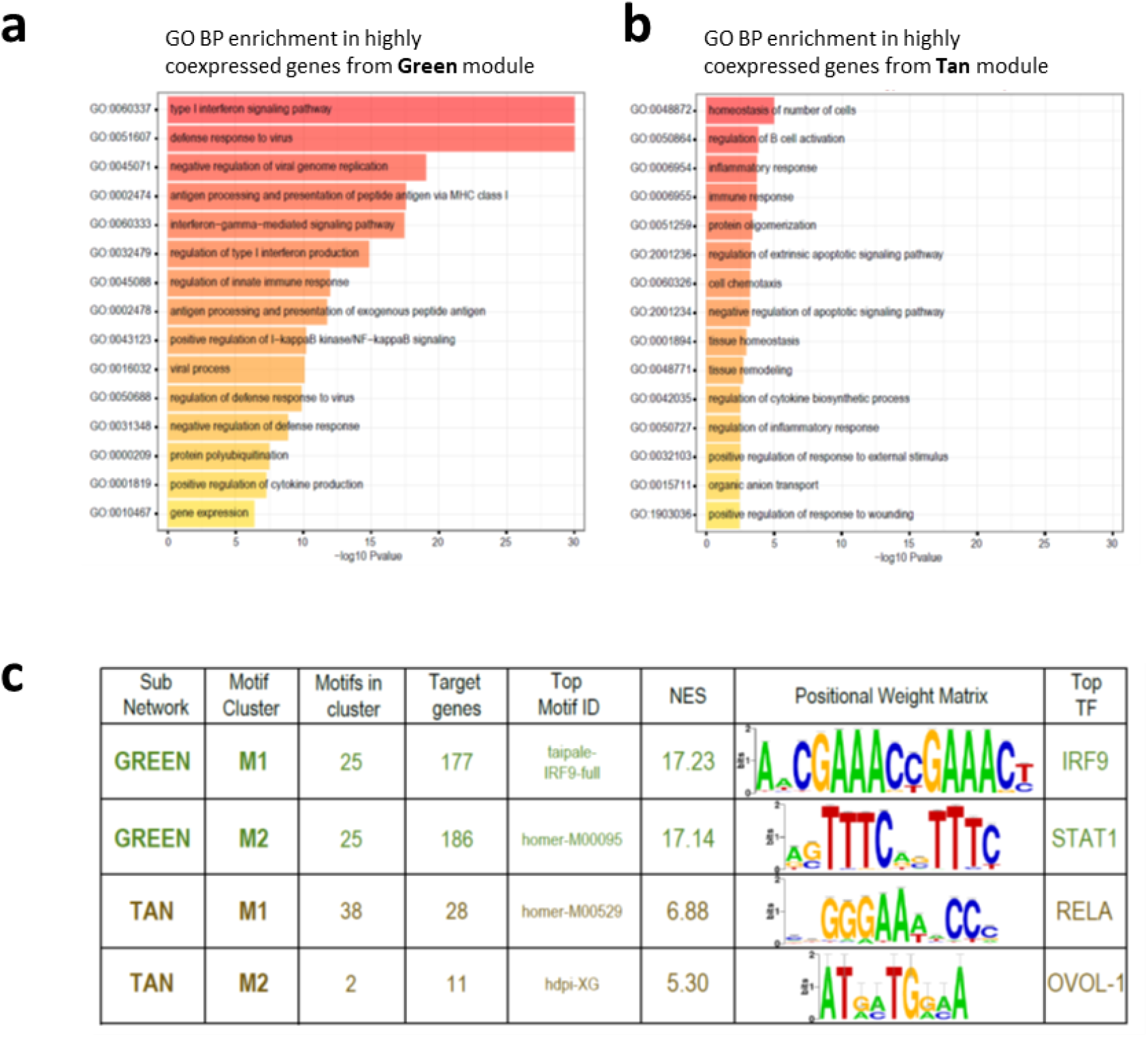
Functional characterization of Green and Tan modules. **a-b**. Results of gene ontology enrichment analysis performed on Green (**a**) and Tan (**b**) sub-networks; bar plots depict the –log p-values for the top-15 Biological Process terms. **c**. Tabular results of the analysis of regulatory motifs performed on Green and Tan sub-networks by IRegulon. The motif clusters identified as significantly enriched for each module (NES > 5) are reported. D-E. Circular plots illustrating the regulatory relationship among each motif cluster and its putative target genes for Green and Tan modules. The thickness of each line and correspondent gene sector represents the number of motifs in the cluster found as putative regulators of the gene.

Co-expressed genes have been successfully used as input datasets for the search of transcription factors (TF) binding sites (Auerbach *et al*, 2013; Yan *et al*, 2013), thus we went further into the functional characterization of our gene modules by inferring their upstream transcriptional regulatory network: to do so we applied the ranking-and-recovery method implemented with the iRegulon motifs collection (Janky *et al*, 2014). Using the iRegulon Cytoscape plugin we searched the Green subnetwork (282 genes) for most probable regulons, i.e. the set of transcription factors and theirs targets, on the basis of presence of shared TF binding sites in the cis-regulatory control region of the genes. We found two highly enriched motif clusters, each composed by 25 regulatory sequences. On the basis of the Normalized Enrichment Score (NES), taipale-IRF9-full was selected as top motif for the first cluster (Green-M1), with IRF9 as top-ranked master regulator. Similarly, homer-M00095 and STAT1 were found respectively as top motif and top regulator for the second cluster, called Green-M2 (**Figure 3c**). Among the 282 genes of the green sub-network that are putative targets, 163 are regulated by both regulators classes (**Supplementary Figure 5a**). Since IRF9-STAT1 axes is the key pathway regulating the IFN response (Stark & Darnell, 2012), the identification of IRF9- and STAT1-related regulons in Green sub-network further highlights its relevance in the cellular response to interferon treatment.

The same analysis was then applied on Tan genes. For this sub-network, two different motif clusters were found to be overrepresented: homer-M00529 (Tan-M1) and hdpi-XG (Tan-M2), with associated master regulators RelA, subunit of NF-kB complex required during the early response to RNA viruses, and OVOL1, respectively (**Figure 3c**). A total of 30 putative target genes were identified, with 9 genes predicted to be the target of both transcription factors classes (**Supplementary Figure 5b**). Interestingly, the expression of OVOL1, a transcription factor involved in cell differentiation seems to be dependent on IRF6, suggesting that OVOL1 responsive genes could be activated at a later time point and play a role in tissue remodeling after a damage induced by the pathogen or the host immune response (Kwa *et al*, 2014).

Taken together these data confirm the ability of our strategy to detect canonical genes involved in IFN response through STAT1-IRF9 pathway as well as novel gene clusters involved in a less characterized IFN modulation.

### Novel potential ISGs are uncovered with network pruning and knowledge subtraction

Network metrics and gene ontology annotations can be exploited to identify the most central players (most connected nodes, or hub nodes) in the network, allowing topological data reduction and facilitating the discovery of not previously annotated key molecular drivers that may not emerge otherwise (**Figure 4a**). We examined more in detail two measures of network centrality in the previously selected sub-network: the eigenvector centrality, which is a measure of the influence of a node in a network, and the closeness centrality, a measure of the proximity of each node to all the others, thus evaluating each node’s contribution to the network structure. The two metrics showed a linear correlation for the highly connected nodes (above a certain threshold), identifying a group of peripheral nodes, for both Green and Tan sub-networks (**Figure 4b**,**c**). The highly co-expressed (and as such, highly connected) genes are likely to play a more relevant role in molecular regulation of IFN induction and when visualized in the network graph they appear as densely connected nodes. The nodes with lower eigenvector (we set an arbitrary cut-off at 0.02) have lower connectedness/co-expression and indeed appeared topologically peripheral in the network graphs (**Supplementary Figures 6-7**). Using this cut-off, we identified as peripheral and discarded 96 and 6 nodes from the Green and Tan sub-networks respectively (**Figure 4b,d**).

**Figure 4.**
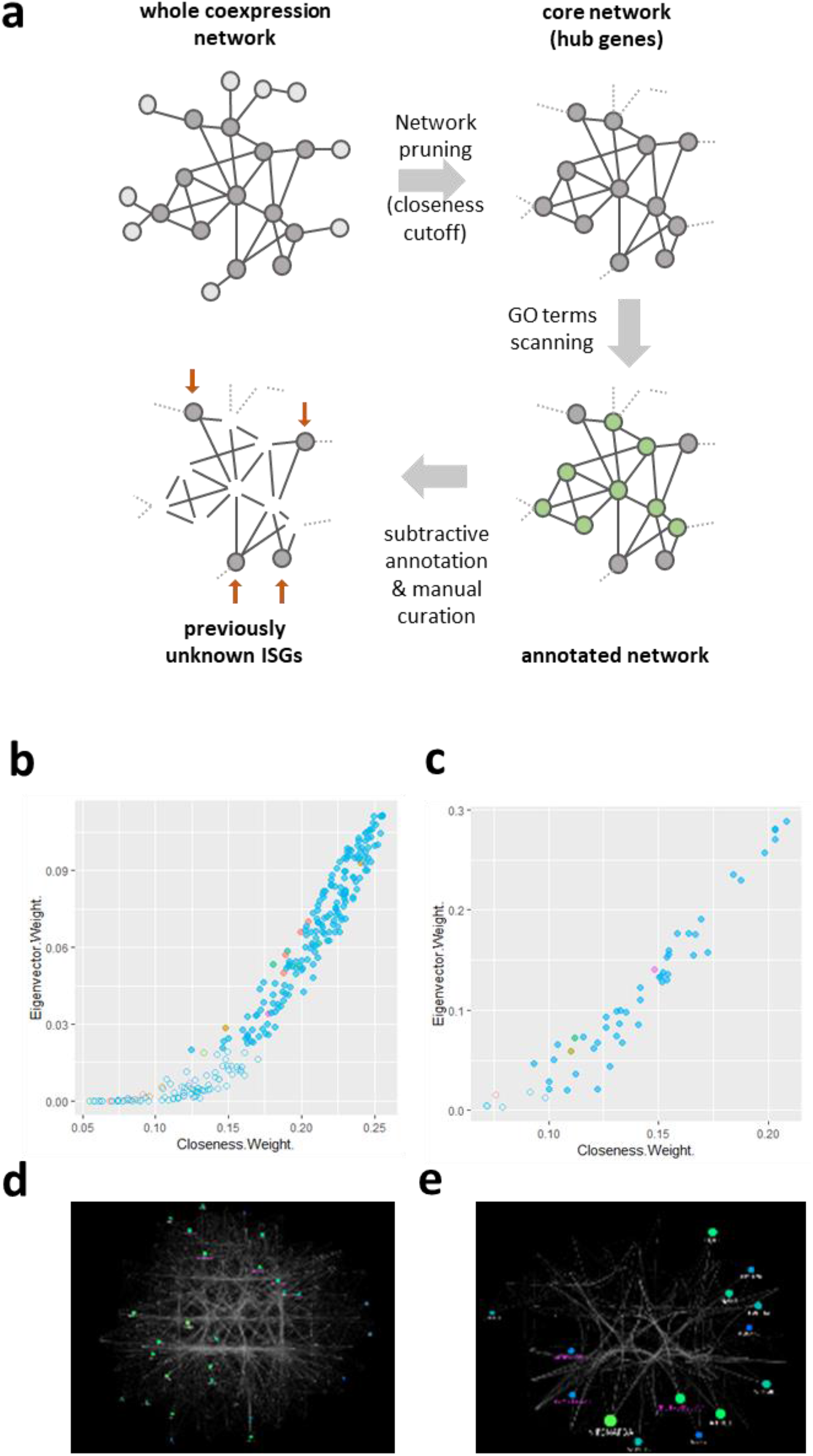
Green and Tan sub networks of putative new ISGs. **a**. The method for extraction of putative, previously unknown IFN sensitive genes, using network pruning by centrality metrics cutoffs and subtractive gene ontology annotations. **b**. Eigenvectors and closeness of genes in the Green module. These two metrics are linearly correlated above a certain threshold: higher eigenvector values correspond to higher closeness and identify hub genes. Nodes with eigenvector < 0.02 are defined as peripheral and depicted as transparent. **c**. Eigenvectors and closeness of genes in the Tan module. **d**. Cytoscape-generated sub-network for the Green module. Gene ontology annotation of surviving nodes after subtractive gene ontology annotation are displayed, while nodes (genes) associated to immune-related biological processes are not, allowing focus on non-peripheral novel players. Eigenvector is proportional to node size, closeness to color gradient. **e**. Similar cytoscape-generated sub-network for the Tan module.

We used the result of the previously performed gene ontology analysis and specifically took into account the enrichment terms related to innate immunity response. We then tagged as known players all nodes associated to at least one term among the following: innate immune and inflammatory response; type I interferon production; defense response to virus and viral process; antigen processing and presentation of peptide antigen via MHC class I; I-kappaB kinase/NF-kappaB signalling.

These nodes can be considered as known players: we then subtracted them and the remaining, un-annotated nodes were selected as potential novel players not yet characterized as related to innate immunity. Moreover, to be more stringent we also complemented this analysis with a manual curated literature search, so that only genes not previously reported as IFN-related were included in our final list. Adopting this approach, we identified 26 putative new and relevant mediators of interferon antiviral activity in the green network. The same procedure was then applied to the Tan network: topological network analysis was performed on a sub-network of 54 nodes and 601 edges; 48 genes were selected as non-peripheral and subjected to ontology annotation and manual literature search. Finally, 14 protein-coding and non-coding were selected as potential new players. The resulting combined list of putative novel mediators of interferon antiviral activity is made of 40 genes from the two selected gene modules, among which 13 non-coding RNAs (**Table 2** and “novel” classification in **Supplementary File 3**).

**Table 2.** Novel IFN sensitive genes. The 40 novel putative IFN sensitive genes (ISGs) annotated with the source module they were extracted from (Green or Tan modules), gene names, Ensembl gene IDs, biotype and description as retrieved from Ensembl Biomart.

| Source module | Gene name | Ensembl Gene ID | Biotype | Description (Biomart) |
| --- | --- | --- | --- | --- |
| Green | SIDT1 | ENSG00000072858 | protein_coding | SID1 transmembrane family member 1 |
| Green | CYTH1 | ENSG00000108669 | protein_coding | cytohesin 1 |
| Green | BLZF1 | ENSG00000117475 | protein_coding | basic leucine zipper nuclear factor 1 |
| Green | DNPEP | ENSG00000123992 | protein_coding | aspartyl aminopeptidase |
| Green | RARRES3 | ENSG00000133321 | protein_coding | phospholipase A and acyltransferase 4 |
| Green | MOB3C | ENSG00000142961 | protein_coding | MOB kinase activator 3C |
| Green | NME7 | ENSG00000143156 | protein_coding | NME/NM23 family member 7 |
| Green | GRIP2 | ENSG00000144596 | protein_coding | glutamate receptor interacting protein 2 |
| Green | GPR158 | ENSG00000151025 | protein_coding | G protein-coupled receptor 158 |
| Green | MFSD12 | ENSG00000161091 | protein_coding | major facilitator superfamily domain containing 12 |
| Green | KCNT2 | ENSG00000162687 | protein_coding | potassium sodium-activated channel subfamily T member 2 |
| Green | PLEKHA7 | ENSG00000166689 | protein_coding | pleckstrin homology domain containing A7 |
| Green | GRINA | ENSG00000178719 | protein_coding | glutamate ionotropic receptor NMDA type subunit associated protein 1 |
| Green | ZNF107 | ENSG00000196247 | protein_coding | zinc finger protein 107 |
| Green | LAP3P2 | ENSG00000213500 | processed_pseudogene | leucine aminopeptidase 3 pseudogene 2 |
| Green | SMTNL1 | ENSG00000214872 | protein_coding | smoothelin like 1 |
| Green | AC009948.5 | ENSG00000223960 | antisense | cholesterol induced regulator of metabolism RNA |
| Green | PSME2P2 | ENSG00000225131 | processed_pseudogene | proteasome activator subunit 2 pseudogene 2 |
| Green | AC009950.2 | ENSG00000225963 | antisense | novel transcript, antisense to SP110 |
| Green | AP001610.5 | ENSG00000228318 | antisense | novel transcript |
| Green | PNPT1P1 | ENSG00000229241 | processed_pseudogene | polyribonucleotide nucleotidyltransferase 1 pseudogene 1 |
| Green | PATL2 | ENSG00000229474 | protein_coding | PAT1 homolog 2 |
| Green | NAMPTL | ENSG00000229644 | processed_pseudogene | nicotinamide phosphoribosyltransferase pseudogene 1 |
| Green | HLA-K | ENSG00000230795 | unprocessed_pseudogene | major histocompatibility complex, class I, K (pseudogene) |
| Green | RP11-274E7.2 | ENSG00000238000 | processed_pseudogene | proteasome activator subunit 2 pseudogene 1 |
| Green | RP11-54O7.17 | ENSG00000272512 | lincRNA | novel transcript |
| Tan | STOML1 | ENSG00000067221 | protein_coding | stomatin like 1 |
| Tan | NANS | ENSG00000095380 | protein_coding | N-acetylneuraminate synthase |
| Tan | ATP10B | ENSG00000118322 | protein_coding | ATPase phospholipid transporting 10B (putative) |
| Tan | PRRG4 | ENSG00000135378 | protein_coding | proline rich and Gla domain 4 |
| Tan | NIPSNAP3A | ENSG00000136783 | protein_coding | nipsnap homolog 3A |
| Tan | ATHL1 | ENSG00000142102 | protein_coding | protein-glucosylgalactosylhydroxylysine glucosidase |
| Tan | GPR128 | ENSG00000144820 | protein_coding | adhesion G protein-coupled receptor G7 |
| Tan | TMCO4 | ENSG00000162542 | protein_coding | transmembrane and coiled-coil domains 4 |
| Tan | AGBL2 | ENSG00000165923 | protein_coding | ATP/GTP binding protein like 2 |
| Tan | PLEKHN1 | ENSG00000187583 | protein_coding | pleckstrin homology domain containing N1 |
| Tan | AC009299.3 | ENSG00000227403 | lincRNA | long intergenic non-protein coding RNA 1806 |
| Tan | RNF144A-AS1 | ENSG00000228203 | processed_transcript | glycosaminoglycan regulatory associated long non-coding RNA |
| Tan | RP11-395B7.7 | ENSG00000260336 | Sense_overlapping | *na, deprecated id* |
| Tan | CADPS2 | ENSG00000081803 | protein_coding | calcium dependent secretion activator 2 |

### The newly selected ISGs are enriched in IFN perturbed or HCV infected datasets

In order to validate our new IFN-stimulated signature in datasets from independent works we searched the literature and selected published datasets obtained on human samples treated with IFN. We selected and downloaded gene expression datasets of four works (Bolen *et al*, 2014; Carnero *et* al, 2014; Blackham *et al*, 2010; Grünvogel *et al*, 2015) (**Supplementary Table 1**) and for each work we obtained the ranked gene lists according to the differential expression metrics used (mainly log fold change) to compare treatments versus control samples. We then verified if our newly selected ISGs were positioned at the extremes of the gene expression datasets (i.e. if they were significantly over-represented among down or up regulated genes). To do so we used the 40 unique putative mediators of interferon antiviral activity identified by our analysis as elements of three custom gene sets (Green, Tan and the combination of the two) to scan all prepared ranked datasets with the GSEA algorithm (Subramanian *et al*, 2005). Strong upregulation of the combined geneset (genes from both the green and tan modules) was indeed observed in data obtained after induction of different interferons (**Figure 5a**), even if the genes directly responsible for the observed enrichment were not always the same with a recurrent subset of 7 genes (**Figure 5b**) among the annotated genes in the signature (DNPEP, REC8, GRINA, CYTH1, PCK2, NME7, SIDT1). With minimal differences due to time and method of induction, our newly selected gene signature was always detected among the up-regulated genes after induction or interferon related perturbation: in all analyzed published datasets taken into account the custom Green, Tan and total genesets showed a significant enrichment in most cases, with the Tan geneset being the weakest with a p-Value below the cutoff in three out of four datasets and the combined geneset just below the cutoff in GSE68927 data (**Figure 5c**). Individual GSEA enrichment plots are shown in **Supplementary Figures 8-13**. These validation analyses confirmed that integrating WGCNA analysis with subtractive gene ontology annotation was an effective way of finding mediators of interferon antiviral activity.

**Figure 5.**
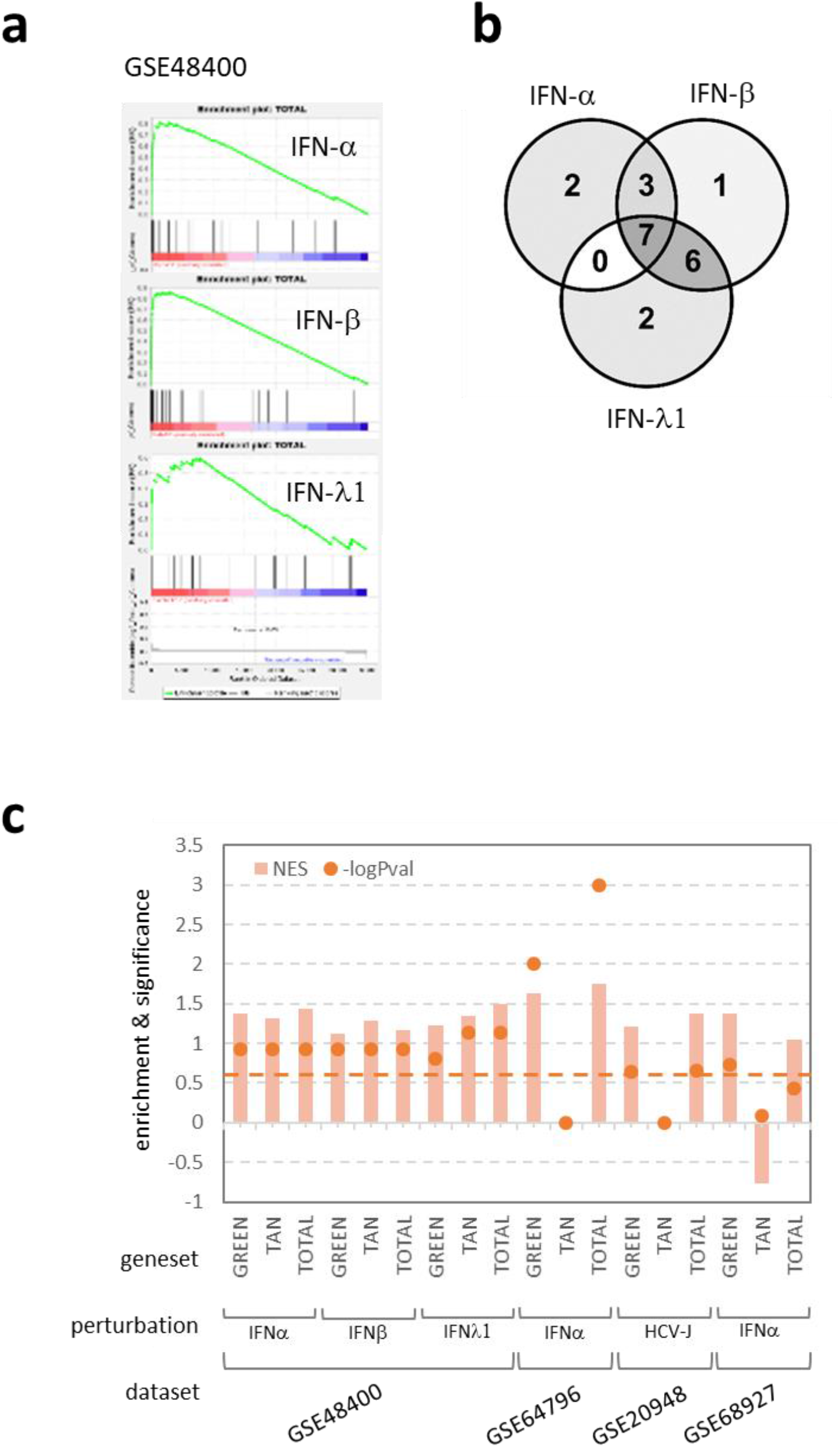
New IFN-stimulated signature is confirmed as ISGs with GSEA. **a**. GSEA running sums showing the enrichment of custom geneset obtained by the combination of the Green and Tan modules in the Boolen et al 2014 datasets (GSE48400) after 12 hours of induction with IFN alpha, beta and lambda1. **b**. Venn diagram of overlapping genes belonging to the leading edges of the three GSEA analyses in the previous panel. Seven genes (DNPEP, REC8, GRINA, CYTH1, PCK2, NME7, SIDT1) have been consistently found upregulated after different IFNs induction. **c**. Normalized enrichment scores (NES) and relative GSEA p-Values for all sets of performed GSEAs (for datasets with more time points the 12h point was selected and plotted). Dashed horizontal line is the standard GSEA significant cutoff: enrichments whose p-Value is above the dashed line are thus significant. Green, Tan and Total refer to genesets from Green module, Tan module and the combination of the two, respectively.

## Discussion

Here we report a systems biology approach which leverages the possibility to integrate datasets at different levels: transcription, membership to co-expression modules, correlation with phenotypic traits, knowledge organized as bio-ontologies, transcription factors motifs prediction and networks centrality metrics. These aspects were coherently integrated in our workflow with the aim to identify novel molecular drivers of IFN response.

Gene expression is often victim of confounding effects and it is not always possible to pinpoint all the subtle regulations and perturbations by simple, cut-off dependent, differential expression (DE) methods. A multivariate approach to gene selection allows to dive deeper into hidden relationships between molecular actors in a complex system.

By applying such an approach, we were able to find a group of genes that are specifically associated to the continuous IFN mediated modulation that were not previously described as such, and that were, at least partly, undetected in the conventional differential expression approach.

When using differential expression methods, the user has to set cut-offs and findings are necessarily dependent upon those: the more stringent the cut-offs the fewer the results. This can be harmful in the selection process favoring the more extreme perturbations over the subtler ones, thus yielding a tip-of-the-iceberg effect. Using WGCNA and the independent external measure of the IFN sensitive genes (a measure acting as a proxy to the phenotype under investigation), almost all differentially expressed genes were also selected, plus many more, without the need of setting the hard cut-offs of canonical differential expression analyses. Almost all (180 out of 185) differentially expressed genes shared by the analyzed contrasts belong to one of the two identified gene modules, while only 44% (180 out of 411) of Tan or Green modules genes are also selected as differentially expressed by canonical analysis. This means that 56% of modules genes (231 genes) we found are not selected as DE genes with the used cut-offs (0.05 FDR and 2 fold changes). In a less strict way, taking into account the sum (without repetition) of all DE genes, i.e. the genes differentially expressed in all six (2 cell lines and 3 IFN types) analyzed contrasts, the fraction of modules genes not selected as DE genes drops at 19%, yet maintaining a relevant number of genes (111) exclusively selected by the multivariate approach.

Differential expression is *per se* limited to pairwise comparisons (i.e. treated versus untreated), and each result is valid in the context of the considered pair. Taking into account only the differentially expressed genes in all comparisons under study, as it was done for the 185 perturbed in all contrasts analyzed, underestimates the biological complexity of a given set of comparisons, de facto flattening the differences. DE analysis remains the election method to select culprits in carefully designed and biologically strong contrasts, but it could be not the right choice when analyzing complex designs and looking for regulatory relationships. The fact that all genes are tested for differential expression one by one with completely independent tests is even more underestimated, since this approach once again does not take into account the natural interdependence of genes collaborating in biochemical pathways.

On the contrary, topology (centrality) metrics are a way of putting into play the relationships among genes, thus the approach is much more prone to let emerge the cooperative role of genes more than the antagonistic phenotypes. Among the genes selected in this way there could be genes that would have been pruned by an arbitrary differential expression cut-off, thus a network driven systems biology approach is more sensitive and more fit for regulatory network discovery.

For this reason, we took the advantage of having two co-expression modules well associated with the phenotype of interest to leverage them to uncover possible enrichment of transcriptional factor motifs. The findings of STAT1 and IRF9 motifs enrichment also in genes not previously described as IFN related strengthen their involvement in this process and internally validate our findings. Both IFN-α and IFN-λ induce phosphorylation of STAT1 inducing, even if with different kinetics, the expression of similar sets of ISGs, with IFN-α antiviral action being more potent than that of IFN-λ, and STAT1-mediated differential inhibition of HCV seems to be essential for IFN-λ but not for IFN-α (Yamauchi *et al*, 2016). We indeed observed an overall duality between IFN-α/β and IFN-λ gene expression patterns, but the multivariate approach that considers expression patterns across the whole samples set should act independently from such differences. The fact that motifs for IRF9 and STAT1 transcription factors are evenly distributed across the green module genes (**Supplementary Figure 5a**) could account for the above duality in order to conserve the possibility to adapt to environmental situations responding to both classes of treatment. A similar distribution is not observed for RELA and OVOL-1 motifs across genes in the Tan module (**Supplementary Figure 5b**); to this extent, the Tan module is much more a RELA related than an OVOL-1 related regulon. In addition, we also identified lncRNA specifically modulated by Type I and Type III IFN treatment. Several gene expression studies have shown that IFN profoundly changes the lncRNA cellular landscape and many lncRNAs are considered ISGs. Indeed, lncRNAs specifically regulate IFN response (Suarez *et al*, 2020). Interestingly, we identified miR-146a as an upregulated lncRNA after IFN-α treatment. miR-146a plays an important role in modulating type I IFN response and inflammation by targeting TRAF6 (Wu *et al*, 2013). Increase in miR-146a levels suppresses type I interferon and facilitates infection of several viruses like EV71, dengue virus and HIV (Fu *et al*, 2017; Wu *et al*, 2013; Teng *et al*, 2019). On the other hand, inhibition of miR-146a restores type I interferon production and blocks enterovirus-induced cell death (Ho *et al*, 2014). Finally, we validated our datasets of new IFN-stimulated genes, in independent works suggesting that our method is suitable to interrogate different datasets for the discovery of new mediators of interferon pathways.

## Methods

### Cells and IFN perturbation

Huh7 and Huh7.5 cells were maintained in Dulbecco’s modified Eagle medium (DMEM) supplemented with 10% fetal bovine serum (FBS), penicillin-streptomycin and 2mM L-glutamine (Blight *et al*, 2002). Cells were treated for 8h with 10ng/ml of Interferon-α2A (cod.H6041, SIGMA), 5000 U/ml of Interferon-β1A (cod.I4151, SIGMA) or 100ng/ml of interferon-λ3 (cod.E92028Hu, Uscn Life Science Inc) prior RNA extraction and library preparation.

### RNA extraction and library preparation

RNA-Seq experiments were performed on Huh7 and Huh7.5 treated with Interferon-α2A, Interferon-β1A (cod.I4151, SIGMA) or Interferon-λ3 as described above. Mock-treated cells were used as control for each treatment. At least 3 biological replicates were analysed for each biological condition. Total RNA was extracted with mirVana miRNA isolation kit and DNAse digested (Ambion) according to manufacturer’s instruction. RNA samples were submitted to quality control by RNA integrity number evaluation. Libraries for Illumina sequencing were constructed by polyA selection with the Illumina Kit. A paired end (2×100) run was performed.

### RNAseq gene expression and differential expression analysis

After initial quality check reads were filtered to remove low quality calls by Trimmomatic (version 0.XX) (24695404) using default parameters. Processed reads were then aligned to human genome assembly GRCh38 (Ensembl release 77) with STAR mapping software (version 2.3.0e) (Dobin *et al*, 2013). HTSeq-count algorithm (version 0.6.1) (Anders *et al*, 2015) option –str = no, gene annotation release 77 from Ensembl) was employed to produce read counts at the gene level for each sample. To estimate differential expression, the matrix of gene counts produced by HTSeq was analysed by DESeq2 (Love *et al*, 2014). Differential expression analysis was performed to compare each treatment with the respective mock-treated cells (6 comparisons, 43 samples overall). Read counts were normalized by calculating size factors, while Variance Stabilizing Transformed (vst) values were employed for principal component analysis, as implemented in DESeq2.

Independent filtering procedure was then applied, setting the threshold to the 76 percentile (5.74 read counts); 15421 genes were subsequently tested for differential expression. Significantly modulated genes were selected by considering an absolute value of log2 fold change (Log2FC) higher than 1 and a false discovery rate (FDR) lower than 5%.

### Weighted Gene Co-expression network generation and module identification

Variance Stabilizing Transformed values (as calculated by DESeq2) were used as input data for network analysis. Starting from the set of 15421 genes identified by independent filtering, a gene selection strategy was applied by calculating for each gene the coefficient of variation (CV) across the 43 biological samples. A 60-percentile threshold was then imposed, thus selecting the 40% of genes showing the highest variability (highest CV values, 6168 genes). Sample average-linkage hierarchical cluster analysis was performed as a quality control step on this gene subset. A signed hybrid gene co-expression network was generated relying on the WGCNA R package (version 1.43-10). (Langfelder & Horvath, 2008) The correlation matrix was calculated based on the Pearson metrics and then transformed into an adjacency matrix by raising it to the power of β = 12, chosen on the basis of the scale-free topology criterion. Topological Overlap Measure was calculated from the adjacency matrix and the relative dissimilarity matrix was used as input for average-linkage hierarchical clustering and gene dendrogram generation. Network modules were detected as branches of the dendrogram by using the DynamicTree Cut algorithm (deepSplit=1, minimum cluster size= 40) (Langfelder *et al*, 2008). Muldi-dimensional scaling was calculated on the topological overlap dissimilarity matrix.

### Module-trait correlation

As a phenotypic trait for module-trait correlation, the transcripts of the two well-known ISGs MX1 and IFIT2 were quantified by real-time PCR in the 43 samples.

For real-time PCR, TaqMan gene expression assay (Thermo Fisher scientific) was performed according to manufacturer’s instruction. ΔCt versus the gene normalizer GAPDH and then ΔΔCt versus the mean ΔCt value in corresponding non-treated cells were calculated for each of the six treatments. Minus ΔΔCt was used for downstream analysis, so that an induction of ISGs was quantified by a high -ΔΔCt value. For each module, Pearson correlation between the module eigengene and the -ΔΔCt value was calculated. For the selected modules (Green and Tan) sample clustering and heat map illustrating the gene expression profiles were generated starting from Variance Stabilizing Transformed values.

### Sub-network generation and topological network analysis in Cytoscape

Node and edge information for the selected modules (Green and Tan) were exported from the adjacency matrix relying on the appropriate WGCNA function and imposing a threshold of 0.1 as minimum edge weight. 282 nodes and 18085 edges for the Green module and 54 nodes and 601 edges for the Tan module were therefore imported in Cytoscape for further analysis (Shannon *et al*, 2003). Topological network analysis for Green and Tan sub-networks was performed in Cytoscape relying on CytoNCA app using the “with weight” analysis (Kijima & Kijima, 1987). Peripheral genes were identified as those showing an eigenvector value lower than 0.02.

### Gene Ontology Enrichment Analysis

Gene ontology enrichment analysis for the Biological Process domain was performed on the 282 genes belonging to the green sub-network, using the list of 6168 genes selected for network generation as a custom reference set. Analysis was performed by TopGO (version 2.18.0) (Alexa *et al*, 2006) relying on Fisher test and Weight01 method to take into account ontology hierarchy; node size was set at 20. A 0.01 p-value cut off was applied to select significantly enriched GO terms. The same procedure was performed on the 54 nodes of the Tan sub-network. Overall screening gene ontology analysis contemplating more databases (ontologies, pathways, motifs, proteins and phenotypes) on cell-line specific differentially expressed genes were also performed using g:Profiler (Raudvere *et al*, 2019).

### Regulatory motif analysis

Cytoscape app (version 1.3) (Janky *et al*, 2014) was used to identify enriched regulatory regions in Green and Tan sub-networks. Analysis was performed on 5000 bp upstream the TSS, using 10K motif collection and retrieving motifs with a Normalized Enrichment score higher than 5; maximum FDR on motif similarity was set at 0.005. Circular plots for result visualization were produced in R by the circlize package (version 0.2.4) (Gu *et al*, 2014).

### Functional annotation

Starting from the results of the gene ontology analysis, the terms found as enriched considering a threshold of p-Value < 0.01 were then classified as related or unrelated to innate immunity phenomena. For the identification of genes related to innate immune response, descendent terms of the following categories were considered: GO:0045087 - innate immune response; GO:0006954 - inflammatory response; GO:0032606 - type I interferon production; GO:0051607 - defense response to virus; GO:0016032 - viral process; GO:0002474 - antigen processing and presentation of peptide antigen via MHC class I; GO:0007249 - I-kappaB kinase/NF-kappaB signalling. Genes associated with at least one of these groups of terms were therefore defined as ‘known players’, while those not associated with any were defined “IFN- unrelated”.

### Gene Set Enrichment Analyses

Gene set enrichment analysis (GSEA) was used to validate the selected gene signature of putative ISGs as actually over represented in other published datasets of IFN induction studies. GSEA was performed as described in the original paper (Subramanian *et al*, 2005) using the Broad Institute official java implementation, with custom genesets. Normalized gene expression data of the selected literature were downloaded from the GEO Omnibus repository and used as GSEA input datasets; ranked gene lists of differential expression analyses performed were obtained using Signal to Noise or Log2 Ratios metrics. Custom genesets of 28, 14 and 42 genes were built using the candidate ISGs from the Green module, Tan module and both combined, respectively, after harmonization of gene names annotation across the gene expression platforms used in the four different works. GSEA was then performed with default parameters and cutoffs. GSEA running plots depicts enrichment scores (ES), while in the summary plot of all GSEA results were used normalized enrichment scores (NES) with cognate p-Values.

## Authors Contributions

RDF conceived the study. CC, LM, RDF, and RLR designed experiments and analytical workflows. LD and VB performed experiments. RJPB generated RNAseq raw counts data. CC wrote code, processed and analyzed RNAseq, network and functional enrichment data. RLR analyzed functional enrichment data, GSEA validations and integrated whole data. CC, LM, LD, VB, RLR and RDF interpreted the result. CC, LM, RLR wrote the manuscript. CC and RLR made figures. LM contributed virological expertise and finalized the manuscript. RLR coordinated the efforts and RDF supervised them.

## Acknowledgements

We would like to thank Valeria Zanoni for technical laboratory support.

