## Supplemental File 1 for "Novel antiviral interferon sensitive genes unveiled by correlation-driven gene selection and integrated systems biology approaches"

| cell_line | interferon | ensembl_gene_id | gene_symbol | gene_biotype |
|---|---|---|---|---|
| Huh7 | alfa | ENSG00000225886 | | antisense |
| Huh7 | alfa | ENSG00000242539 | | antisense |
| Huh7 | alfa | ENSG00000225963 | | antisense |
| Huh7 | alfa | ENSG00000223960 | | antisense |
| Huh7 | alfa | ENSG00000228318 | | antisense |
| Huh7 | alfa | ENSG00000263069 | | antisense |
| Huh7 | alfa | ENSG00000215068 | | antisense |
| Huh7 | alfa | ENSG00000225377 | NRSN2-AS1 | antisense |
| Huh7 | alfa | ENSG00000255031 | | antisense |
| Huh7 | alfa | ENSG00000225075 | | antisense |
| Huh7 | alfa | ENSG00000280007 | | antisense |
| Huh7 | alfa | ENSG00000232498 | | antisense |
| Huh7 | alfa | ENSG00000267270 | PARD6G-AS1 | antisense |
| Huh7 | alfa | ENSG00000229873 | OGFR-AS1 | antisense |
| Huh7 | alfa | ENSG00000269728 | | lincRNA |
| Huh7 | alfa | ENSG00000269640 | | lincRNA |
| Huh7 | alfa | ENSG00000272512 | | lincRNA |
| Huh7 | alfa | ENSG00000245571 | | lincRNA |
| Huh7 | alfa | ENSG00000260302 | | lincRNA |
| Huh7 | alfa | ENSG00000253522 | MIR146A | lincRNA |
| Huh7 | alfa | ENSG00000262454 | | lincRNA |
| Huh7 | alfa | ENSG00000260750 | | lincRNA |
| Huh7 | alfa | ENSG00000258733 | | lincRNA |
| Huh7 | alfa | ENSG00000251637 | | lincRNA |
| Huh7 | alfa | ENSG00000228222 | | lincRNA |
| Huh7 | alfa | ENSG00000243650 | RN7SL834P | misc_RNA |
| Huh7 | alfa | ENSG00000229644 | NAMPTL | processed_pseudogene |
| Huh7 | alfa | ENSG00000238000 | | processed_pseudogene |
| Huh7 | alfa | ENSG00000225131 | PSME2P2 | processed_pseudogene |
| Huh7 | alfa | ENSG00000229241 | PNPT1P1 | processed_pseudogene |
| Huh7 | alfa | ENSG00000213500 | LAP3P2 | processed_pseudogene |
| Huh7 | alfa | ENSG00000271550 | BNIP3P11 | processed_pseudogene |
| Huh7 | alfa | ENSG00000241556 | | processed_pseudogene |
| Huh7 | alfa | ENSG00000242262 | | processed_pseudogene |
| Huh7 | alfa | ENSG00000258441 | LINC00641 | processed_transcript |
| Huh7 | alfa | ENSG00000160710 | ADAR | protein_coding |
| Huh7 | alfa | ENSG00000138035 | PNPT1 | protein_coding |
| Huh7 | alfa | ENSG00000115415 | STAT1 | protein_coding |
| Huh7 | alfa | ENSG00000101347 | SAMHD1 | protein_coding |
| Huh7 | alfa | ENSG00000173821 | RNF213 | protein_coding |
| Huh7 | alfa | ENSG00000055332 | EIF2AK2 | protein_coding |
| Huh7 | alfa | ENSG00000170581 | STAT2 | protein_coding |
| Huh7 | alfa | ENSG00000121060 | TRIM25 | protein_coding |
| Huh7 | alfa | ENSG00000188313 | PLSCR1 | protein_coding |
| Huh7 | alfa | ENSG00000163840 | DTX3L | protein_coding |
| Huh7 | alfa | ENSG00000184979 | USP18 | protein_coding |
| Huh7 | alfa | ENSG00000156587 | UBE2L6 | protein_coding |
| Huh7 | alfa | ENSG00000138496 | PARP9 | protein_coding |
| Huh7 | alfa | ENSG00000181381 | DDX60L | protein_coding |
| Huh7 | alfa | ENSG00000173193 | PARP14 | protein_coding |
| Huh7 | alfa | ENSG00000187608 | ISG15 | protein_coding |
| Huh7 | alfa | ENSG00000221963 | APOL6 | protein_coding |
| Huh7 | alfa | ENSG00000111331 | OAS3 | protein_coding |
| Huh7 | alfa | ENSG00000157601 | MX1 | protein_coding |
| Huh7 | alfa | ENSG00000185745 | IFIT1 | protein_coding |
| Huh7 | alfa | ENSG00000115267 | IFIH1 | protein_coding |
| Huh7 | alfa | ENSG00000067066 | SP100 | protein_coding |
| Huh7 | alfa | ENSG00000107201 | DDX58 | protein_coding |
| Huh7 | alfa | ENSG00000059378 | PARP12 | protein_coding |
| Huh7 | alfa | ENSG00000172936 | MYD88 | protein_coding |
| Huh7 | alfa | ENSG00000105939 | ZC3HAV1 | protein_coding |

| | | | | |
|---|---|---|---|---|
| Huh7 | alfa | ENSG00000178685 | PARP10 | protein_coding |
| Huh7 | alfa | ENSG00000168394 | TAP1 | protein_coding |
| Huh7 | alfa | ENSG00000119917 | IFIT3 | protein_coding |
| Huh7 | alfa | ENSG00000196116 | TDRD7 | protein_coding |
| Huh7 | alfa | ENSG00000155629 | PIK3AP1 | protein_coding |
| Huh7 | alfa | ENSG00000111335 | OAS2 | protein_coding |
| Huh7 | alfa | ENSG00000002549 | LAP3 | protein_coding |
| Huh7 | alfa | ENSG00000133106 | EPSTI1 | protein_coding |
| Huh7 | alfa | ENSG00000135899 | SP110 | protein_coding |
| Huh7 | alfa | ENSG00000166710 | B2M | protein_coding |
| Huh7 | alfa | ENSG00000126709 | IFI6 | protein_coding |
| Huh7 | alfa | ENSG00000152778 | IFIT5 | protein_coding |
| Huh7 | alfa | ENSG00000165806 | CASP7 | protein_coding |
| Huh7 | alfa | ENSG00000108679 | LGALS3BP | protein_coding |
| Huh7 | alfa | ENSG00000089127 | OAS1 | protein_coding |
| Huh7 | alfa | ENSG00000124201 | ZNFX1 | protein_coding |
| Huh7 | alfa | ENSG00000137628 | DDX60 | protein_coding |
| Huh7 | alfa | ENSG00000117228 | GBP1 | protein_coding |
| Huh7 | alfa | ENSG00000138642 | HERC6 | protein_coding |
| Huh7 | alfa | ENSG00000134321 | RSAD2 | protein_coding |
| Huh7 | alfa | ENSG00000142089 | IFITM3 | protein_coding |
| Huh7 | alfa | ENSG00000173786 | CNP | protein_coding |
| Huh7 | alfa | ENSG00000078081 | LAMP3 | protein_coding |
| Huh7 | alfa | ENSG00000169245 | CXCL10 | protein_coding |
| Huh7 | alfa | ENSG00000123609 | NMI | protein_coding |
| Huh7 | alfa | ENSG00000204592 | HLA-E | protein_coding |
| Huh7 | alfa | ENSG00000168016 | TRANK1 | protein_coding |
| Huh7 | alfa | ENSG00000134326 | CMPK2 | protein_coding |
| Huh7 | alfa | ENSG00000119922 | IFIT2 | protein_coding |
| Huh7 | alfa | ENSG00000204525 | HLA-C | protein_coding |
| Huh7 | alfa | ENSG00000130303 | BST2 | protein_coding |
| Huh7 | alfa | ENSG00000205413 | SAMD9 | protein_coding |
| Huh7 | alfa | ENSG00000110446 | SLC15A3 | protein_coding |
| Huh7 | alfa | ENSG00000140853 | NLRC5 | protein_coding |
| Huh7 | alfa | ENSG00000135114 | OASL | protein_coding |
| Huh7 | alfa | ENSG00000128335 | APOL2 | protein_coding |
| Huh7 | alfa | ENSG00000137959 | IFI44L | protein_coding |
| Huh7 | alfa | ENSG00000130589 | HELZ2 | protein_coding |
| Huh7 | alfa | ENSG00000132274 | TRIM22 | protein_coding |
| Huh7 | alfa | ENSG00000138646 | HERC5 | protein_coding |
| Huh7 | alfa | ENSG00000105287 | PRKD2 | protein_coding |
| Huh7 | alfa | ENSG00000231925 | TAPBP | protein_coding |
| Huh7 | alfa | ENSG00000173530 | TNFRSF10D | protein_coding |
| Huh7 | alfa | ENSG00000132256 | TRIM5 | protein_coding |
| Huh7 | alfa | ENSG00000185880 | TRIM69 | protein_coding |
| Huh7 | alfa | ENSG00000100342 | APOL1 | protein_coding |
| Huh7 | alfa | ENSG00000164307 | ERAP1 | protein_coding |
| Huh7 | alfa | ENSG00000140464 | PML | protein_coding |
| Huh7 | alfa | ENSG00000111912 | NCOA7 | protein_coding |
| Huh7 | alfa | ENSG00000121858 | TNFSF10 | protein_coding |
| Huh7 | alfa | ENSG00000108669 | CYTH1 | protein_coding |
| Huh7 | alfa | ENSG00000013374 | NUB1 | protein_coding |
| Huh7 | alfa | ENSG00000068079 | IFI35 | protein_coding |
| Huh7 | alfa | ENSG00000182179 | UBA7 | protein_coding |
| Huh7 | alfa | ENSG00000169429 | CXCL8 | protein_coding |
| Huh7 | alfa | ENSG00000130813 | C19orf66 | protein_coding |
| Huh7 | alfa | ENSG00000204267 | TAP2 | protein_coding |
| Huh7 | alfa | ENSG00000204264 | PSMB8 | protein_coding |
| Huh7 | alfa | ENSG00000152689 | RASGRP3 | protein_coding |
| Huh7 | alfa | ENSG00000132109 | TRIM21 | protein_coding |
| Huh7 | alfa | ENSG00000163644 | PPM1K | protein_coding |
| Huh7 | alfa | ENSG00000123240 | OPTN | protein_coding |

| | | | | |
|---|---|---|---|---|
| Huh7 | alfa | ENSG00000000971 | CFH | protein_coding |
| Huh7 | alfa | ENSG00000132530 | XAF1 | protein_coding |
| Huh7 | alfa | ENSG00000240065 | PSMB9 | protein_coding |
| Huh7 | alfa | ENSG00000079385 | CEACAM1 | protein_coding |
| Huh7 | alfa | ENSG00000185404 | SP140L | protein_coding |
| Huh7 | alfa | ENSG00000117226 | GBP3 | protein_coding |
| Huh7 | alfa | ENSG00000116663 | FBXO6 | protein_coding |
| Huh7 | alfa | ENSG00000026950 | BTN3A1 | protein_coding |
| Huh7 | alfa | ENSG00000162654 | GBP4 | protein_coding |
| Huh7 | alfa | ENSG00000105559 | PLEKHA4 | protein_coding |
| Huh7 | alfa | ENSG00000086061 | DNAJA1 | protein_coding |
| Huh7 | alfa | ENSG00000168062 | BATF2 | protein_coding |
| Huh7 | alfa | ENSG00000172183 | ISG20 | protein_coding |
| Huh7 | alfa | ENSG00000123992 | DNPEP | protein_coding |
| Huh7 | alfa | ENSG00000010030 | ETV7 | protein_coding |
| Huh7 | alfa | ENSG00000164342 | TLR3 | protein_coding |
| Huh7 | alfa | ENSG00000168404 | MLKL | protein_coding |
| Huh7 | alfa | ENSG00000118503 | TNFAIP3 | protein_coding |
| Huh7 | alfa | ENSG00000177409 | SAMD9L | protein_coding |
| Huh7 | alfa | ENSG00000168297 | PXK | protein_coding |
| Huh7 | alfa | ENSG00000173702 | MUC13 | protein_coding |
| Huh7 | alfa | ENSG00000182326 | C1S | protein_coding |
| Huh7 | alfa | ENSG00000137965 | IFI44 | protein_coding |
| Huh7 | alfa | ENSG00000114450 | GNB4 | protein_coding |
| Huh7 | alfa | ENSG00000105835 | NAMPT | protein_coding |
| Huh7 | alfa | ENSG00000092010 | PSME1 | protein_coding |
| Huh7 | alfa | ENSG00000120539 | MASTL | protein_coding |
| Huh7 | alfa | ENSG00000117475 | BLZF1 | protein_coding |
| Huh7 | alfa | ENSG00000234745 | HLA-B | protein_coding |
| Huh7 | alfa | ENSG00000106785 | TRIM14 | protein_coding |
| Huh7 | alfa | ENSG00000186470 | BTN3A2 | protein_coding |
| Huh7 | alfa | ENSG00000122643 | NT5C3A | protein_coding |
| Huh7 | alfa | ENSG00000100911 | PSME2 | protein_coding |
| Huh7 | alfa | ENSG00000148468 | FAM171A1 | protein_coding |
| Huh7 | alfa | ENSG00000151025 | GPR158 | protein_coding |
| Huh7 | alfa | ENSG00000146859 | TMEM140 | protein_coding |
| Huh7 | alfa | ENSG00000165949 | IFI27 | protein_coding |
| Huh7 | alfa | ENSG00000137200 | CMTR1 | protein_coding |
| Huh7 | alfa | ENSG00000064012 | CASP8 | protein_coding |
| Huh7 | alfa | ENSG00000145623 | OSMR | protein_coding |
| Huh7 | alfa | ENSG00000113441 | LNPEP | protein_coding |
| Huh7 | alfa | ENSG00000166689 | PLEKHA7 | protein_coding |
| Huh7 | alfa | ENSG00000178719 | GRINA | protein_coding |
| Huh7 | alfa | ENSG00000164308 | ERAP2 | protein_coding |
| Huh7 | alfa | ENSG00000135148 | TRAFD1 | protein_coding |
| Huh7 | alfa | ENSG00000169248 | CXCL11 | protein_coding |
| Huh7 | alfa | ENSG00000114127 | XRN1 | protein_coding |
| Huh7 | alfa | ENSG00000163545 | NUAK2 | protein_coding |
| Huh7 | alfa | ENSG00000078018 | MAP2 | protein_coding |
| Huh7 | alfa | ENSG00000164211 | STARD4 | protein_coding |
| Huh7 | alfa | ENSG00000164761 | TNFRSF11B | protein_coding |
| Huh7 | alfa | ENSG00000133321 | RARRES3 | protein_coding |
| Huh7 | alfa | ENSG00000115009 | CCL20 | protein_coding |
| Huh7 | alfa | ENSG00000023445 | BIRC3 | protein_coding |
| Huh7 | alfa | ENSG00000184898 | RBM43 | protein_coding |
| Huh7 | alfa | ENSG00000213928 | IRF9 | protein_coding |
| Huh7 | alfa | ENSG00000125347 | IRF1 | protein_coding |
| Huh7 | alfa | ENSG00000120217 | CD274 | protein_coding |
| Huh7 | alfa | ENSG00000196247 | ZNF107 | protein_coding |
| Huh7 | alfa | ENSG00000180628 | PCGF5 | protein_coding |
| Huh7 | alfa | ENSG00000155363 | MOV10 | protein_coding |
| Huh7 | alfa | ENSG00000185885 | IFITM1 | protein_coding |

| | | | | |
|---|---|---|---|---|
| Huh7 | alfa | ENSG00000164136 | IL15 | protein_coding |
| Huh7 | alfa | ENSG00000156966 | B3GNT7 | protein_coding |
| Huh7 | alfa | ENSG00000079215 | SLC1A3 | protein_coding |
| Huh7 | alfa | ENSG00000168310 | IRF2 | protein_coding |
| Huh7 | alfa | ENSG00000136147 | PHF11 | protein_coding |
| Huh7 | alfa | ENSG00000134716 | CYP2J2 | protein_coding |
| Huh7 | alfa | ENSG00000100918 | REC8 | protein_coding |
| Huh7 | alfa | ENSG00000108771 | DHX58 | protein_coding |
| Huh7 | alfa | ENSG00000134470 | IL15RA | protein_coding |
| Huh7 | alfa | ENSG00000120549 | KIAA1217 | protein_coding |
| Huh7 | alfa | ENSG00000198959 | TGM2 | protein_coding |
| Huh7 | alfa | ENSG00000196954 | CASP4 | protein_coding |
| Huh7 | alfa | ENSG00000060491 | OGFR | protein_coding |
| Huh7 | alfa | ENSG00000142677 | IL22RA1 | protein_coding |
| Huh7 | alfa | ENSG00000162687 | KCNT2 | protein_coding |
| Huh7 | alfa | ENSG00000142961 | MOB3C | protein_coding |
| Huh7 | alfa | ENSG00000234127 | TRIM26 | protein_coding |
| Huh7 | alfa | ENSG00000173930 | SLCO4C1 | protein_coding |
| Huh7 | alfa | ENSG00000149131 | SERPING1 | protein_coding |
| Huh7 | alfa | ENSG00000155287 | SLC25A28 | protein_coding |
| Huh7 | alfa | ENSG00000119487 | MAPKAP1 | protein_coding |
| Huh7 | alfa | ENSG00000139083 | ETV6 | protein_coding |
| Huh7 | alfa | ENSG00000159403 | C1R | protein_coding |
| Huh7 | alfa | ENSG00000104518 | GSDMD | protein_coding |
| Huh7 | alfa | ENSG00000214872 | SMTNL1 | protein_coding |
| Huh7 | alfa | ENSG00000141682 | PMAIP1 | protein_coding |
| Huh7 | alfa | ENSG00000197536 | C5orf56 | protein_coding |
| Huh7 | alfa | ENSG00000136514 | RTP4 | protein_coding |
| Huh7 | alfa | ENSG00000137752 | CASP1 | protein_coding |
| Huh7 | alfa | ENSG00000130487 | KLHDC7B | protein_coding |
| Huh7 | alfa | ENSG00000144596 | GRIP2 | protein_coding |
| Huh7 | alfa | ENSG00000162645 | GBP2 | protein_coding |
| Huh7 | alfa | ENSG00000128394 | APOBEC3F | protein_coding |
| Huh7 | alfa | ENSG00000177989 | ODF3B | protein_coding |
| Huh7 | alfa | ENSG00000072858 | SIDT1 | protein_coding |
| Huh7 | alfa | ENSG00000127666 | TICAM1 | protein_coding |
| Huh7 | alfa | ENSG00000161091 | MFSD12 | protein_coding |
| Huh7 | alfa | ENSG00000137842 | TMEM62 | protein_coding |
| Huh7 | alfa | ENSG00000166016 | ABTB2 | protein_coding |
| Huh7 | alfa | ENSG00000229474 | PATL2 | protein_coding |
| Huh7 | alfa | ENSG00000172123 | SLFN12 | protein_coding |
| Huh7 | alfa | ENSG00000184371 | CSF1 | protein_coding |
| Huh7 | alfa | ENSG00000166801 | FAM111A | protein_coding |
| Huh7 | alfa | ENSG00000185201 | IFITM2 | protein_coding |
| Huh7 | alfa | ENSG00000105402 | NAPA | protein_coding |
| Huh7 | alfa | ENSG00000163739 | CXCL1 | protein_coding |
| Huh7 | alfa | ENSG00000181541 | MAB21L2 | protein_coding |
| Huh7 | alfa | ENSG00000125826 | RBCK1 | protein_coding |
| Huh7 | alfa | ENSG00000169871 | TRIM56 | protein_coding |
| Huh7 | alfa | ENSG00000144852 | NR1I2 | protein_coding |
| Huh7 | alfa | ENSG00000117266 | CDK18 | protein_coding |
| Huh7 | alfa | ENSG00000151470 | C4orf33 | protein_coding |
| Huh7 | alfa | ENSG00000111801 | BTN3A3 | protein_coding |
| Huh7 | alfa | ENSG00000073605 | GSDMB | protein_coding |
| Huh7 | alfa | ENSG00000049192 | ADAMTS6 | protein_coding |
| Huh7 | alfa | ENSG00000205364 | MT1M | protein_coding |
| Huh7 | alfa | ENSG00000205220 | PSMB10 | protein_coding |
| Huh7 | alfa | ENSG00000110057 | UNC93B1 | protein_coding |
| Huh7 | alfa | ENSG00000140511 | HAPLN3 | protein_coding |
| Huh7 | alfa | ENSG00000163131 | CTSS | protein_coding |
| Huh7 | alfa | ENSG00000188613 | NANOS1 | protein_coding |
| Huh7 | alfa | ENSG00000008441 | NFIX | protein_coding |

| | | | | |
|---|---|---|---|---|
| Huh7 | alfa | ENSG00000135378 | PRRG4 | protein_coding |
| Huh7 | alfa | ENSG00000112343 | TRIM38 | protein_coding |
| Huh7 | alfa | ENSG00000157303 | SUSD3 | protein_coding |
| Huh7 | alfa | ENSG00000174808 | BTC | protein_coding |
| Huh7 | alfa | ENSG00000206503 | HLA-A | protein_coding |
| Huh7 | alfa | ENSG00000163823 | CCR1 | protein_coding |
| Huh7 | alfa | ENSG00000103056 | SMPD3 | protein_coding |
| Huh7 | alfa | ENSG00000125148 | MT2A | protein_coding |
| Huh7 | alfa | ENSG00000137767 | SQRDL | protein_coding |
| Huh7 | alfa | ENSG00000259529 | IRF9 | protein_coding |
| Huh7 | alfa | ENSG00000106302 | HYAL4 | protein_coding |
| Huh7 | alfa | ENSG00000169894 | MUC3A | protein_coding |
| Huh7 | alfa | ENSG00000168497 | SDPR | protein_coding |
| Huh7 | alfa | ENSG00000263528 | IKBKE | protein_coding |
| Huh7 | alfa | ENSG00000122042 | UBL3 | protein_coding |
| Huh7 | alfa | ENSG00000042980 | ADAM28 | protein_coding |
| Huh7 | alfa | ENSG00000141655 | TNFRSF11A | protein_coding |
| Huh7 | alfa | ENSG00000128342 | LIF | protein_coding |
| Huh7 | alfa | ENSG00000121380 | BCL2L14 | protein_coding |
| Huh7 | alfa | ENSG00000157873 | TNFRSF14 | protein_coding |
| Huh7 | alfa | ENSG00000166741 | NNMT | protein_coding |
| Huh7 | alfa | ENSG00000139192 | TAPBPL | protein_coding |
| Huh7 | alfa | ENSG00000141664 | ZCCHC2 | protein_coding |
| Huh7 | alfa | ENSG00000187583 | PLEKHN1 | protein_coding |
| Huh7 | alfa | ENSG00000087085 | ACHE | protein_coding |
| Huh7 | alfa | ENSG00000156515 | HK1 | protein_coding |
| Huh7 | alfa | ENSG00000107593 | PKD2L1 | protein_coding |
| Huh7 | alfa | ENSG00000120341 | SEC16B | protein_coding |
| Huh7 | alfa | ENSG00000204397 | CARD16 | protein_coding |
| Huh7 | alfa | ENSG00000138435 | CHRNA1 | protein_coding |
| Huh7 | alfa | ENSG00000147168 | IL2RG | protein_coding |
| Huh7 | alfa | ENSG00000248713 | | protein_coding |
| Huh7 | alfa | ENSG00000166278 | C2 | protein_coding |
| Huh7 | alfa | ENSG00000129654 | FOXJ1 | protein_coding |
| Huh7 | alfa | ENSG00000255398 | HCAR3 | protein_coding |
| Huh7 | alfa | ENSG00000183397 | C19orf71 | protein_coding |
| Huh7 | alfa | ENSG00000156500 | FAM122C | protein_coding |
| Huh7 | alfa | ENSG00000105967 | TFEC | protein_coding |
| Huh7 | alfa | ENSG00000025708 | TYMP | protein_coding |
| Huh7 | alfa | ENSG00000093134 | VNN3 | protein_coding |
| Huh7 | alfa | ENSG00000174428 | GTF2IRD2B | protein_coding |
| Huh7 | alfa | ENSG00000182782 | HCAR2 | protein_coding |
| Huh7 | alfa | ENSG00000104432 | IL7 | protein_coding |
| Huh7 | alfa | ENSG00000197408 | CYP2B6 | protein_coding |
| Huh7 | alfa | ENSG00000117834 | SLC5A9 | protein_coding |
| Huh7 | alfa | ENSG00000214226 | C17orf67 | protein_coding |
| Huh7 | alfa | ENSG00000155158 | TTC39B | protein_coding |
| Huh7 | alfa | ENSG00000108622 | ICAM2 | protein_coding |
| Huh7 | alfa | ENSG00000178597 | PSAPL1 | protein_coding |
| Huh7 | alfa | ENSG00000135374 | ELF5 | protein_coding |
| Huh7 | alfa | ENSG00000081041 | CXCL2 | protein_coding |
| Huh7 | alfa | ENSG00000167601 | AXL | protein_coding |
| Huh7 | alfa | ENSG00000185950 | IRS2 | protein_coding |
| Huh7 | alfa | ENSG00000163121 | NEURL3 | protein_coding |
| Huh7 | alfa | ENSG00000104213 | PDGFRL | protein_coding |
| Huh7 | alfa | ENSG00000177721 | ANXA2R | protein_coding |
| Huh7 | alfa | ENSG00000043355 | ZIC2 | protein_coding |
| Huh7 | alfa | ENSG00000234965 | SHISA8 | protein_coding |
| Huh7 | alfa | ENSG00000144820 | GPR128 | protein_coding |
| Huh7 | alfa | ENSG00000133943 | C14orf159 | protein_coding |
| Huh7 | alfa | ENSG00000255408 | PCDHA3 | protein_coding |
| Huh7 | alfa | ENSG00000243649 | CFB | protein_coding |

| | | | | |
|---|---|---|---|---|
| Huh7 | alfa | ENSG00000153933 | DGKE | protein_coding |
| Huh7 | alfa | ENSG00000186994 | KANK3 | protein_coding |
| Huh7 | alfa | ENSG00000214140 | PRCD | protein_coding |
| Huh7 | alfa | ENSG00000272034 | SNORD14A | snoRNA |
| Huh7 | alfa | ENSG00000279861 | | TEC |
| Huh7 | alfa | ENSG00000213492 | NT5C3AP1 | transcribed_processed_pseudogene |
| Huh7 | alfa | ENSG00000186704 | DTX2P1 | transcribed_unprocessed_pseudogene |
| Huh7 | alfa | ENSG00000257489 | | transcribed_unprocessed_pseudogene |
| Huh7 | alfa | ENSG00000240216 | CPHL1P | unitary_pseudogene |
| Huh7 | alfa | ENSG00000230795 | HLA-K | unprocessed_pseudogene |
| Huh7 | alfa | ENSG00000206341 | HLA-H | unprocessed_pseudogene |
| Huh7 | beta | ENSG00000160710 | ADAR | protein_coding |
| Huh7 | beta | ENSG00000101347 | SAMHD1 | protein_coding |
| Huh7 | beta | ENSG00000173821 | RNF213 | protein_coding |
| Huh7 | beta | ENSG00000115415 | STAT1 | protein_coding |
| Huh7 | beta | ENSG00000170581 | STAT2 | protein_coding |
| Huh7 | beta | ENSG00000055332 | EIF2AK2 | protein_coding |
| Huh7 | beta | ENSG00000121060 | TRIM25 | protein_coding |
| Huh7 | beta | ENSG00000163840 | DTX3L | protein_coding |
| Huh7 | beta | ENSG00000188313 | PLSCR1 | protein_coding |
| Huh7 | beta | ENSG00000184979 | USP18 | protein_coding |
| Huh7 | beta | ENSG00000156587 | UBE2L6 | protein_coding |
| Huh7 | beta | ENSG00000138496 | PARP9 | protein_coding |
| Huh7 | beta | ENSG00000181381 | DDX60L | protein_coding |
| Huh7 | beta | ENSG00000173193 | PARP14 | protein_coding |
| Huh7 | beta | ENSG00000187608 | ISG15 | protein_coding |
| Huh7 | beta | ENSG00000111331 | OAS3 | protein_coding |
| Huh7 | beta | ENSG00000157601 | MX1 | protein_coding |
| Huh7 | beta | ENSG00000185745 | IFIT1 | protein_coding |
| Huh7 | beta | ENSG00000138035 | PNPT1 | protein_coding |
| Huh7 | beta | ENSG00000221963 | APOL6 | protein_coding |
| Huh7 | beta | ENSG00000067066 | SP100 | protein_coding |
| Huh7 | beta | ENSG00000107201 | DDX58 | protein_coding |
| Huh7 | beta | ENSG00000115267 | IFIH1 | protein_coding |
| Huh7 | beta | ENSG00000119917 | IFIT3 | protein_coding |
| Huh7 | beta | ENSG00000059378 | PARP12 | protein_coding |
| Huh7 | beta | ENSG00000168394 | TAP1 | protein_coding |
| Huh7 | beta | ENSG00000196116 | TDRD7 | protein_coding |
| Huh7 | beta | ENSG00000111335 | OAS2 | protein_coding |
| Huh7 | beta | ENSG00000135899 | SP110 | protein_coding |
| Huh7 | beta | ENSG00000178685 | PARP10 | protein_coding |
| Huh7 | beta | ENSG00000089127 | OAS1 | protein_coding |
| Huh7 | beta | ENSG00000105939 | ZC3HAV1 | protein_coding |
| Huh7 | beta | ENSG00000172936 | MYD88 | protein_coding |
| Huh7 | beta | ENSG00000155629 | PIK3AP1 | protein_coding |
| Huh7 | beta | ENSG00000133106 | EPSTI1 | protein_coding |
| Huh7 | beta | ENSG00000126709 | IFI6 | protein_coding |
| Huh7 | beta | ENSG00000152778 | IFIT5 | protein_coding |
| Huh7 | beta | ENSG00000117228 | GBP1 | protein_coding |
| Huh7 | beta | ENSG00000137628 | DDX60 | protein_coding |
| Huh7 | beta | ENSG00000108679 | LGALS3BP | protein_coding |
| Huh7 | beta | ENSG00000002549 | LAP3 | protein_coding |
| Huh7 | beta | ENSG00000134321 | RSAD2 | protein_coding |
| Huh7 | beta | ENSG00000142089 | IFITM3 | protein_coding |
| Huh7 | beta | ENSG00000165806 | CASP7 | protein_coding |
| Huh7 | beta | ENSG00000138642 | HERC6 | protein_coding |
| Huh7 | beta | ENSG00000173786 | CNP | protein_coding |
| Huh7 | beta | ENSG00000078081 | LAMP3 | protein_coding |
| Huh7 | beta | ENSG00000166710 | B2M | protein_coding |
| Huh7 | beta | ENSG00000168016 | TRANK1 | protein_coding |
| Huh7 | beta | ENSG00000124201 | ZNFX1 | protein_coding |
| Huh7 | beta | ENSG00000123609 | NMI | protein_coding |

| | | | | |
|---|---|---|---|---|
| Huh7 | beta | ENSG00000169245 | CXCL10 | protein_coding |
| Huh7 | beta | ENSG00000134326 | CMPK2 | protein_coding |
| Huh7 | beta | ENSG00000205413 | SAMD9 | protein_coding |
| Huh7 | beta | ENSG00000119922 | IFIT2 | protein_coding |
| Huh7 | beta | ENSG00000137959 | IFI44L | protein_coding |
| Huh7 | beta | ENSG00000130303 | BST2 | protein_coding |
| Huh7 | beta | ENSG00000204525 | HLA-C | protein_coding |
| Huh7 | beta | ENSG00000135114 | OASL | protein_coding |
| Huh7 | beta | ENSG00000130589 | HELZ2 | protein_coding |
| Huh7 | beta | ENSG00000185880 | TRIM69 | protein_coding |
| Huh7 | beta | ENSG00000204592 | HLA-E | protein_coding |
| Huh7 | beta | ENSG00000140853 | NLRC5 | protein_coding |
| Huh7 | beta | ENSG00000138646 | HERC5 | protein_coding |
| Huh7 | beta | ENSG00000110446 | SLC15A3 | protein_coding |
| Huh7 | beta | ENSG00000173530 | TNFRSF10D | protein_coding |
| Huh7 | beta | ENSG00000132256 | TRIM5 | protein_coding |
| Huh7 | beta | ENSG00000128335 | APOL2 | protein_coding |
| Huh7 | beta | ENSG00000132274 | TRIM22 | protein_coding |
| Huh7 | beta | ENSG00000164307 | ERAP1 | protein_coding |
| Huh7 | beta | ENSG00000100342 | APOL1 | protein_coding |
| Huh7 | beta | ENSG00000231925 | TAPBP | protein_coding |
| Huh7 | beta | ENSG00000105287 | PRKD2 | protein_coding |
| Huh7 | beta | ENSG00000132109 | TRIM21 | protein_coding |
| Huh7 | beta | ENSG00000111912 | NCOA7 | protein_coding |
| Huh7 | beta | ENSG00000140464 | PML | protein_coding |
| Huh7 | beta | ENSG00000130813 | C19orf66 | protein_coding |
| Huh7 | beta | ENSG00000068079 | IFI35 | protein_coding |
| Huh7 | beta | ENSG00000169429 | CXCL8 | protein_coding |
| Huh7 | beta | ENSG00000204267 | TAP2 | protein_coding |
| Huh7 | beta | ENSG00000182179 | UBA7 | protein_coding |
| Huh7 | beta | ENSG00000117226 | GBP3 | protein_coding |
| Huh7 | beta | ENSG00000132530 | XAF1 | protein_coding |
| Huh7 | beta | ENSG00000108669 | CYTH1 | protein_coding |
| Huh7 | beta | ENSG00000204264 | PSMB8 | protein_coding |
| Huh7 | beta | ENSG00000013374 | NUB1 | protein_coding |
| Huh7 | beta | ENSG00000152689 | RASGRP3 | protein_coding |
| Huh7 | beta | ENSG00000240065 | PSMB9 | protein_coding |
| Huh7 | beta | ENSG00000121858 | TNFSF10 | protein_coding |
| Huh7 | beta | ENSG00000168062 | BATF2 | protein_coding |
| Huh7 | beta | ENSG00000163644 | PPM1K | protein_coding |
| Huh7 | beta | ENSG00000105559 | PLEKHA4 | protein_coding |
| Huh7 | beta | ENSG00000116663 | FBXO6 | protein_coding |
| Huh7 | beta | ENSG00000172183 | ISG20 | protein_coding |
| Huh7 | beta | ENSG00000185404 | SP140L | protein_coding |
| Huh7 | beta | ENSG00000000971 | CFH | protein_coding |
| Huh7 | beta | ENSG00000026950 | BTN3A1 | protein_coding |
| Huh7 | beta | ENSG00000106785 | TRIM14 | protein_coding |
| Huh7 | beta | ENSG00000123240 | OPTN | protein_coding |
| Huh7 | beta | ENSG00000168404 | MLKL | protein_coding |
| Huh7 | beta | ENSG00000177409 | SAMD9L | protein_coding |
| Huh7 | beta | ENSG00000123992 | DNPEP | protein_coding |
| Huh7 | beta | ENSG00000010030 | ETV7 | protein_coding |
| Huh7 | beta | ENSG00000164342 | TLR3 | protein_coding |
| Huh7 | beta | ENSG00000137965 | IFI44 | protein_coding |
| Huh7 | beta | ENSG00000186470 | BTN3A2 | protein_coding |
| Huh7 | beta | ENSG00000125347 | IRF1 | protein_coding |
| Huh7 | beta | ENSG00000234745 | HLA-B | protein_coding |
| Huh7 | beta | ENSG00000168297 | PXK | protein_coding |
| Huh7 | beta | ENSG00000258441 | LINC00641 | processed_transcript |
| Huh7 | beta | ENSG00000079385 | CEACAM1 | protein_coding |
| Huh7 | beta | ENSG00000163545 | NUAK2 | protein_coding |
| Huh7 | beta | ENSG00000118503 | TNFAIP3 | protein_coding |

| | | | | |
|---|---|---|---|---|
| Huh7 | beta | ENSG00000182326 | C1S | protein_coding |
| Huh7 | beta | ENSG00000173702 | MUC13 | protein_coding |
| Huh7 | beta | ENSG00000117475 | BLZF1 | protein_coding |
| Huh7 | beta | ENSG00000113441 | LNPEP | protein_coding |
| Huh7 | beta | ENSG00000162654 | GBP4 | protein_coding |
| Huh7 | beta | ENSG00000165949 | IFI27 | protein_coding |
| Huh7 | beta | ENSG00000146859 | TMEM140 | protein_coding |
| Huh7 | beta | ENSG00000122643 | NT5C3A | protein_coding |
| Huh7 | beta | ENSG00000092010 | PSME1 | protein_coding |
| Huh7 | beta | ENSG00000225886 | | antisense |
| Huh7 | beta | ENSG00000064012 | CASP8 | protein_coding |
| Huh7 | beta | ENSG00000213928 | IRF9 | protein_coding |
| Huh7 | beta | ENSG00000178719 | GRINA | protein_coding |
| Huh7 | beta | ENSG00000145623 | OSMR | protein_coding |
| Huh7 | beta | ENSG00000164308 | ERAP2 | protein_coding |
| Huh7 | beta | ENSG00000114450 | GNB4 | protein_coding |
| Huh7 | beta | ENSG00000100911 | PSME2 | protein_coding |
| Huh7 | beta | ENSG00000136147 | PHF11 | protein_coding |
| Huh7 | beta | ENSG00000137200 | CMTR1 | protein_coding |
| Huh7 | beta | ENSG00000067082 | KLF6 | protein_coding |
| Huh7 | beta | ENSG00000078018 | MAP2 | protein_coding |
| Huh7 | beta | ENSG00000151025 | GPR158 | protein_coding |
| Huh7 | beta | ENSG00000185885 | IFITM1 | protein_coding |
| Huh7 | beta | ENSG00000100918 | REC8 | protein_coding |
| Huh7 | beta | ENSG00000269640 | | lincRNA |
| Huh7 | beta | ENSG00000184898 | RBM43 | protein_coding |
| Huh7 | beta | ENSG00000023445 | BIRC3 | protein_coding |
| Huh7 | beta | ENSG00000060491 | OGFR | protein_coding |
| Huh7 | beta | ENSG00000120217 | CD274 | protein_coding |
| Huh7 | beta | ENSG00000156966 | B3GNT7 | protein_coding |
| Huh7 | beta | ENSG00000133321 | RARRES3 | protein_coding |
| Huh7 | beta | ENSG00000164136 | IL15 | protein_coding |
| Huh7 | beta | ENSG00000166689 | PLEKHA7 | protein_coding |
| Huh7 | beta | ENSG00000155363 | MOV10 | protein_coding |
| Huh7 | beta | ENSG00000134716 | CYP2J2 | protein_coding |
| Huh7 | beta | ENSG00000169248 | CXCL11 | protein_coding |
| Huh7 | beta | ENSG00000104518 | GSDMD | protein_coding |
| Huh7 | beta | ENSG00000197536 | C5orf56 | protein_coding |
| Huh7 | beta | ENSG00000155287 | SLC25A28 | protein_coding |
| Huh7 | beta | ENSG00000142961 | MOB3C | protein_coding |
| Huh7 | beta | ENSG00000134470 | IL15RA | protein_coding |
| Huh7 | beta | ENSG00000161091 | MFSD12 | protein_coding |
| Huh7 | beta | ENSG00000162687 | KCNT2 | protein_coding |
| Huh7 | beta | ENSG00000137842 | TMEM62 | protein_coding |
| Huh7 | beta | ENSG00000127666 | TICAM1 | protein_coding |
| Huh7 | beta | ENSG00000163739 | CXCL1 | protein_coding |
| Huh7 | beta | ENSG00000108771 | DHX58 | protein_coding |
| Huh7 | beta | ENSG00000213500 | LAP3P2 | processed_pseudogene |
| Huh7 | beta | ENSG00000142677 | IL22RA1 | protein_coding |
| Huh7 | beta | ENSG00000242539 | | antisense |
| Huh7 | beta | ENSG00000166016 | ABTB2 | protein_coding |
| Huh7 | beta | ENSG00000184371 | CSF1 | protein_coding |
| Huh7 | beta | ENSG00000177989 | ODF3B | protein_coding |
| Huh7 | beta | ENSG00000106829 | TLE4 | protein_coding |
| Huh7 | beta | ENSG00000269728 | | lincRNA |
| Huh7 | beta | ENSG00000125826 | RBCK1 | protein_coding |
| Huh7 | beta | ENSG00000169871 | TRIM56 | protein_coding |
| Huh7 | beta | ENSG00000151470 | C4orf33 | protein_coding |
| Huh7 | beta | ENSG00000225963 | | antisense |
| Huh7 | beta | ENSG00000130487 | KLHDC7B | protein_coding |
| Huh7 | beta | ENSG00000162772 | ATF3 | protein_coding |
| Huh7 | beta | ENSG00000229241 | PNPT1P1 | processed_pseudogene |

| | | | | |
|---|---|---|---|---|
| Huh7 | beta | ENSG00000157303 | SUSD3 | protein_coding |
| Huh7 | beta | ENSG00000163823 | CCR1 | protein_coding |
| Huh7 | beta | ENSG00000159403 | C1R | protein_coding |
| Huh7 | beta | ENSG00000114268 | PFKFB4 | protein_coding |
| Huh7 | beta | ENSG00000110057 | UNC93B1 | protein_coding |
| Huh7 | beta | ENSG00000139083 | ETV6 | protein_coding |
| Huh7 | beta | ENSG00000168310 | IRF2 | protein_coding |
| Huh7 | beta | ENSG00000144596 | GRIP2 | protein_coding |
| Huh7 | beta | ENSG00000136514 | RTP4 | protein_coding |
| Huh7 | beta | ENSG00000128394 | APOBEC3F | protein_coding |
| Huh7 | beta | ENSG00000238000 | | processed_pseudogene |
| Huh7 | beta | ENSG00000214872 | SMTNL1 | protein_coding |
| Huh7 | beta | ENSG00000206341 | HLA-H | unprocessed_pseudogene |
| Huh7 | beta | ENSG00000112343 | TRIM38 | protein_coding |
| Huh7 | beta | ENSG00000141655 | TNFRSF11A | protein_coding |
| Huh7 | beta | ENSG00000141682 | PMAIP1 | protein_coding |
| Huh7 | beta | ENSG00000008441 | NFIX | protein_coding |
| Huh7 | beta | ENSG00000228318 | | antisense |
| Huh7 | beta | ENSG00000229474 | PATL2 | protein_coding |
| Huh7 | beta | ENSG00000111801 | BTN3A3 | protein_coding |
| Huh7 | beta | ENSG00000149131 | SERPING1 | protein_coding |
| Huh7 | beta | ENSG00000103056 | SMPD3 | protein_coding |
| Huh7 | beta | ENSG00000243650 | RN7SL834P | misc_RNA |
| Huh7 | beta | ENSG00000225131 | PSME2P2 | processed_pseudogene |
| Huh7 | beta | ENSG00000156500 | FAM122C | protein_coding |
| Huh7 | beta | ENSG00000272512 | | lincRNA |
| Huh7 | beta | ENSG00000230795 | HLA-K | unprocessed_pseudogene |
| Huh7 | beta | ENSG00000105402 | NAPA | protein_coding |
| Huh7 | beta | ENSG00000163121 | NEURL3 | protein_coding |
| Huh7 | beta | ENSG00000185201 | IFITM2 | protein_coding |
| Huh7 | beta | ENSG00000072858 | SIDT1 | protein_coding |
| Huh7 | beta | ENSG00000263069 | | antisense |
| Huh7 | beta | ENSG00000141664 | ZCCHC2 | protein_coding |
| Huh7 | beta | ENSG00000248713 | | protein_coding |
| Huh7 | beta | ENSG00000137752 | CASP1 | protein_coding |
| Huh7 | beta | ENSG00000204397 | CARD16 | protein_coding |
| Huh7 | beta | ENSG00000137767 | SQRDL | protein_coding |
| Huh7 | beta | ENSG00000266074 | BAHCC1 | protein_coding |
| Huh7 | beta | ENSG00000168497 | SDPR | protein_coding |
| Huh7 | beta | ENSG00000214226 | C17orf67 | protein_coding |
| Huh7 | beta | ENSG00000140511 | HAPLN3 | protein_coding |
| Huh7 | beta | ENSG00000259529 | IRF9 | protein_coding |
| Huh7 | beta | ENSG00000106302 | HYAL4 | protein_coding |
| Huh7 | beta | ENSG00000206503 | HLA-A | protein_coding |
| Huh7 | beta | ENSG00000081041 | CXCL2 | protein_coding |
| Huh7 | beta | ENSG00000205220 | PSMB10 | protein_coding |
| Huh7 | beta | ENSG00000144852 | NR1I2 | protein_coding |
| Huh7 | beta | ENSG00000255398 | HCAR3 | protein_coding |
| Huh7 | beta | ENSG00000263528 | IKBKE | protein_coding |
| Huh7 | beta | ENSG00000133943 | C14orf159 | protein_coding |
| Huh7 | beta | ENSG00000139192 | TAPBPL | protein_coding |
| Huh7 | beta | ENSG00000126778 | SIX1 | protein_coding |
| Huh7 | beta | ENSG00000185022 | MAFF | protein_coding |
| Huh7 | beta | ENSG00000181541 | MAB21L2 | protein_coding |
| Huh7 | beta | ENSG00000271550 | BNIP3P11 | processed_pseudogene |
| Huh7 | beta | ENSG00000163661 | PTX3 | protein_coding |
| Huh7 | beta | ENSG00000128342 | LIF | protein_coding |
| Huh7 | beta | ENSG00000107593 | PKD2L1 | protein_coding |
| Huh7 | beta | ENSG00000088881 | EBF4 | protein_coding |
| Huh7 | beta | ENSG00000188613 | NANOS1 | protein_coding |
| Huh7 | beta | ENSG00000118322 | ATP10B | protein_coding |
| Huh7 | beta | ENSG00000276980 | | sense_intronic |

| | | | | |
|---|---|---|---|---|
| Huh7 | beta | ENSG00000025708 | TYMP | protein_coding |
| Huh7 | beta | ENSG00000245571 | | lincRNA |
| Huh7 | beta | ENSG00000187583 | PLEKHN1 | protein_coding |
| Huh7 | beta | ENSG00000254132 | MTND6P3 | processed_pseudogene |
| Huh7 | beta | ENSG00000248988 | | processed_pseudogene |
| Huh7 | beta | ENSG00000279861 | | TEC |
| Huh7 | beta | ENSG00000228118 | | processed_pseudogene |
| Huh7 | beta | ENSG00000188290 | HES4 | protein_coding |
| Huh7 | beta | ENSG00000213886 | UBD | protein_coding |
| Huh7 | beta | ENSG00000166741 | NNMT | protein_coding |
| Huh7 | beta | ENSG00000225075 | | antisense |
| Huh7 | beta | ENSG00000008735 | MAPK8IP2 | protein_coding |
| Huh7 | beta | ENSG00000183022 | TPM3P8 | processed_pseudogene |
| Huh7 | beta | ENSG00000229873 | OGFR-AS1 | antisense |
| Huh7 | beta | ENSG00000186994 | KANK3 | protein_coding |
| Huh7 | beta | ENSG00000183397 | C19orf71 | protein_coding |
| Huh7 | beta | ENSG00000260966 | | sense_overlapping |
| Huh7 | beta | ENSG00000184058 | TBX1 | protein_coding |
| Huh7 | beta | ENSG00000157873 | TNFRSF14 | protein_coding |
| Huh7 | beta | ENSG00000196664 | TLR7 | protein_coding |
| Huh7 | beta | ENSG00000167601 | AXL | protein_coding |
| Huh7 | beta | ENSG00000104432 | IL7 | protein_coding |
| Huh7 | beta | ENSG00000260302 | | lincRNA |
| Huh7 | beta | ENSG00000233817 | | lincRNA |
| Huh7 | beta | ENSG00000280007 | | antisense |
| Huh7 | beta | ENSG00000184454 | NCMAP | protein_coding |
| Huh7 | lambda | ENSG00000160710 | ADAR | protein_coding |
| Huh7 | lambda | ENSG00000115415 | STAT1 | protein_coding |
| Huh7 | lambda | ENSG00000055332 | EIF2AK2 | protein_coding |
| Huh7 | lambda | ENSG00000163840 | DTX3L | protein_coding |
| Huh7 | lambda | ENSG00000173193 | PARP14 | protein_coding |
| Huh7 | lambda | ENSG00000138496 | PARP9 | protein_coding |
| Huh7 | lambda | ENSG00000187608 | ISG15 | protein_coding |
| Huh7 | lambda | ENSG00000157601 | MX1 | protein_coding |
| Huh7 | lambda | ENSG00000188313 | PLSCR1 | protein_coding |
| Huh7 | lambda | ENSG00000185745 | IFIT1 | protein_coding |
| Huh7 | lambda | ENSG00000121060 | TRIM25 | protein_coding |
| Huh7 | lambda | ENSG00000111331 | OAS3 | protein_coding |
| Huh7 | lambda | ENSG00000105939 | ZC3HAV1 | protein_coding |
| Huh7 | lambda | ENSG00000126709 | IFI6 | protein_coding |
| Huh7 | lambda | ENSG00000181381 | DDX60L | protein_coding |
| Huh7 | lambda | ENSG00000115267 | IFIH1 | protein_coding |
| Huh7 | lambda | ENSG00000107201 | DDX58 | protein_coding |
| Huh7 | lambda | ENSG00000184979 | USP18 | protein_coding |
| Huh7 | lambda | ENSG00000152778 | IFIT5 | protein_coding |
| Huh7 | lambda | ENSG00000119917 | IFIT3 | protein_coding |
| Huh7 | lambda | ENSG00000133106 | EPSTI1 | protein_coding |
| Huh7 | lambda | ENSG00000059378 | PARP12 | protein_coding |
| Huh7 | lambda | ENSG00000137628 | DDX60 | protein_coding |
| Huh7 | lambda | ENSG00000135899 | SP110 | protein_coding |
| Huh7 | lambda | ENSG00000138642 | HERC6 | protein_coding |
| Huh7 | lambda | ENSG00000067066 | SP100 | protein_coding |
| Huh7 | lambda | ENSG00000130589 | HELZ2 | protein_coding |
| Huh7 | lambda | ENSG00000142089 | IFITM3 | protein_coding |
| Huh7 | lambda | ENSG00000221963 | APOL6 | protein_coding |
| Huh7 | lambda | ENSG00000168394 | TAP1 | protein_coding |
| Huh7 | lambda | ENSG00000089127 | OAS1 | protein_coding |
| Huh7 | lambda | ENSG00000170581 | STAT2 | protein_coding |
| Huh7 | lambda | ENSG00000196116 | TDRD7 | protein_coding |
| Huh7 | lambda | ENSG00000178685 | PARP10 | protein_coding |
| Huh7 | lambda | ENSG00000134326 | CMPK2 | protein_coding |
| Huh7 | lambda | ENSG00000124201 | ZNFX1 | protein_coding |

| | | | | |
|---|---|---|---|---|
| Huh7 | lambda | ENSG00000205413 | SAMD9 | protein_coding |
| Huh7 | lambda | ENSG00000111335 | OAS2 | protein_coding |
| Huh7 | lambda | ENSG00000156587 | UBE2L6 | protein_coding |
| Huh7 | lambda | ENSG00000101347 | SAMHD1 | protein_coding |
| Huh7 | lambda | ENSG00000140464 | PML | protein_coding |
| Huh7 | lambda | ENSG00000108679 | LGALS3BP | protein_coding |
| Huh7 | lambda | ENSG00000106785 | TRIM14 | protein_coding |
| Huh7 | lambda | ENSG00000132109 | TRIM21 | protein_coding |
| Huh7 | lambda | ENSG00000117228 | GBP1 | protein_coding |
| Huh7 | lambda | ENSG00000119922 | IFIT2 | protein_coding |
| Huh7 | lambda | ENSG00000213928 | IRF9 | protein_coding |
| Huh7 | lambda | ENSG00000185880 | TRIM69 | protein_coding |
| Huh7 | lambda | ENSG00000138646 | HERC5 | protein_coding |
| Huh7 | lambda | ENSG00000100918 | REC8 | protein_coding |
| Huh7 | lambda | ENSG00000225886 | | antisense |
| Huh7 | lambda | ENSG00000110446 | SLC15A3 | protein_coding |
| Huh7 | lambda | ENSG00000137959 | IFI44L | protein_coding |
| Huh7 | lambda | ENSG00000130303 | BST2 | protein_coding |
| Huh7 | lambda | ENSG00000130813 | C19orf66 | protein_coding |
| Huh7 | lambda | ENSG00000134321 | RSAD2 | protein_coding |
| Huh7 | lambda | ENSG00000135114 | OASL | protein_coding |
| Huh7 | lambda | ENSG00000168016 | TRANK1 | protein_coding |
| Huh7 | lambda | ENSG00000060491 | OGFR | protein_coding |
| Huh7 | lambda | ENSG00000140853 | NLRC5 | protein_coding |
| Huh7 | lambda | ENSG00000168062 | BATF2 | protein_coding |
| Huh7 | lambda | ENSG00000068079 | IFI35 | protein_coding |
| Huh7 | lambda | ENSG00000136147 | PHF11 | protein_coding |
| Huh7 | lambda | ENSG00000204525 | HLA-C | protein_coding |
| Huh7 | lambda | ENSG00000078081 | LAMP3 | protein_coding |
| Huh7 | lambda | ENSG00000132530 | XAF1 | protein_coding |
| Huh7 | lambda | ENSG00000204267 | TAP2 | protein_coding |
| Huh7 | lambda | ENSG00000100342 | APOL1 | protein_coding |
| Huh7 | lambda | ENSG00000259529 | IRF9 | protein_coding |
| Huh7 | lambda | ENSG00000177409 | SAMD9L | protein_coding |
| Huh7 | lambda | ENSG00000177989 | ODF3B | protein_coding |
| Huh7 | lambda | ENSG00000117226 | GBP3 | protein_coding |
| Huh7 | lambda | ENSG00000182179 | UBA7 | protein_coding |
| Huh7 | lambda | ENSG00000204264 | PSMB8 | protein_coding |
| Huh7 | lambda | ENSG00000130487 | KLHDC7B | protein_coding |
| Huh7 | lambda | ENSG00000229241 | PNPT1P1 | processed_pseudogene |
| Huh7 | lambda | ENSG00000110057 | UNC93B1 | protein_coding |
| Huh7 | lambda | ENSG00000240065 | PSMB9 | protein_coding |
| Huh7 | lambda | ENSG00000132274 | TRIM22 | protein_coding |
| Huh7.5 | alfa | ENSG00000160710 | ADAR | protein_coding |
| Huh7.5 | alfa | ENSG00000196116 | TDRD7 | protein_coding |
| Huh7.5 | alfa | ENSG00000173821 | RNF213 | protein_coding |
| Huh7.5 | alfa | ENSG00000138035 | PNPT1 | protein_coding |
| Huh7.5 | alfa | ENSG00000101347 | SAMHD1 | protein_coding |
| Huh7.5 | alfa | ENSG00000170581 | STAT2 | protein_coding |
| Huh7.5 | alfa | ENSG00000121060 | TRIM25 | protein_coding |
| Huh7.5 | alfa | ENSG00000055332 | EIF2AK2 | protein_coding |
| Huh7.5 | alfa | ENSG00000115415 | STAT1 | protein_coding |
| Huh7.5 | alfa | ENSG00000163840 | DTX3L | protein_coding |
| Huh7.5 | alfa | ENSG00000067066 | SP100 | protein_coding |
| Huh7.5 | alfa | ENSG00000188313 | PLSCR1 | protein_coding |
| Huh7.5 | alfa | ENSG00000184979 | USP18 | protein_coding |
| Huh7.5 | alfa | ENSG00000138496 | PARP9 | protein_coding |
| Huh7.5 | alfa | ENSG00000107201 | DDX58 | protein_coding |
| Huh7.5 | alfa | ENSG00000181381 | DDX60L | protein_coding |
| Huh7.5 | alfa | ENSG00000117228 | GBP1 | protein_coding |
| Huh7.5 | alfa | ENSG00000156587 | UBE2L6 | protein_coding |
| Huh7.5 | alfa | ENSG00000173193 | PARP14 | protein_coding |

| | | | | |
|---|---|---|---|---|
| Huh7.5 | alfa | ENSG00000221963 | APOL6 | protein_coding |
| Huh7.5 | alfa | ENSG00000187608 | ISG15 | protein_coding |
| Huh7.5 | alfa | ENSG00000119917 | IFIT3 | protein_coding |
| Huh7.5 | alfa | ENSG00000137628 | DDX60 | protein_coding |
| Huh7.5 | alfa | ENSG00000111331 | OAS3 | protein_coding |
| Huh7.5 | alfa | ENSG00000115267 | IFIH1 | protein_coding |
| Huh7.5 | alfa | ENSG00000185745 | IFIT1 | protein_coding |
| Huh7.5 | alfa | ENSG00000157601 | MX1 | protein_coding |
| Huh7.5 | alfa | ENSG00000169245 | CXCL10 | protein_coding |
| Huh7.5 | alfa | ENSG00000002549 | LAP3 | protein_coding |
| Huh7.5 | alfa | ENSG00000168394 | TAP1 | protein_coding |
| Huh7.5 | alfa | ENSG00000126709 | IFI6 | protein_coding |
| Huh7.5 | alfa | ENSG00000089127 | OAS1 | protein_coding |
| Huh7.5 | alfa | ENSG00000178685 | PARP10 | protein_coding |
| Huh7.5 | alfa | ENSG00000111335 | OAS2 | protein_coding |
| Huh7.5 | alfa | ENSG00000138642 | HERC6 | protein_coding |
| Huh7.5 | alfa | ENSG00000172936 | MYD88 | protein_coding |
| Huh7.5 | alfa | ENSG00000123609 | NMI | protein_coding |
| Huh7.5 | alfa | ENSG00000173786 | CNP | protein_coding |
| Huh7.5 | alfa | ENSG00000135899 | SP110 | protein_coding |
| Huh7.5 | alfa | ENSG00000155629 | PIK3AP1 | protein_coding |
| Huh7.5 | alfa | ENSG00000059378 | PARP12 | protein_coding |
| Huh7.5 | alfa | ENSG00000134321 | RSAD2 | protein_coding |
| Huh7.5 | alfa | ENSG00000152778 | IFIT5 | protein_coding |
| Huh7.5 | alfa | ENSG00000105939 | ZC3HAV1 | protein_coding |
| Huh7.5 | alfa | ENSG00000108679 | LGALS3BP | protein_coding |
| Huh7.5 | alfa | ENSG00000142089 | IFITM3 | protein_coding |
| Huh7.5 | alfa | ENSG00000205413 | SAMD9 | protein_coding |
| Huh7.5 | alfa | ENSG00000166710 | B2M | protein_coding |
| Huh7.5 | alfa | ENSG00000124201 | ZNFX1 | protein_coding |
| Huh7.5 | alfa | ENSG00000137959 | IFI44L | protein_coding |
| Huh7.5 | alfa | ENSG00000134326 | CMPK2 | protein_coding |
| Huh7.5 | alfa | ENSG00000130303 | BST2 | protein_coding |
| Huh7.5 | alfa | ENSG00000138646 | HERC5 | protein_coding |
| Huh7.5 | alfa | ENSG00000204592 | HLA-E | protein_coding |
| Huh7.5 | alfa | ENSG00000119922 | IFIT2 | protein_coding |
| Huh7.5 | alfa | ENSG00000130813 | C19orf66 | protein_coding |
| Huh7.5 | alfa | ENSG00000133106 | EPSTI1 | protein_coding |
| Huh7.5 | alfa | ENSG00000135114 | OASL | protein_coding |
| Huh7.5 | alfa | ENSG00000165806 | CASP7 | protein_coding |
| Huh7.5 | alfa | ENSG00000140853 | NLRC5 | protein_coding |
| Huh7.5 | alfa | ENSG00000140464 | PML | protein_coding |
| Huh7.5 | alfa | ENSG00000105287 | PRKD2 | protein_coding |
| Huh7.5 | alfa | ENSG00000130589 | HELZ2 | protein_coding |
| Huh7.5 | alfa | ENSG00000110446 | SLC15A3 | protein_coding |
| Huh7.5 | alfa | ENSG00000182179 | UBA7 | protein_coding |
| Huh7.5 | alfa | ENSG00000164307 | ERAP1 | protein_coding |
| Huh7.5 | alfa | ENSG00000013374 | NUB1 | protein_coding |
| Huh7.5 | alfa | ENSG00000137965 | IFI44 | protein_coding |
| Huh7.5 | alfa | ENSG00000204267 | TAP2 | protein_coding |
| Huh7.5 | alfa | ENSG00000132274 | TRIM22 | protein_coding |
| Huh7.5 | alfa | ENSG00000132109 | TRIM21 | protein_coding |
| Huh7.5 | alfa | ENSG00000117226 | GBP3 | protein_coding |
| Huh7.5 | alfa | ENSG00000128335 | APOL2 | protein_coding |
| Huh7.5 | alfa | ENSG00000132256 | TRIM5 | protein_coding |
| Huh7.5 | alfa | ENSG00000182326 | C1S | protein_coding |
| Huh7.5 | alfa | ENSG00000132530 | XAF1 | protein_coding |
| Huh7.5 | alfa | ENSG00000078081 | LAMP3 | protein_coding |
| Huh7.5 | alfa | ENSG00000185880 | TRIM69 | protein_coding |
| Huh7.5 | alfa | ENSG00000204525 | HLA-C | protein_coding |
| Huh7.5 | alfa | ENSG00000079385 | CEACAM1 | protein_coding |
| Huh7.5 | alfa | ENSG00000231925 | TAPBP | protein_coding |

| | | | | |
|---|---|---|---|---|
| Huh7.5 | alfa | ENSG00000185404 | SP140L | protein_coding |
| Huh7.5 | alfa | ENSG00000100342 | APOL1 | protein_coding |
| Huh7.5 | alfa | ENSG00000121858 | TNFSF10 | protein_coding |
| Huh7.5 | alfa | ENSG00000240065 | PSMB9 | protein_coding |
| Huh7.5 | alfa | ENSG00000111912 | NCOA7 | protein_coding |
| Huh7.5 | alfa | ENSG00000064012 | CASP8 | protein_coding |
| Huh7.5 | alfa | ENSG00000168016 | TRANK1 | protein_coding |
| Huh7.5 | alfa | ENSG00000177409 | SAMD9L | protein_coding |
| Huh7.5 | alfa | ENSG00000172183 | ISG20 | protein_coding |
| Huh7.5 | alfa | ENSG00000120539 | MASTL | protein_coding |
| Huh7.5 | alfa | ENSG00000116663 | FBXO6 | protein_coding |
| Huh7.5 | alfa | ENSG00000105835 | NAMPT | protein_coding |
| Huh7.5 | alfa | ENSG00000204264 | PSMB8 | protein_coding |
| Huh7.5 | alfa | ENSG00000106785 | TRIM14 | protein_coding |
| Huh7.5 | alfa | ENSG00000123240 | OPTN | protein_coding |
| Huh7.5 | alfa | ENSG00000163644 | PPM1K | protein_coding |
| Huh7.5 | alfa | ENSG00000173530 | TNFRSF10D | protein_coding |
| Huh7.5 | alfa | ENSG00000152689 | RASGRP3 | protein_coding |
| Huh7.5 | alfa | ENSG00000108669 | CYTH1 | protein_coding |
| Huh7.5 | alfa | ENSG00000026950 | BTN3A1 | protein_coding |
| Huh7.5 | alfa | ENSG00000165949 | IFI27 | protein_coding |
| Huh7.5 | alfa | ENSG00000105559 | PLEKHA4 | protein_coding |
| Huh7.5 | alfa | ENSG00000168062 | BATF2 | protein_coding |
| Huh7.5 | alfa | ENSG00000068079 | IFI35 | protein_coding |
| Huh7.5 | alfa | ENSG00000164342 | TLR3 | protein_coding |
| Huh7.5 | alfa | ENSG00000114450 | GNB4 | protein_coding |
| Huh7.5 | alfa | ENSG00000180628 | PCGF5 | protein_coding |
| Huh7.5 | alfa | ENSG00000155363 | MOV10 | protein_coding |
| Huh7.5 | alfa | ENSG00000173702 | MUC13 | protein_coding |
| Huh7.5 | alfa | ENSG00000100911 | PSME2 | protein_coding |
| Huh7.5 | alfa | ENSG00000234745 | HLA-B | protein_coding |
| Huh7.5 | alfa | ENSG00000159403 | C1R | protein_coding |
| Huh7.5 | alfa | ENSG00000117475 | BLZF1 | protein_coding |
| Huh7.5 | alfa | ENSG00000213928 | IRF9 | protein_coding |
| Huh7.5 | alfa | ENSG00000169248 | CXCL11 | protein_coding |
| Huh7.5 | alfa | ENSG00000184898 | RBM43 | protein_coding |
| Huh7.5 | alfa | ENSG00000168297 | PXK | protein_coding |
| Huh7.5 | alfa | ENSG00000225886 | | antisense |
| Huh7.5 | alfa | ENSG00000137200 | CMTR1 | protein_coding |
| Huh7.5 | alfa | ENSG00000100918 | REC8 | protein_coding |
| Huh7.5 | alfa | ENSG00000092010 | PSME1 | protein_coding |
| Huh7.5 | alfa | ENSG00000113441 | LNPEP | protein_coding |
| Huh7.5 | alfa | ENSG00000078018 | MAP2 | protein_coding |
| Huh7.5 | alfa | ENSG00000269640 | | lincRNA |
| Huh7.5 | alfa | ENSG00000133321 | RARRES3 | protein_coding |
| Huh7.5 | alfa | ENSG00000123992 | DNPEP | protein_coding |
| Huh7.5 | alfa | ENSG00000186470 | BTN3A2 | protein_coding |
| Huh7.5 | alfa | ENSG00000136147 | PHF11 | protein_coding |
| Huh7.5 | alfa | ENSG00000168404 | MLKL | protein_coding |
| Huh7.5 | alfa | ENSG00000164308 | ERAP2 | protein_coding |
| Huh7.5 | alfa | ENSG00000004468 | CD38 | protein_coding |
| Huh7.5 | alfa | ENSG00000162645 | GBP2 | protein_coding |
| Huh7.5 | alfa | ENSG00000137752 | CASP1 | protein_coding |
| Huh7.5 | alfa | ENSG00000010030 | ETV7 | protein_coding |
| Huh7.5 | alfa | ENSG00000120217 | CD274 | protein_coding |
| Huh7.5 | alfa | ENSG00000151025 | GPR158 | protein_coding |
| Huh7.5 | alfa | ENSG00000146859 | TMEM140 | protein_coding |
| Huh7.5 | alfa | ENSG00000060491 | OGFR | protein_coding |
| Huh7.5 | alfa | ENSG00000164136 | IL15 | protein_coding |
| Huh7.5 | alfa | ENSG00000177989 | ODF3B | protein_coding |
| Huh7.5 | alfa | ENSG00000136874 | STX17 | protein_coding |
| Huh7.5 | alfa | ENSG00000134716 | CYP2J2 | protein_coding |

| | | | | |
|---|---|---|---|---|
| Huh7.5 | alfa | ENSG00000196247 | ZNF107 | protein_coding |
| Huh7.5 | alfa | ENSG00000142961 | MOB3C | protein_coding |
| Huh7.5 | alfa | ENSG00000162687 | KCNT2 | protein_coding |
| Huh7.5 | alfa | ENSG00000137842 | TMEM62 | protein_coding |
| Huh7.5 | alfa | ENSG00000185885 | IFITM1 | protein_coding |
| Huh7.5 | alfa | ENSG00000229644 | NAMPTL | processed_pseudogene |
| Huh7.5 | alfa | ENSG00000144596 | GRIP2 | protein_coding |
| Huh7.5 | alfa | ENSG00000125826 | RBCK1 | protein_coding |
| Huh7.5 | alfa | ENSG00000155287 | SLC25A28 | protein_coding |
| Huh7.5 | alfa | ENSG00000238000 | | processed_pseudogene |
| Huh7.5 | alfa | ENSG00000214872 | SMTNL1 | protein_coding |
| Huh7.5 | alfa | ENSG00000149131 | SERPING1 | protein_coding |
| Huh7.5 | alfa | ENSG00000136514 | RTP4 | protein_coding |
| Huh7.5 | alfa | ENSG00000128394 | APOBEC3F | protein_coding |
| Huh7.5 | alfa | ENSG00000161091 | MFSD12 | protein_coding |
| Huh7.5 | alfa | ENSG00000132744 | ACY3 | protein_coding |
| Huh7.5 | alfa | ENSG00000184371 | CSF1 | protein_coding |
| Huh7.5 | alfa | ENSG00000223960 | | antisense |
| Huh7.5 | alfa | ENSG00000242539 | | antisense |
| Huh7.5 | alfa | ENSG00000229241 | PNPT1P1 | processed_pseudogene |
| Huh7.5 | alfa | ENSG00000197536 | C5orf56 | protein_coding |
| Huh7.5 | alfa | ENSG00000169871 | TRIM56 | protein_coding |
| Huh7.5 | alfa | ENSG00000229474 | PATL2 | protein_coding |
| Huh7.5 | alfa | ENSG00000151466 | SCLT1 | protein_coding |
| Huh7.5 | alfa | ENSG00000073605 | GSDMB | protein_coding |
| Huh7.5 | alfa | ENSG00000140511 | HAPLN3 | protein_coding |
| Huh7.5 | alfa | ENSG00000225131 | PSME2P2 | processed_pseudogene |
| Huh7.5 | alfa | ENSG00000142677 | IL22RA1 | protein_coding |
| Huh7.5 | alfa | ENSG00000110057 | UNC93B1 | protein_coding |
| Huh7.5 | alfa | ENSG00000228318 | | antisense |
| Huh7.5 | alfa | ENSG00000156966 | B3GNT7 | protein_coding |
| Huh7.5 | alfa | ENSG00000213500 | LAP3P2 | processed_pseudogene |
| Huh7.5 | alfa | ENSG00000151470 | C4orf33 | protein_coding |
| Huh7.5 | alfa | ENSG00000135378 | PRRG4 | protein_coding |
| Huh7.5 | alfa | ENSG00000243650 | RN7SL834P | misc_RNA |
| Huh7.5 | alfa | ENSG00000141655 | TNFRSF11A | protein_coding |
| Huh7.5 | alfa | ENSG00000112343 | TRIM38 | protein_coding |
| Huh7.5 | alfa | ENSG00000269728 | | lincRNA |
| Huh7.5 | alfa | ENSG00000105402 | NAPA | protein_coding |
| Huh7.5 | alfa | ENSG00000225963 | | antisense |
| Huh7.5 | alfa | ENSG00000163823 | CCR1 | protein_coding |
| Huh7.5 | alfa | ENSG00000130487 | KLHDC7B | protein_coding |
| Huh7.5 | alfa | ENSG00000206341 | HLA-H | unprocessed_pseudogene |
| Huh7.5 | alfa | ENSG00000214226 | C17orf67 | protein_coding |
| Huh7.5 | alfa | ENSG00000162654 | GBP4 | protein_coding |
| Huh7.5 | alfa | ENSG00000072858 | SIDT1 | protein_coding |
| Huh7.5 | alfa | ENSG00000156500 | FAM122C | protein_coding |
| Huh7.5 | alfa | ENSG00000205364 | MT1M | protein_coding |
| Huh7.5 | alfa | ENSG00000259529 | IRF9 | protein_coding |
| Huh7.5 | alfa | ENSG00000108771 | DHX58 | protein_coding |
| Huh7.5 | alfa | ENSG00000141664 | ZCCHC2 | protein_coding |
| Huh7.5 | alfa | ENSG00000187583 | PLEKHN1 | protein_coding |
| Huh7.5 | alfa | ENSG00000169894 | MUC3A | protein_coding |
| Huh7.5 | alfa | ENSG00000230795 | HLA-K | unprocessed_pseudogene |
| Huh7.5 | alfa | ENSG00000144852 | NR1I2 | protein_coding |
| Huh7.5 | alfa | ENSG00000272452 | | lincRNA |
| Huh7.5 | alfa | ENSG00000225492 | GBP1P1 | transcribed_unprocessed_pseudogene |
| Huh7.5 | alfa | ENSG00000276980 | | sense_intronic |
| Huh7.5 | alfa | ENSG00000205220 | PSMB10 | protein_coding |
| Huh7.5 | alfa | ENSG00000137767 | SQRDL | protein_coding |
| Huh7.5 | alfa | ENSG00000139192 | TAPBPL | protein_coding |
| Huh7.5 | alfa | ENSG00000204397 | CARD16 | protein_coding |

| | | | | |
|---|---|---|---|---|
| Huh7.5 | alfa | ENSG00000181541 | MAB21L2 | protein_coding |
| Huh7.5 | alfa | ENSG00000157303 | SUSD3 | protein_coding |
| Huh7.5 | alfa | ENSG00000134470 | IL15RA | protein_coding |
| Huh7.5 | alfa | ENSG00000088881 | EBF4 | protein_coding |
| Huh7.5 | alfa | ENSG00000104432 | IL7 | protein_coding |
| Huh7.5 | alfa | ENSG00000124882 | EREG | protein_coding |
| Huh7.5 | alfa | ENSG00000225075 | | antisense |
| Huh7.5 | alfa | ENSG00000266074 | BAHCC1 | protein_coding |
| Huh7.5 | alfa | ENSG00000025708 | TYMP | protein_coding |
| Huh7.5 | alfa | ENSG00000103056 | SMPD3 | protein_coding |
| Huh7.5 | alfa | ENSG00000111801 | BTN3A3 | protein_coding |
| Huh7.5 | alfa | ENSG00000008735 | MAPK8IP2 | protein_coding |
| Huh7.5 | alfa | ENSG00000121380 | BCL2L14 | protein_coding |
| Huh7.5 | alfa | ENSG00000272512 | | lincRNA |
| Huh7.5 | alfa | ENSG00000042980 | ADAM28 | protein_coding |
| Huh7.5 | alfa | ENSG00000177721 | ANXA2R | protein_coding |
| Huh7.5 | alfa | ENSG00000107593 | PKD2L1 | protein_coding |
| Huh7.5 | alfa | ENSG00000196664 | TLR7 | protein_coding |
| Huh7.5 | alfa | ENSG00000188290 | HES4 | protein_coding |
| Huh7.5 | alfa | ENSG00000087085 | ACHE | protein_coding |
| Huh7.5 | alfa | ENSG00000245571 | | lincRNA |
| Huh7.5 | alfa | ENSG00000184058 | TBX1 | protein_coding |
| Huh7.5 | alfa | ENSG00000213492 | NT5C3AP1 | transcribed_processed_pseudogene |
| Huh7.5 | alfa | ENSG00000124406 | ATP8A1 | protein_coding |
| Huh7.5 | alfa | ENSG00000140832 | MARVELD3 | protein_coding |
| Huh7.5 | alfa | ENSG00000248988 | | processed_pseudogene |
| Huh7.5 | alfa | ENSG00000155158 | TTC39B | protein_coding |
| Huh7.5 | alfa | ENSG00000185201 | IFITM2 | protein_coding |
| Huh7.5 | alfa | ENSG00000232498 | | antisense |
| Huh7.5 | alfa | ENSG00000133943 | C14orf159 | protein_coding |
| Huh7.5 | alfa | ENSG00000229873 | OGFR-AS1 | antisense |
| Huh7.5 | alfa | ENSG00000157873 | TNFRSF14 | protein_coding |
| Huh7.5 | alfa | ENSG00000260302 | | lincRNA |
| Huh7.5 | alfa | ENSG00000188613 | NANOS1 | protein_coding |
| Huh7.5 | alfa | ENSG00000144820 | GPR128 | protein_coding |
| Huh7.5 | alfa | ENSG00000280007 | | antisense |
| Huh7.5 | alfa | ENSG00000275302 | CCL4 | protein_coding |
| Huh7.5 | alfa | ENSG00000185338 | SOCS1 | protein_coding |
| Huh7.5 | alfa | ENSG00000120341 | SEC16B | protein_coding |
| Huh7.5 | alfa | ENSG00000119514 | GALNT12 | protein_coding |
| Huh7.5 | alfa | ENSG00000152154 | TMEM178A | protein_coding |
| Huh7.5 | alfa | ENSG00000263069 | | antisense |
| Huh7.5 | alfa | ENSG00000249740 | OSMR-AS1 | lincRNA |
| Huh7.5 | alfa | ENSG00000255753 | | processed_pseudogene |
| Huh7.5 | alfa | ENSG00000128564 | VGF | protein_coding |
| Huh7.5 | alfa | ENSG00000243649 | CFB | protein_coding |
| Huh7.5 | alfa | ENSG00000166741 | NNMT | protein_coding |
| Huh7.5 | beta | ENSG00000253982 | | antisense |
| Huh7.5 | beta | ENSG00000173530 | TNFRSF10D | protein_coding |
| Huh7.5 | beta | ENSG00000188613 | NANOS1 | protein_coding |
| Huh7.5 | beta | ENSG00000256802 | | antisense |
| Huh7.5 | beta | ENSG00000229891 | LINC01315 | lincRNA |
| Huh7.5 | beta | ENSG00000204099 | NEU4 | protein_coding |
| Huh7.5 | beta | ENSG00000103253 | HAGHL | protein_coding |
| Huh7.5 | beta | ENSG00000142303 | ADAMTS10 | protein_coding |
| Huh7.5 | beta | ENSG00000275966 | | lincRNA |
| Huh7.5 | beta | ENSG00000188672 | RHCE | protein_coding |
| Huh7.5 | beta | ENSG00000172346 | CSDC2 | protein_coding |
| Huh7.5 | beta | ENSG00000273619 | | antisense |
| Huh7.5 | beta | ENSG00000272667 | | lincRNA |
| Huh7.5 | beta | ENSG00000171119 | NRTN | protein_coding |
| Huh7.5 | beta | ENSG00000248121 | SMURF2P1 | transcribed_unprocessed_pseudogene |

| | | | | |
|---|---|---|---|---|
| Huh7.5 | beta | ENSG00000167733 | HSD11B1L | protein_coding |
| Huh7.5 | beta | ENSG00000130675 | MNX1 | protein_coding |
| Huh7.5 | beta | ENSG00000262814 | MRPL12 | protein_coding |
| Huh7.5 | beta | ENSG00000116014 | KISS1R | protein_coding |
| Huh7.5 | beta | ENSG00000174586 | ZNF497 | protein_coding |
| Huh7.5 | beta | ENSG00000259605 | | processed_transcript |
| Huh7.5 | beta | ENSG00000205078 | SYCE1L | protein_coding |
| Huh7.5 | beta | ENSG00000162576 | MXRA8 | protein_coding |
| Huh7.5 | beta | ENSG00000233041 | PHGR1 | protein_coding |
| Huh7.5 | beta | ENSG00000167700 | MFSD3 | protein_coding |
| Huh7.5 | beta | ENSG00000168140 | VASN | protein_coding |
| Huh7.5 | beta | ENSG00000183114 | FAM43B | protein_coding |
| Huh7.5 | beta | ENSG00000197766 | CFD | protein_coding |
| Huh7.5 | beta | ENSG00000261308 | FIGNL2 | processed_pseudogene |
| Huh7.5 | beta | ENSG00000167971 | CASKIN1 | protein_coding |
| Huh7.5 | beta | ENSG00000099617 | EFNA2 | protein_coding |
| Huh7.5 | beta | ENSG00000184163 | FAM132A | protein_coding |
| Huh7.5 | beta | ENSG00000242110 | AMACR | protein_coding |
| Huh7.5 | beta | ENSG00000165886 | UBTD1 | protein_coding |
| Huh7.5 | beta | ENSG00000131116 | ZNF428 | protein_coding |
| Huh7.5 | beta | ENSG00000250588 | IQCJ-SCHIP1 | protein_coding |
| Huh7.5 | beta | ENSG00000267296 | CEBPA-AS1 | antisense |
| Huh7.5 | beta | ENSG00000275457 | | antisense |
| Huh7.5 | beta | ENSG00000104903 | LYL1 | protein_coding |
| Huh7.5 | beta | ENSG00000247092 | SNHG10 | antisense |
| Huh7.5 | beta | ENSG00000167996 | FTH1 | protein_coding |
| Huh7.5 | beta | ENSG00000159674 | SPON2 | protein_coding |
| Huh7.5 | beta | ENSG00000275557 | | lincRNA |
| Huh7.5 | beta | ENSG00000165804 | ZNF219 | protein_coding |
| Huh7.5 | beta | ENSG00000159884 | CCDC107 | protein_coding |
| Huh7.5 | beta | ENSG00000171813 | PWWP2B | protein_coding |
| Huh7.5 | beta | ENSG00000152475 | ZNF837 | protein_coding |
| Huh7.5 | beta | ENSG00000130731 | C16orf13 | protein_coding |
| Huh7.5 | beta | ENSG00000141873 | SLC39A3 | protein_coding |
| Huh7.5 | beta | ENSG00000279873 | LINC01126 | TEC |
| Huh7.5 | beta | ENSG00000228981 | | processed_pseudogene |
| Huh7.5 | beta | ENSG00000232838 | PET117 | protein_coding |
| Huh7.5 | beta | ENSG00000099330 | OCEL1 | protein_coding |
| Huh7.5 | beta | ENSG00000261221 | ZNF865 | protein_coding |
| Huh7.5 | beta | ENSG00000114126 | TFDP2 | protein_coding |
| Huh7.5 | beta | ENSG00000105538 | RASIP1 | protein_coding |
| Huh7.5 | beta | ENSG00000165644 | COMTD1 | protein_coding |
| Huh7.5 | beta | ENSG00000006015 | C19orf60 | protein_coding |
| Huh7.5 | beta | ENSG00000213015 | ZNF580 | protein_coding |
| Huh7.5 | beta | ENSG00000243479 | MNX1-AS1 | lincRNA |
| Huh7.5 | beta | ENSG00000125652 | ALKBH7 | protein_coding |
| Huh7.5 | beta | ENSG00000280071 | | protein_coding |
| Huh7.5 | beta | ENSG00000167641 | PPP1R14A | protein_coding |
| Huh7.5 | beta | ENSG00000186603 | HPDL | protein_coding |
| Huh7.5 | beta | ENSG00000134874 | DZIP1 | protein_coding |
| Huh7.5 | beta | ENSG00000051596 | THOC3 | protein_coding |
| Huh7.5 | beta | ENSG00000247345 | | antisense |
| Huh7.5 | beta | ENSG00000124074 | ENKD1 | protein_coding |
| Huh7.5 | beta | ENSG00000167680 | SEMA6B | protein_coding |
| Huh7.5 | beta | ENSG00000178531 | CTXN1 | protein_coding |
| Huh7.5 | beta | ENSG00000143793 | C1orf35 | protein_coding |
| Huh7.5 | beta | ENSG00000127220 | ABHD8 | protein_coding |
| Huh7.5 | beta | ENSG00000263366 | | processed_pseudogene |
| Huh7.5 | beta | ENSG00000115257 | PCSK4 | protein_coding |
| Huh7.5 | beta | ENSG00000204618 | RNF39 | protein_coding |
| Huh7.5 | beta | ENSG00000135617 | PRADC1 | protein_coding |
| Huh7.5 | beta | ENSG00000279692 | | TEC |

| | | | | |
|---|---|---|---|---|
| Huh7.5 | beta | ENSG00000099953 | MMP11 | protein_coding |
| Huh7.5 | beta | ENSG00000269190 | FBXO17 | protein_coding |
| Huh7.5 | beta | ENSG00000169972 | PUSL1 | protein_coding |
| Huh7.5 | beta | ENSG00000065621 | GSTO2 | protein_coding |
| Huh7.5 | beta | ENSG00000183463 | URAD | protein_coding |
| Huh7.5 | beta | ENSG00000185522 | LMNTD2 | protein_coding |
| Huh7.5 | beta | ENSG00000126243 | LRFN3 | protein_coding |
| Huh7.5 | beta | ENSG00000182600 | C2orf82 | protein_coding |
| Huh7.5 | beta | ENSG00000196954 | CASP4 | protein_coding |
| Huh7.5 | beta | ENSG00000206341 | HLA-H | unprocessed_pseudogene |
| Huh7.5 | beta | ENSG00000172123 | SLFN12 | protein_coding |
| Huh7.5 | beta | ENSG00000221500 | SNORD100 | snoRNA |
| Huh7.5 | beta | ENSG00000151466 | SCLT1 | protein_coding |
| Huh7.5 | beta | ENSG00000196247 | ZNF107 | protein_coding |
| Huh7.5 | beta | ENSG00000245571 | | lincRNA |
| Huh7.5 | beta | ENSG00000257743 | | protein_coding |
| Huh7.5 | beta | ENSG00000213383 | | processed_pseudogene |
| Huh7.5 | beta | ENSG00000156966 | B3GNT7 | protein_coding |
| Huh7.5 | beta | ENSG00000225075 | | antisense |
| Huh7.5 | beta | ENSG00000240216 | CPHL1P | unitary_pseudogene |
| Huh7.5 | beta | ENSG00000164054 | SHISA5 | protein_coding |
| Huh7.5 | beta | ENSG00000120341 | SEC16B | protein_coding |
| Huh7.5 | beta | ENSG00000272512 | | lincRNA |
| Huh7.5 | beta | ENSG00000144820 | GPR128 | protein_coding |
| Huh7.5 | beta | ENSG00000183397 | C19orf71 | protein_coding |
| Huh7.5 | beta | ENSG00000114268 | PFKFB4 | protein_coding |
| Huh7.5 | beta | ENSG00000276171 | | miRNA |
| Huh7.5 | beta | ENSG00000151470 | C4orf33 | protein_coding |
| Huh7.5 | beta | ENSG00000280007 | | antisense |
| Huh7.5 | beta | ENSG00000255398 | HCAR3 | protein_coding |
| Huh7.5 | beta | ENSG00000132256 | TRIM5 | protein_coding |
| Huh7.5 | beta | ENSG00000243302 | | processed_pseudogene |
| Huh7.5 | beta | ENSG00000060491 | OGFR | protein_coding |
| Huh7.5 | beta | ENSG00000163823 | CCR1 | protein_coding |
| Huh7.5 | beta | ENSG00000173702 | MUC13 | protein_coding |
| Huh7.5 | beta | ENSG00000042980 | ADAM28 | protein_coding |
| Huh7.5 | beta | ENSG00000013374 | NUB1 | protein_coding |
| Huh7.5 | beta | ENSG00000227619 | | antisense |
| Huh7.5 | beta | ENSG00000185201 | IFITM2 | protein_coding |
| Huh7.5 | beta | ENSG00000104432 | IL7 | protein_coding |
| Huh7.5 | beta | ENSG00000166278 | C2 | protein_coding |
| Huh7.5 | beta | ENSG00000267745 | | processed_transcript |
| Huh7.5 | beta | ENSG00000113441 | LNPEP | protein_coding |
| Huh7.5 | beta | ENSG00000155158 | TTC39B | protein_coding |
| Huh7.5 | beta | ENSG00000271646 | | lincRNA |
| Huh7.5 | beta | ENSG00000279861 | | TEC |
| Huh7.5 | beta | ENSG00000134716 | CYP2J2 | protein_coding |
| Huh7.5 | beta | ENSG00000232498 | | antisense |
| Huh7.5 | beta | ENSG00000139192 | TAPBPL | protein_coding |
| Huh7.5 | beta | ENSG00000111912 | NCOA7 | protein_coding |
| Huh7.5 | beta | ENSG00000177721 | ANXA2R | protein_coding |
| Huh7.5 | beta | ENSG00000133943 | C14orf159 | protein_coding |
| Huh7.5 | beta | ENSG00000146859 | TMEM140 | protein_coding |
| Huh7.5 | beta | ENSG00000103056 | SMPD3 | protein_coding |
| Huh7.5 | beta | ENSG00000149131 | SERPING1 | protein_coding |
| Huh7.5 | beta | ENSG00000134470 | IL15RA | protein_coding |
| Huh7.5 | beta | ENSG00000125148 | MT2A | protein_coding |
| Huh7.5 | beta | ENSG00000243649 | CFB | protein_coding |
| Huh7.5 | beta | ENSG00000206503 | HLA-A | protein_coding |
| Huh7.5 | beta | ENSG00000117475 | BLZF1 | protein_coding |
| Huh7.5 | beta | ENSG00000157873 | TNFRSF14 | protein_coding |
| Huh7.5 | beta | ENSG00000275302 | CCL4 | protein_coding |

| | | | | |
|---|---|---|---|---|
| Huh7.5 | beta | ENSG00000269728 | | lincRNA |
| Huh7.5 | beta | ENSG00000123992 | DNPEP | protein_coding |
| Huh7.5 | beta | ENSG00000271550 | BNIP3P11 | processed_pseudogene |
| Huh7.5 | beta | ENSG00000196664 | TLR7 | protein_coding |
| Huh7.5 | beta | ENSG00000272821 | | antisense |
| Huh7.5 | beta | ENSG00000225131 | PSME2P2 | processed_pseudogene |
| Huh7.5 | beta | ENSG00000100911 | PSME2 | protein_coding |
| Huh7.5 | beta | ENSG00000231925 | TAPBP | protein_coding |
| Huh7.5 | beta | ENSG00000130487 | KLHDC7B | protein_coding |
| Huh7.5 | beta | ENSG00000137767 | SQRDL | protein_coding |
| Huh7.5 | beta | ENSG00000136147 | PHF11 | protein_coding |
| Huh7.5 | beta | ENSG00000114450 | GNB4 | protein_coding |
| Huh7.5 | beta | ENSG00000260302 | | lincRNA |
| Huh7.5 | beta | ENSG00000164136 | IL15 | protein_coding |
| Huh7.5 | beta | ENSG00000164307 | ERAP1 | protein_coding |
| Huh7.5 | beta | ENSG00000169894 | MUC3A | protein_coding |
| Huh7.5 | beta | ENSG00000064012 | CASP8 | protein_coding |
| Huh7.5 | beta | ENSG00000163644 | PPM1K | protein_coding |
| Huh7.5 | beta | ENSG00000155363 | MOV10 | protein_coding |
| Huh7.5 | beta | ENSG00000151025 | GPR158 | protein_coding |
| Huh7.5 | beta | ENSG00000276980 | | sense_intronic |
| Huh7.5 | beta | ENSG00000078018 | MAP2 | protein_coding |
| Huh7.5 | beta | ENSG00000079385 | CEACAM1 | protein_coding |
| Huh7.5 | beta | ENSG00000173786 | CNP | protein_coding |
| Huh7.5 | beta | ENSG00000242539 | | antisense |
| Huh7.5 | beta | ENSG00000137842 | TMEM62 | protein_coding |
| Huh7.5 | beta | ENSG00000107593 | PKD2L1 | protein_coding |
| Huh7.5 | beta | ENSG00000105402 | NAPA | protein_coding |
| Huh7.5 | beta | ENSG00000140511 | HAPLN3 | protein_coding |
| Huh7.5 | beta | ENSG00000152689 | RASGRP3 | protein_coding |
| Huh7.5 | beta | ENSG00000105287 | PRKD2 | protein_coding |
| Huh7.5 | beta | ENSG00000205220 | PSMB10 | protein_coding |
| Huh7.5 | beta | ENSG00000185404 | SP140L | protein_coding |
| Huh7.5 | beta | ENSG00000165806 | CASP7 | protein_coding |
| Huh7.5 | beta | ENSG00000137200 | CMTR1 | protein_coding |
| Huh7.5 | beta | ENSG00000177989 | ODF3B | protein_coding |
| Huh7.5 | beta | ENSG00000111801 | BTN3A3 | protein_coding |
| Huh7.5 | beta | ENSG00000182326 | C1S | protein_coding |
| Huh7.5 | beta | ENSG00000092010 | PSME1 | protein_coding |
| Huh7.5 | beta | ENSG00000213500 | LAP3P2 | processed_pseudogene |
| Huh7.5 | beta | ENSG00000238000 | | processed_pseudogene |
| Huh7.5 | beta | ENSG00000229241 | PNPT1P1 | processed_pseudogene |
| Huh7.5 | beta | ENSG00000112343 | TRIM38 | protein_coding |
| Huh7.5 | beta | ENSG00000072858 | SIDT1 | protein_coding |
| Huh7.5 | beta | ENSG00000160710 | ADAR | protein_coding |
| Huh7.5 | beta | ENSG00000168297 | PXK | protein_coding |
| Huh7.5 | beta | ENSG00000186470 | BTN3A2 | protein_coding |
| Huh7.5 | beta | ENSG00000128394 | APOBEC3F | protein_coding |
| Huh7.5 | beta | ENSG00000105939 | ZC3HAV1 | protein_coding |
| Huh7.5 | beta | ENSG00000142961 | MOB3C | protein_coding |
| Huh7.5 | beta | ENSG00000172936 | MYD88 | protein_coding |
| Huh7.5 | beta | ENSG00000166710 | B2M | protein_coding |
| Huh7.5 | beta | ENSG00000155629 | PIK3AP1 | protein_coding |
| Huh7.5 | beta | ENSG00000168016 | TRANK1 | protein_coding |
| Huh7.5 | beta | ENSG00000197536 | C5orf56 | protein_coding |
| Huh7.5 | beta | ENSG00000142677 | IL22RA1 | protein_coding |
| Huh7.5 | beta | ENSG00000108771 | DHX58 | protein_coding |
| Huh7.5 | beta | ENSG00000204397 | CARD16 | protein_coding |
| Huh7.5 | beta | ENSG00000230795 | HLA-K | unprocessed_pseudogene |
| Huh7.5 | beta | ENSG00000105559 | PLEKHA4 | protein_coding |
| Huh7.5 | beta | ENSG00000128335 | APOL2 | protein_coding |
| Huh7.5 | beta | ENSG00000116663 | FBXO6 | protein_coding |

| | | | | |
|---|---|---|---|---|
| Huh7.5 | beta | ENSG00000026950 | BTN3A1 | protein_coding |
| Huh7.5 | beta | ENSG00000225963 | | antisense |
| Huh7.5 | beta | ENSG00000184898 | RBM43 | protein_coding |
| Huh7.5 | beta | ENSG00000229474 | PATL2 | protein_coding |
| Huh7.5 | beta | ENSG00000259529 | IRF9 | protein_coding |
| Huh7.5 | beta | ENSG00000214872 | SMTNL1 | protein_coding |
| Huh7.5 | beta | ENSG00000196116 | TDRD7 | protein_coding |
| Huh7.5 | beta | ENSG00000243650 | RN7SL834P | misc_RNA |
| Huh7.5 | beta | ENSG00000124201 | ZNFX1 | protein_coding |
| Huh7.5 | beta | ENSG00000059378 | PARP12 | protein_coding |
| Huh7.5 | beta | ENSG00000173821 | RNF213 | protein_coding |
| Huh7.5 | beta | ENSG00000002549 | LAP3 | protein_coding |
| Huh7.5 | beta | ENSG00000106785 | TRIM14 | protein_coding |
| Huh7.5 | beta | ENSG00000140464 | PML | protein_coding |
| Huh7.5 | beta | ENSG00000137752 | CASP1 | protein_coding |
| Huh7.5 | beta | ENSG00000138035 | PNPT1 | protein_coding |
| Huh7.5 | beta | ENSG00000228318 | | antisense |
| Huh7.5 | beta | ENSG00000168404 | MLKL | protein_coding |
| Huh7.5 | beta | ENSG00000133321 | RARRES3 | protein_coding |
| Huh7.5 | beta | ENSG00000010030 | ETV7 | protein_coding |
| Huh7.5 | beta | ENSG00000123609 | NMI | protein_coding |
| Huh7.5 | beta | ENSG00000185880 | TRIM69 | protein_coding |
| Huh7.5 | beta | ENSG00000120217 | CD274 | protein_coding |
| Huh7.5 | beta | ENSG00000100918 | REC8 | protein_coding |
| Huh7.5 | beta | ENSG00000144596 | GRIP2 | protein_coding |
| Huh7.5 | beta | ENSG00000136514 | RTP4 | protein_coding |
| Huh7.5 | beta | ENSG00000164308 | ERAP2 | protein_coding |
| Huh7.5 | beta | ENSG00000159403 | C1R | protein_coding |
| Huh7.5 | beta | ENSG00000101347 | SAMHD1 | protein_coding |
| Huh7.5 | beta | ENSG00000169248 | CXCL11 | protein_coding |
| Huh7.5 | beta | ENSG00000185885 | IFITM1 | protein_coding |
| Huh7.5 | beta | ENSG00000172183 | ISG20 | protein_coding |
| Huh7.5 | beta | ENSG00000213928 | IRF9 | protein_coding |
| Huh7.5 | beta | ENSG00000121858 | TNFSF10 | protein_coding |
| Huh7.5 | beta | ENSG00000170581 | STAT2 | protein_coding |
| Huh7.5 | beta | ENSG00000121060 | TRIM25 | protein_coding |
| Huh7.5 | beta | ENSG00000204592 | HLA-E | protein_coding |
| Huh7.5 | beta | ENSG00000068079 | IFI35 | protein_coding |
| Huh7.5 | beta | ENSG00000132109 | TRIM21 | protein_coding |
| Huh7.5 | beta | ENSG00000164342 | TLR3 | protein_coding |
| Huh7.5 | beta | ENSG00000234745 | HLA-B | protein_coding |
| Huh7.5 | beta | ENSG00000115415 | STAT1 | protein_coding |
| Huh7.5 | beta | ENSG00000204525 | HLA-C | protein_coding |
| Huh7.5 | beta | ENSG00000165949 | IFI27 | protein_coding |
| Huh7.5 | beta | ENSG00000055332 | EIF2AK2 | protein_coding |
| Huh7.5 | beta | ENSG00000130589 | HELZ2 | protein_coding |
| Huh7.5 | beta | ENSG00000130813 | C19orf66 | protein_coding |
| Huh7.5 | beta | ENSG00000067066 | SP100 | protein_coding |
| Huh7.5 | beta | ENSG00000117226 | GBP3 | protein_coding |
| Huh7.5 | beta | ENSG00000188313 | PLSCR1 | protein_coding |
| Huh7.5 | beta | ENSG00000163840 | DTX3L | protein_coding |
| Huh7.5 | beta | ENSG00000269640 | | lincRNA |
| Huh7.5 | beta | ENSG00000204267 | TAP2 | protein_coding |
| Huh7.5 | beta | ENSG00000225886 | | antisense |
| Huh7.5 | beta | ENSG00000184979 | USP18 | protein_coding |
| Huh7.5 | beta | ENSG00000204264 | PSMB8 | protein_coding |
| Huh7.5 | beta | ENSG00000168062 | BATF2 | protein_coding |
| Huh7.5 | beta | ENSG00000177409 | SAMD9L | protein_coding |
| Huh7.5 | beta | ENSG00000178685 | PARP10 | protein_coding |
| Huh7.5 | beta | ENSG00000169245 | CXCL10 | protein_coding |
| Huh7.5 | beta | ENSG00000107201 | DDX58 | protein_coding |
| Huh7.5 | beta | ENSG00000100342 | APOL1 | protein_coding |

| | | | | |
|---|---|---|---|---|
| Huh7.5 | beta | ENSG00000078081 | LAMP3 | protein_coding |
| Huh7.5 | beta | ENSG00000138646 | HERC5 | protein_coding |
| Huh7.5 | beta | ENSG00000152778 | IFIT5 | protein_coding |
| Huh7.5 | beta | ENSG00000182179 | UBA7 | protein_coding |
| Huh7.5 | beta | ENSG00000240065 | PSMB9 | protein_coding |
| Huh7.5 | beta | ENSG00000138496 | PARP9 | protein_coding |
| Huh7.5 | beta | ENSG00000181381 | DDX60L | protein_coding |
| Huh7.5 | beta | ENSG00000117228 | GBP1 | protein_coding |
| Huh7.5 | beta | ENSG00000132274 | TRIM22 | protein_coding |
| Huh7.5 | beta | ENSG00000135899 | SP110 | protein_coding |
| Huh7.5 | beta | ENSG00000108679 | LGALS3BP | protein_coding |
| Huh7.5 | beta | ENSG00000156587 | UBE2L6 | protein_coding |
| Huh7.5 | beta | ENSG00000140853 | NLRC5 | protein_coding |
| Huh7.5 | beta | ENSG00000132530 | XAF1 | protein_coding |
| Huh7.5 | beta | ENSG00000142089 | IFITM3 | protein_coding |
| Huh7.5 | beta | ENSG00000173193 | PARP14 | protein_coding |
| Huh7.5 | beta | ENSG00000133106 | EPSTI1 | protein_coding |
| Huh7.5 | beta | ENSG00000110446 | SLC15A3 | protein_coding |
| Huh7.5 | beta | ENSG00000137965 | IFI44 | protein_coding |
| Huh7.5 | beta | ENSG00000119922 | IFIT2 | protein_coding |
| Huh7.5 | beta | ENSG00000135114 | OASL | protein_coding |
| Huh7.5 | beta | ENSG00000221963 | APOL6 | protein_coding |
| Huh7.5 | beta | ENSG00000187608 | ISG15 | protein_coding |
| Huh7.5 | beta | ENSG00000205413 | SAMD9 | protein_coding |
| Huh7.5 | beta | ENSG00000168394 | TAP1 | protein_coding |
| Huh7.5 | beta | ENSG00000138642 | HERC6 | protein_coding |
| Huh7.5 | beta | ENSG00000130303 | BST2 | protein_coding |
| Huh7.5 | beta | ENSG00000134326 | CMPK2 | protein_coding |
| Huh7.5 | beta | ENSG00000137628 | DDX60 | protein_coding |
| Huh7.5 | beta | ENSG00000111331 | OAS3 | protein_coding |
| Huh7.5 | beta | ENSG00000119917 | IFIT3 | protein_coding |
| Huh7.5 | beta | ENSG00000137959 | IFI44L | protein_coding |
| Huh7.5 | beta | ENSG00000115267 | IFIH1 | protein_coding |
| Huh7.5 | beta | ENSG00000134321 | RSAD2 | protein_coding |
| Huh7.5 | beta | ENSG00000089127 | OAS1 | protein_coding |
| Huh7.5 | beta | ENSG00000126709 | IFI6 | protein_coding |
| Huh7.5 | beta | ENSG00000111335 | OAS2 | protein_coding |
| Huh7.5 | beta | ENSG00000185745 | IFIT1 | protein_coding |
| Huh7.5 | beta | ENSG00000157601 | MX1 | protein_coding |
| Huh7.5 | lambda | ENSG00000157601 | MX1 | protein_coding |
| Huh7.5 | lambda | ENSG00000126709 | IFI6 | protein_coding |
| Huh7.5 | lambda | ENSG00000185745 | IFIT1 | protein_coding |
| Huh7.5 | lambda | ENSG00000115267 | IFIH1 | protein_coding |
| Huh7.5 | lambda | ENSG00000111331 | OAS3 | protein_coding |
| Huh7.5 | lambda | ENSG00000137628 | DDX60 | protein_coding |
| Huh7.5 | lambda | ENSG00000119917 | IFIT3 | protein_coding |
| Huh7.5 | lambda | ENSG00000134326 | CMPK2 | protein_coding |
| Huh7.5 | lambda | ENSG00000187608 | ISG15 | protein_coding |
| Huh7.5 | lambda | ENSG00000089127 | OAS1 | protein_coding |
| Huh7.5 | lambda | ENSG00000138642 | HERC6 | protein_coding |
| Huh7.5 | lambda | ENSG00000138496 | PARP9 | protein_coding |
| Huh7.5 | lambda | ENSG00000137959 | IFI44L | protein_coding |
| Huh7.5 | lambda | ENSG00000225886 | | antisense |
| Huh7.5 | lambda | ENSG00000111335 | OAS2 | protein_coding |
| Huh7.5 | lambda | ENSG00000205413 | SAMD9 | protein_coding |
| Huh7.5 | lambda | ENSG00000133106 | EPSTI1 | protein_coding |
| Huh7.5 | lambda | ENSG00000173193 | PARP14 | protein_coding |
| Huh7.5 | lambda | ENSG00000130303 | BST2 | protein_coding |
| Huh7.5 | lambda | ENSG00000221963 | APOL6 | protein_coding |
| Huh7.5 | lambda | ENSG00000142089 | IFITM3 | protein_coding |
| Huh7.5 | lambda | ENSG00000135899 | SP110 | protein_coding |
| Huh7.5 | lambda | ENSG00000168394 | TAP1 | protein_coding |

| | | | | |
|---|---|---|---|---|
| Huh7.5 | lambda | ENSG00000152778 | IFIT5 | protein_coding |
| Huh7.5 | lambda | ENSG00000107201 | DDX58 | protein_coding |
| Huh7.5 | lambda | ENSG00000181381 | DDX60L | protein_coding |
| Huh7.5 | lambda | ENSG00000134321 | RSAD2 | protein_coding |
| Huh7.5 | lambda | ENSG00000163840 | DTX3L | protein_coding |
| Huh7.5 | lambda | ENSG00000130589 | HELZ2 | protein_coding |
| Huh7.5 | lambda | ENSG00000055332 | EIF2AK2 | protein_coding |
| Huh7.5 | lambda | ENSG00000108679 | LGALS3BP | protein_coding |
| Huh7.5 | lambda | ENSG00000188313 | PLSCR1 | protein_coding |
| Huh7.5 | lambda | ENSG00000213928 | IRF9 | protein_coding |
| Huh7.5 | lambda | ENSG00000119922 | IFIT2 | protein_coding |
| Huh7.5 | lambda | ENSG00000115415 | STAT1 | protein_coding |
| Huh7.5 | lambda | ENSG00000178685 | PARP10 | protein_coding |
| Huh7.5 | lambda | ENSG00000135114 | OASL | protein_coding |
| Huh7.5 | lambda | ENSG00000100918 | REC8 | protein_coding |
| Huh7.5 | lambda | ENSG00000117228 | GBP1 | protein_coding |
| Huh7.5 | lambda | ENSG00000184979 | USP18 | protein_coding |
| Huh7.5 | lambda | ENSG00000067066 | SP100 | protein_coding |
| Huh7.5 | lambda | ENSG00000138646 | HERC5 | protein_coding |
| Huh7.5 | lambda | ENSG00000110446 | SLC15A3 | protein_coding |
| Huh7.5 | lambda | ENSG00000132530 | XAF1 | protein_coding |
| Huh7.5 | lambda | ENSG00000168062 | BATF2 | protein_coding |
| Huh7.5 | lambda | ENSG00000121060 | TRIM25 | protein_coding |
| Huh7.5 | lambda | ENSG00000132109 | TRIM21 | protein_coding |
| Huh7.5 | lambda | ENSG00000156587 | UBE2L6 | protein_coding |
| Huh7.5 | lambda | ENSG00000228318 | | antisense |
| Huh7.5 | lambda | ENSG00000259529 | IRF9 | protein_coding |
| Huh7.5 | lambda | ENSG00000106785 | TRIM14 | protein_coding |
| Huh7.5 | lambda | ENSG00000169245 | CXCL10 | protein_coding |
| Huh7.5 | lambda | ENSG00000204267 | TAP2 | protein_coding |
| Huh7.5 | lambda | ENSG00000105939 | ZC3HAV1 | protein_coding |
| Huh7.5 | lambda | ENSG00000059378 | PARP12 | protein_coding |
| Huh7.5 | lambda | ENSG00000269640 | | lincRNA |
| Huh7.5 | lambda | ENSG00000243650 | RN7SL834P | misc_RNA |
| Huh7.5 | lambda | ENSG00000185880 | TRIM69 | protein_coding |
| Huh7.5 | lambda | ENSG00000137965 | IFI44 | protein_coding |
| Huh7.5 | lambda | ENSG00000140464 | PML | protein_coding |
| Huh7.5 | lambda | ENSG00000117226 | GBP3 | protein_coding |
| Huh7.5 | lambda | ENSG00000130813 | C19orf66 | protein_coding |
| Huh7.5 | lambda | ENSG00000170581 | STAT2 | protein_coding |
| Huh7.5 | lambda | ENSG00000140853 | NLRC5 | protein_coding |
| Huh7.5 | lambda | ENSG00000204525 | HLA-C | protein_coding |
| Huh7.5 | lambda | ENSG00000124201 | ZNFX1 | protein_coding |
| Huh7.5 | lambda | ENSG00000196116 | TDRD7 | protein_coding |
| Huh7.5 | lambda | ENSG00000177989 | ODF3B | protein_coding |
| Huh7.5 | lambda | ENSG00000225963 | | antisense |
| Huh7.5 | lambda | ENSG00000164308 | ERAP2 | protein_coding |
| Huh7.5 | lambda | ENSG00000184898 | RBM43 | protein_coding |
| Huh7.5 | lambda | ENSG00000160710 | ADAR | protein_coding |
| Huh7.5 | lambda | ENSG00000136147 | PHF11 | protein_coding |
| Huh7.5 | lambda | ENSG00000177409 | SAMD9L | protein_coding |
| Huh7.5 | lambda | ENSG00000182179 | UBA7 | protein_coding |
| Huh7.5 | lambda | ENSG00000138035 | PNPT1 | protein_coding |
| Huh7.5 | lambda | ENSG00000002549 | LAP3 | protein_coding |
| Huh7.5 | lambda | ENSG00000123609 | NMI | protein_coding |
