## Supplemental File 2 for "Novel antiviral interferon sensitive genes unveiled by correlation-driven gene selection and integrated systems biology approaches"

| ensemble_gene_id | module |
|---|---|
| ENSG00000002549 | Green |
| ENSG00000006062 | Green |
| ENSG00000008735 | Green |
| ENSG00000010030 | Green |
| ENSG00000013374 | Green |
| ENSG00000025708 | Green |
| ENSG00000026950 | Green |
| ENSG00000027075 | Green |
| ENSG00000049192 | Green |
| ENSG00000055332 | Green |
| ENSG00000059378 | Green |
| ENSG00000060491 | Green |
| ENSG00000060718 | Green |
| ENSG00000064012 | Green |
| ENSG00000067066 | Green |
| ENSG00000068079 | Green |
| ENSG00000072858 | Green |
| ENSG00000073605 | Green |
| ENSG00000078018 | Green |
| ENSG00000078081 | Green |
| ENSG00000079385 | Green |
| ENSG00000087085 | Green |
| ENSG00000088881 | Green |
| ENSG00000089127 | Green |
| ENSG00000089199 | Green |
| ENSG00000091138 | Green |
| ENSG00000092010 | Green |
| ENSG00000092098 | Green |
| ENSG00000100226 | Green |
| ENSG00000100342 | Green |
| ENSG00000100505 | Green |
| ENSG00000100911 | Green |
| ENSG00000100918 | Green |
| ENSG00000101347 | Green |
| ENSG00000103044 | Green |
| ENSG00000103056 | Green |
| ENSG00000104213 | Green |
| ENSG00000104320 | Green |
| ENSG00000104518 | Green |
| ENSG00000104689 | Green |
| ENSG00000105287 | Green |
| ENSG00000105402 | Green |
| ENSG00000105559 | Green |
| ENSG00000105835 | Green |
| ENSG00000105939 | Green |
| ENSG00000106100 | Green |
| ENSG00000106785 | Green |
| ENSG00000106829 | Green |
| ENSG00000106948 | Green |

| | |
|---|---|
| ENSG00000107201 | Green |
| ENSG00000107593 | Green |
| ENSG00000108669 | Green |
| ENSG00000108679 | Green |
| ENSG00000108771 | Green |
| ENSG00000110057 | Green |
| ENSG00000110446 | Green |
| ENSG00000111012 | Green |
| ENSG00000111331 | Green |
| ENSG00000111335 | Green |
| ENSG00000111801 | Green |
| ENSG00000111912 | Green |
| ENSG00000112343 | Green |
| ENSG00000113441 | Green |
| ENSG00000114127 | Green |
| ENSG00000114268 | Green |
| ENSG00000115267 | Green |
| ENSG00000115415 | Green |
| ENSG00000116663 | Green |
| ENSG00000117226 | Green |
| ENSG00000117228 | Green |
| ENSG00000117266 | Green |
| ENSG00000117475 | Green |
| ENSG00000118507 | Green |
| ENSG00000119917 | Green |
| ENSG00000119922 | Green |
| ENSG00000120217 | Green |
| ENSG00000120341 | Green |
| ENSG00000120539 | Green |
| ENSG00000120549 | Green |
| ENSG00000120738 | Green |
| ENSG00000121060 | Green |
| ENSG00000121380 | Green |
| ENSG00000121769 | Green |
| ENSG00000121858 | Green |
| ENSG00000122176 | Green |
| ENSG00000122643 | Green |
| ENSG00000123240 | Green |
| ENSG00000123609 | Green |
| ENSG00000123685 | Green |
| ENSG00000123992 | Green |
| ENSG00000124201 | Green |
| ENSG00000124613 | Green |
| ENSG00000125148 | Green |
| ENSG00000125347 | Green |
| ENSG00000125826 | Green |
| ENSG00000126709 | Green |
| ENSG00000126778 | Green |
| ENSG00000127666 | Green |
| ENSG00000128335 | Green |

| | |
|---|---|
| ENSG00000128394 | Green |
| ENSG00000128567 | Green |
| ENSG00000130303 | Green |
| ENSG00000130487 | Green |
| ENSG00000130589 | Green |
| ENSG00000130813 | Green |
| ENSG00000131797 | Green |
| ENSG00000132109 | Green |
| ENSG00000132256 | Green |
| ENSG00000132274 | Green |
| ENSG00000132530 | Green |
| ENSG00000132744 | Green |
| ENSG00000133106 | Green |
| ENSG00000133321 | Green |
| ENSG00000133943 | Green |
| ENSG00000134321 | Green |
| ENSG00000134326 | Green |
| ENSG00000134470 | Green |
| ENSG00000134716 | Green |
| ENSG00000135114 | Green |
| ENSG00000135148 | Green |
| ENSG00000135899 | Green |
| ENSG00000136147 | Green |
| ENSG00000136514 | Green |
| ENSG00000136874 | Green |
| ENSG00000137200 | Green |
| ENSG00000137449 | Green |
| ENSG00000137628 | Green |
| ENSG00000137752 | Green |
| ENSG00000137767 | Green |
| ENSG00000137842 | Green |
| ENSG00000137959 | Green |
| ENSG00000137965 | Green |
| ENSG00000138035 | Green |
| ENSG00000138496 | Green |
| ENSG00000138642 | Green |
| ENSG00000138646 | Green |
| ENSG00000139083 | Green |
| ENSG00000139180 | Green |
| ENSG00000139192 | Green |
| ENSG00000140464 | Green |
| ENSG00000140511 | Green |
| ENSG00000140832 | Green |
| ENSG00000140853 | Green |
| ENSG00000141497 | Green |
| ENSG00000141664 | Green |
| ENSG00000142089 | Green |
| ENSG00000142677 | Green |
| ENSG00000142961 | Green |
| ENSG00000143156 | Green |

| | |
|---|---|
| ENSG00000143390 | Green |
| ENSG00000143507 | Green |
| ENSG00000144035 | Green |
| ENSG00000144596 | Green |
| ENSG00000144852 | Green |
| ENSG00000145623 | Green |
| ENSG00000146859 | Green |
| ENSG00000149131 | Green |
| ENSG00000149289 | Green |
| ENSG00000151025 | Green |
| ENSG00000151466 | Green |
| ENSG00000151470 | Green |
| ENSG00000152689 | Green |
| ENSG00000152778 | Green |
| ENSG00000153898 | Green |
| ENSG00000153933 | Green |
| ENSG00000155158 | Green |
| ENSG00000155287 | Green |
| ENSG00000155363 | Green |
| ENSG00000155629 | Green |
| ENSG00000156232 | Green |
| ENSG00000156500 | Green |
| ENSG00000156587 | Green |
| ENSG00000156966 | Green |
| ENSG00000157303 | Green |
| ENSG00000157601 | Green |
| ENSG00000157873 | Green |
| ENSG00000159403 | Green |
| ENSG00000160710 | Green |
| ENSG00000161091 | Green |
| ENSG00000162654 | Green |
| ENSG00000162687 | Green |
| ENSG00000162772 | Green |
| ENSG00000163121 | Green |
| ENSG00000163131 | Green |
| ENSG00000163644 | Green |
| ENSG00000163661 | Green |
| ENSG00000163702 | Green |
| ENSG00000163823 | Green |
| ENSG00000163840 | Green |
| ENSG00000164136 | Green |
| ENSG00000164181 | Green |
| ENSG00000164211 | Green |
| ENSG00000164307 | Green |
| ENSG00000164308 | Green |
| ENSG00000164342 | Green |
| ENSG00000165806 | Green |
| ENSG00000165949 | Green |
| ENSG00000166278 | Green |
| ENSG00000166689 | Green |

| | |
|---|---|
| ENSG00000166710 | Green |
| ENSG00000167705 | Green |
| ENSG00000168016 | Green |
| ENSG00000168062 | Green |
| ENSG00000168297 | Green |
| ENSG00000168310 | Green |
| ENSG00000168394 | Green |
| ENSG00000168404 | Green |
| ENSG00000168899 | Green |
| ENSG00000169245 | Green |
| ENSG00000169248 | Green |
| ENSG00000169429 | Green |
| ENSG00000169871 | Green |
| ENSG00000169894 | Green |
| ENSG00000170006 | Green |
| ENSG00000170345 | Green |
| ENSG00000170581 | Green |
| ENSG00000171132 | Green |
| ENSG00000171310 | Green |
| ENSG00000171649 | Green |
| ENSG00000172183 | Green |
| ENSG00000172936 | Green |
| ENSG00000173193 | Green |
| ENSG00000173221 | Green |
| ENSG00000173702 | Green |
| ENSG00000173786 | Green |
| ENSG00000173821 | Green |
| ENSG00000174808 | Green |
| ENSG00000174992 | Green |
| ENSG00000177409 | Green |
| ENSG00000177721 | Green |
| ENSG00000177989 | Green |
| ENSG00000178685 | Green |
| ENSG00000178719 | Green |
| ENSG00000179627 | Green |
| ENSG00000179674 | Green |
| ENSG00000180628 | Green |
| ENSG00000181284 | Green |
| ENSG00000181381 | Green |
| ENSG00000182179 | Green |
| ENSG00000182326 | Green |
| ENSG00000183397 | Green |
| ENSG00000183625 | Green |
| ENSG00000184371 | Green |
| ENSG00000184898 | Green |
| ENSG00000184979 | Green |
| ENSG00000185201 | Green |
| ENSG00000185404 | Green |
| ENSG00000185745 | Green |
| ENSG00000185880 | Green |

| | |
|---|---|
| ENSG00000185885 | Green |
| ENSG00000186470 | Green |
| ENSG00000186994 | Green |
| ENSG00000187608 | Green |
| ENSG00000188290 | Green |
| ENSG00000188313 | Green |
| ENSG00000189366 | Green |
| ENSG00000196116 | Green |
| ENSG00000196247 | Green |
| ENSG00000196954 | Green |
| ENSG00000197019 | Green |
| ENSG00000197142 | Green |
| ENSG00000197536 | Green |
| ENSG00000197808 | Green |
| ENSG00000198959 | Green |
| ENSG00000204264 | Green |
| ENSG00000204267 | Green |
| ENSG00000204397 | Green |
| ENSG00000204525 | Green |
| ENSG00000204592 | Green |
| ENSG00000205220 | Green |
| ENSG00000205413 | Green |
| ENSG00000206341 | Green |
| ENSG00000206503 | Green |
| ENSG00000213492 | Green |
| ENSG00000213500 | Green |
| ENSG00000213928 | Green |
| ENSG00000214013 | Green |
| ENSG00000214226 | Green |
| ENSG00000214872 | Green |
| ENSG00000215068 | Green |
| ENSG00000215284 | Green |
| ENSG00000217801 | Green |
| ENSG00000221963 | Green |
| ENSG00000223960 | Green |
| ENSG00000225075 | Green |
| ENSG00000225131 | Green |
| ENSG00000225377 | Green |
| ENSG00000225492 | Green |
| ENSG00000225886 | Green |
| ENSG00000225963 | Green |
| ENSG00000226161 | Green |
| ENSG00000228318 | Green |
| ENSG00000228612 | Green |
| ENSG00000229241 | Green |
| ENSG00000229474 | Green |
| ENSG00000229644 | Green |
| ENSG00000229873 | Green |
| ENSG00000230715 | Green |
| ENSG00000230795 | Green |

| | |
|---|---|
| ENSG00000231574 | Green |
| ENSG00000231925 | Green |
| ENSG00000232498 | Green |
| ENSG00000233593 | Green |
| ENSG00000234127 | Green |
| ENSG00000234745 | Green |
| ENSG00000238000 | Green |
| ENSG00000238018 | Green |
| ENSG00000240065 | Green |
| ENSG00000240216 | Green |
| ENSG00000243302 | Green |
| ENSG00000243649 | Green |
| ENSG00000243650 | Green |
| ENSG00000245556 | Green |
| ENSG00000245571 | Green |
| ENSG00000248988 | Green |
| ENSG00000250155 | Green |
| ENSG00000254454 | Green |
| ENSG00000254995 | Green |
| ENSG00000255031 | Green |
| ENSG00000257743 | Green |
| ENSG00000258733 | Green |
| ENSG00000259529 | Green |
| ENSG00000260267 | Green |
| ENSG00000260302 | Green |
| ENSG00000261884 | Green |
| ENSG00000261971 | Green |
| ENSG00000263069 | Green |
| ENSG00000263528 | Green |
| ENSG00000266074 | Green |
| ENSG00000267102 | Green |
| ENSG00000267374 | Green |
| ENSG00000267547 | Green |
| ENSG00000267648 | Green |
| ENSG00000267745 | Green |
| ENSG00000268001 | Green |
| ENSG00000268205 | Green |
| ENSG00000269640 | Green |
| ENSG00000271550 | Green |
| ENSG00000271646 | Green |
| ENSG00000272512 | Green |
| ENSG00000272669 | Green |
| ENSG00000272821 | Green |
| ENSG00000272886 | Green |
| ENSG00000279400 | Green |
| ENSG00000279861 | Green |
| ENSG00000280007 | Green |
| ENSG00000000971 | Tan |
| ENSG00000019582 | Tan |
| ENSG00000023445 | Tan |

| | |
|---|---|
| ENSG00000067221 | Tan |
| ENSG00000070759 | Tan |
| ENSG00000081041 | Tan |
| ENSG00000081803 | Tan |
| ENSG00000083817 | Tan |
| ENSG00000093134 | Tan |
| ENSG00000095380 | Tan |
| ENSG00000100889 | Tan |
| ENSG00000104432 | Tan |
| ENSG00000107798 | Tan |
| ENSG00000112096 | Tan |
| ENSG00000115009 | Tan |
| ENSG00000115271 | Tan |
| ENSG00000118322 | Tan |
| ENSG00000118503 | Tan |
| ENSG00000121236 | Tan |
| ENSG00000122641 | Tan |
| ENSG00000124102 | Tan |
| ENSG00000124406 | Tan |
| ENSG00000128655 | Tan |
| ENSG00000135378 | Tan |
| ENSG00000136783 | Tan |
| ENSG00000136826 | Tan |
| ENSG00000137877 | Tan |
| ENSG00000141655 | Tan |
| ENSG00000141682 | Tan |
| ENSG00000142102 | Tan |
| ENSG00000144802 | Tan |
| ENSG00000144820 | Tan |
| ENSG00000147168 | Tan |
| ENSG00000162542 | Tan |
| ENSG00000162645 | Tan |
| ENSG00000163734 | Tan |
| ENSG00000163739 | Tan |
| ENSG00000164761 | Tan |
| ENSG00000165923 | Tan |
| ENSG00000166016 | Tan |
| ENSG00000166801 | Tan |
| ENSG00000168497 | Tan |
| ENSG00000171428 | Tan |
| ENSG00000171729 | Tan |
| ENSG00000171840 | Tan |
| ENSG00000172123 | Tan |
| ENSG00000173930 | Tan |
| ENSG00000177738 | Tan |
| ENSG00000183508 | Tan |
| ENSG00000185022 | Tan |
| ENSG00000185215 | Tan |
| ENSG00000186198 | Tan |
| ENSG00000187583 | Tan |

| | |
|---|---|
| ENSG00000188641 | Tan |
| ENSG00000196664 | Tan |
| ENSG00000196776 | Tan |
| ENSG00000197408 | Tan |
| ENSG00000213886 | Tan |
| ENSG00000227403 | Tan |
| ENSG00000227619 | Tan |
| ENSG00000228203 | Tan |
| ENSG00000249740 | Tan |
| ENSG00000251637 | Tan |
| ENSG00000256043 | Tan |
| ENSG00000260336 | Tan |
