## Supplemental File 3 for "Novel antiviral interferon sensitive genes unveiled by correlation-driven gene selection and integrated systems biology approaches"

| Module | GeneName | EnsembleGeneID | GeneType | Closeness | Eigenvector | Peripheral | Classification |
|---|---|---|---|---|---|---|---|
| Green | LAP3 | ENSG00000002549 | protein_coding | 0.246841586 | 0.105976112 | No | IFN_related_manual |
| Green | MAP3K14 | ENSG00000006062 | protein_coding | 0.063567399 | 7.75E-06 | Yes | discarted_as_peripheral |
| Green | MAPK8IP2 | ENSG00000008735 | protein_coding | 0.108708995 | 5.14E-04 | Yes | discarted_as_peripheral |
| Green | ETV7 | ENSG00000010030 | protein_coding | 0.20023954 | 0.051188402 | No | IFN_related_manual |
| Green | NUB1 | ENSG00000013374 | protein_coding | 0.222412459 | 0.070422344 | No | IFN_related_automatic |
| Green | TYMP | ENSG00000025708 | protein_coding | 0.115364312 | 0.001082994 | Yes | discarted_as_peripheral |
| Green | BTN3A1 | ENSG00000026950 | protein_coding | 0.229844662 | 0.080825001 | No | IFN_related_manual |
| Green | ADAMTS6 | ENSG00000049192 | protein_coding | 0.137478172 | 0.004836451 | Yes | discarted_as_peripheral |
| Green | EIF2AK2 | ENSG00000055332 | protein_coding | 0.208009875 | 0.07343626 | No | IFN_related_automatic |
| Green | PARP12 | ENSG00000059378 | protein_coding | 0.229104869 | 0.087045796 | No | IFN_related_manual |
| Green | OGFR | ENSG00000060491 | protein_coding | 0.187139036 | 0.042584542 | No | IFN_related_manual |
| Green | CASP8 | ENSG00000064012 | protein_coding | 0.230036981 | 0.081883796 | No | IFN_related_automatic |
| Green | SP100 | ENSG00000067066 | protein_coding | 0.233803438 | 0.09798608 | No | IFN_related_automatic |
| Green | IFI35 | ENSG00000068079 | protein_coding | 0.185819884 | 0.037074834 | No | IFN_related_automatic |
| Green | SIDT1 | ENSG00000072858 | protein_coding | 0.145871515 | 0.020724807 | No | novel |
| Green | GSDMB | ENSG00000073605 | protein_coding | 0.122285589 | 0.005722254 | Yes | discarted_as_peripheral |
| Green | MAP2 | ENSG00000078018 | protein_coding | 0.214489458 | 0.066088051 | No | IFN_related_manual |
| Green | LAMP3 | ENSG00000078081 | protein_coding | 0.209922407 | 0.060754657 | No | IFN_related_manual |
| Green | CEACAM1 | ENSG00000079385 | protein_coding | 0.209926411 | 0.06362772 | No | IFN_related_manual |
| Green | ACHE | ENSG00000087085 | protein_coding | 0.115260768 | 0.001183534 | Yes | discarted_as_peripheral |
| Green | EBF4 | ENSG00000088881 | protein_coding | 0.126067785 | 0.001443363 | Yes | discarted_as_peripheral |
| Green | OAS1 | ENSG00000089127 | protein_coding | 0.239095971 | 0.101576693 | No | IFN_related_automatic |
| Green | PSME1 | ENSG00000092010 | protein_coding | 0.22181327 | 0.069015548 | No | IFN_related_automatic |
| Green | RNF31 | ENSG00000092098 | protein_coding | 0.151566617 | 0.013529049 | Yes | discarted_as_peripheral |
| Green | GTPBP1 | ENSG00000100226 | protein_coding | 0.224426736 | 0.072761461 | No | IFN_related_manual |
| Green | APOL1 | ENSG00000100342 | protein_coding | 0.229940733 | 0.082272567 | No | IFN_related_automatic |
| Green | PSME2 | ENSG00000100911 | protein_coding | 0.228863164 | 0.080346733 | No | IFN_related_automatic |
| Green | REC8 | ENSG00000100918 | protein_coding | 0.18159153 | 0.046024825 | No | IFN_related_manual |
| Green | SAMHD1 | ENSG00000101347 | protein_coding | 0.240869231 | 0.098996848 | No | IFN_related_automatic |
| Green | SMPD3 | ENSG00000103056 | protein_coding | 0.121254144 | 0.011339874 | Yes | discarted_as_peripheral |
| Green | NBN | ENSG00000104320 | protein_coding | 0.156000364 | 0.013236931 | Yes | discarted_as_peripheral |
| Green | GSDMD | ENSG00000104518 | protein_coding | 0.192100666 | 0.044307951 | No | IFN_related_manual |
| Green | TNFRSF10A | ENSG00000104689 | protein_coding | 0.083035846 | 2.10E-04 | Yes | discarted_as_peripheral |
| Green | PRKD2 | ENSG00000105287 | protein_coding | 0.243907311 | 0.096943781 | No | IFN_related_automatic |
| Green | NAPA | ENSG00000105402 | protein_coding | 0.189233304 | 0.041797813 | No | IFN_related_manual |
| Green | PLEKHA4 | ENSG00000105559 | protein_coding | 0.237724539 | 0.08267764 | No | IFN_related_manual |
| Green | NAMPT | ENSG00000105835 | protein_coding | 0.196218846 | 0.054490227 | No | IFN_related_manual |
| Green | ZC3HAV1 | ENSG00000105939 | protein_coding | 0.1894203 | 0.051778335 | No | IFN_related_automatic |
| Green | NOD1 | ENSG00000106100 | protein_coding | 0.074524943 | 1.69E-04 | Yes | discarted_as_peripheral |
| Green | TRIM14 | ENSG00000106785 | protein_coding | 0.202982495 | 0.059833523 | No | IFN_related_automatic |
| Green | TLE4 | ENSG00000106829 | protein_coding | 0.158204611 | 0.011396846 | Yes | discarted_as_peripheral |
| Green | AKNA | ENSG00000106948 | protein_coding | 0.061367388 | 3.29E-06 | Yes | discarted_as_peripheral |
| Green | DDX58 | ENSG00000107201 | protein_coding | 0.220371015 | 0.084601745 | No | IFN_related_automatic |
| Green | PKD2L1 | ENSG00000107593 | protein_coding | 0.080982622 | 0.001033569 | Yes | discarted_as_peripheral |
| Green | CYTH1 | ENSG00000108669 | protein_coding | 0.218883073 | 0.062759243 | No | novel |
| Green | LGALS3BP | ENSG00000108679 | protein_coding | 0.25102536 | 0.107848249 | No | IFN_related_manual |
| Green | DHX58 | ENSG00000108771 | protein_coding | 0.169303017 | 0.033511382 | No | IFN_related_automatic |
| Green | UNC93B1 | ENSG00000110057 | protein_coding | 0.152754133 | 0.010491636 | Yes | discarted_as_peripheral |
| Green | SLC15A3 | ENSG00000110446 | protein_coding | 0.243581374 | 0.101347752 | No | IFN_related_manual |
| Green | OAS3 | ENSG00000111331 | protein_coding | 0.214785669 | 0.079761893 | No | IFN_related_automatic |
| Green | OAS2 | ENSG00000111335 | protein_coding | 0.24974089 | 0.111163557 | No | IFN_related_automatic |
| Green | BTN3A3 | ENSG00000111801 | protein_coding | 0.168682531 | 0.03328459 | No | IFN_related_manual |
| Green | NCOA7 | ENSG00000111912 | protein_coding | 0.211511895 | 0.069745146 | No | IFN_related_manual |
| Green | TRIM38 | ENSG00000112343 | protein_coding | 0.126910055 | 0.007968246 | Yes | discarted_as_peripheral |
| Green | LNPEP | ENSG00000113441 | protein_coding | 0.211117922 | 0.074431002 | No | IFN_related_automatic |
| Green | XRN1 | ENSG00000114127 | protein_coding | 0.171768444 | 0.035393417 | No | IFN_related_manual |
| Green | PFKFB4 | ENSG00000114268 | protein_coding | 0.143265084 | 0.008351543 | Yes | discarted_as_peripheral |
| Green | IFIH1 | ENSG00000115267 | protein_coding | 0.216156157 | 0.081865668 | No | IFN_related_automatic |
| Green | STAT1 | ENSG00000115415 | protein_coding | 0.215235655 | 0.081973001 | No | IFN_related_automatic |
| Green | FBXO6 | ENSG00000116663 | protein_coding | 0.226724461 | 0.083261028 | No | IFN_related_manual |
| Green | GBP3 | ENSG00000117226 | protein_coding | 0.245690883 | 0.098774619 | No | IFN_related_automatic |
| Green | GBP1 | ENSG00000117228 | protein_coding | 0.23598711 | 0.086995773 | No | IFN_related_automatic |

| | | | | | | | |
|---|---|---|---|---|---|---|---|
| Green | CDK18 | ENSG00000117266 | protein_coding | 0.093409407 | 0.001444924 | Yes | discarted_as_peripheral |
| Green | BLZF1 | ENSG00000117475 | protein_coding | 0.205659671 | 0.059662931 | No | novel |
| Green | AKAP7 | ENSG00000118507 | protein_coding | 0.124289258 | 0.004709728 | Yes | discarted_as_peripheral |
| Green | IFIT3 | ENSG00000119917 | protein_coding | 0.232424565 | 0.098354898 | No | IFN_related_automatic |
| Green | IFIT2 | ENSG00000119922 | protein_coding | 0.248488189 | 0.105913945 | No | IFN_related_automatic |
| Green | CD274 | ENSG00000120217 | protein_coding | 0.1959364 | 0.051968992 | No | IFN_related_manual |
| Green | MASTL | ENSG00000120539 | protein_coding | 0.23766853 | 0.095626906 | No | IFN_related_manual |
| Green | KIAA1217 | ENSG00000120549 | protein_coding | 0.160802014 | 0.009812185 | Yes | discarted_as_peripheral |
| Green | TRIM25 | ENSG00000121060 | protein_coding | 0.222545825 | 0.083830021 | No | IFN_related_automatic |
| Green | BCL2L14 | ENSG00000121380 | protein_coding | 0.129151387 | 0.005297245 | Yes | discarted_as_peripheral |
| Green | TNFSF10 | ENSG00000121858 | protein_coding | 0.218679764 | 0.065768309 | No | IFN_related_automatic |
| Green | FMOD | ENSG00000122176 | protein_coding | 0.095230752 | 1.24E-04 | Yes | discarted_as_peripheral |
| Green | NT5C3A | ENSG00000122643 | protein_coding | 0.173240642 | 0.028157664 | No | IFN_related_manual |
| Green | OPTN | ENSG00000123240 | protein_coding | 0.201285028 | 0.042496212 | No | IFN_related_manual |
| Green | NMI | ENSG00000123609 | protein_coding | 0.223154401 | 0.065858915 | No | IFN_related_manual |
| Green | DNPEP | ENSG00000123992 | protein_coding | 0.20821589 | 0.048152775 | No | novel |
| Green | ZNFX1 | ENSG00000124201 | protein_coding | 0.229394347 | 0.094736233 | No | IFN_related_manual |
| Green | MT2A | ENSG00000125148 | protein_coding | 0.112328862 | 0.001038033 | Yes | discarted_as_peripheral |
| Green | IRF1 | ENSG00000125347 | protein_coding | 0.156300553 | 0.012191127 | Yes | discarted_as_peripheral |
| Green | RBCK1 | ENSG00000125826 | protein_coding | 0.16648291 | 0.027688161 | No | IFN_related_automatic |
| Green | IFI6 | ENSG00000126709 | protein_coding | 0.190937499 | 0.0524683 | No | IFN_related_automatic |
| Green | TICAM1 | ENSG00000127666 | protein_coding | 0.149469127 | 0.019352656 | Yes | discarted_as_peripheral |
| Green | APOL2 | ENSG00000128335 | protein_coding | 0.250094727 | 0.099831067 | No | IFN_related_manual |
| Green | APOBEC3F | ENSG00000128394 | protein_coding | 0.162634905 | 0.025236694 | No | IFN_related_automatic |
| Green | PODXL | ENSG00000128567 | protein_coding | 0.090264677 | 2.09E-04 | Yes | discarted_as_peripheral |
| Green | BST2 | ENSG00000130303 | protein_coding | 0.248963862 | 0.108712189 | No | IFN_related_automatic |
| Green | KLHDC7B | ENSG00000130487 | protein_coding | 0.159645314 | 0.02773493 | No | IFN_related_manual |
| Green | HELZ2 | ENSG00000130589 | protein_coding | 0.20307455 | 0.064766467 | No | IFN_related_manual |
| Green | C19orf66 | ENSG00000130813 | protein_coding | 0.223802328 | 0.074698821 | No | IFN_related_manual |
| Green | CLUHP3 | ENSG00000131797 | transcribed_unproce | 0.119010257 | 0.001850162 | Yes | discarted_as_peripheral |
| Green | TRIM21 | ENSG00000132109 | protein_coding | 0.236834706 | 0.094953679 | No | IFN_related_automatic |
| Green | TRIM5 | ENSG00000132256 | protein_coding | 0.227537738 | 0.070795171 | No | IFN_related_automatic |
| Green | TRIM22 | ENSG00000132274 | protein_coding | 0.239686584 | 0.086999625 | No | IFN_related_automatic |
| Green | XAF1 | ENSG00000132530 | protein_coding | 0.253950204 | 0.110976458 | No | IFN_related_automatic |
| Green | ACY3 | ENSG00000132744 | protein_coding | 0.126464615 | 0.004110434 | Yes | discarted_as_peripheral |
| Green | EPSTI1 | ENSG00000133106 | protein_coding | 0.210309988 | 0.070945188 | No | IFN_related_manual |
| Green | RARRES3 | ENSG00000133321 | protein_coding | 0.209019621 | 0.058065783 | No | novel |
| Green | C14orf159 | ENSG00000133943 | protein_coding | 0.112706064 | 0.003425728 | Yes | discarted_as_peripheral |
| Green | RSAD2 | ENSG00000134321 | protein_coding | 0.25544813 | 0.111037053 | No | IFN_related_automatic |
| Green | CMPK2 | ENSG00000134326 | protein_coding | 0.231342356 | 0.096927032 | No | IFN_related_manual |
| Green | IL15RA | ENSG00000134470 | protein_coding | 0.147742555 | 0.016226763 | Yes | discarted_as_peripheral |
| Green | CYP2J2 | ENSG00000134716 | protein_coding | 0.17015823 | 0.016406249 | Yes | discarted_as_peripheral |
| Green | OASL | ENSG00000135114 | protein_coding | 0.255580425 | 0.111582465 | No | IFN_related_automatic |
| Green | TRAFD1 | ENSG00000135148 | protein_coding | 0.176560068 | 0.029981403 | No | IFN_related_automatic |
| Green | SP110 | ENSG00000135899 | protein_coding | 0.226052184 | 0.093254648 | No | IFN_related_automatic |
| Green | PHF11 | ENSG00000136147 | protein_coding | 0.18374501 | 0.036137883 | No | IFN_related_manual |
| Green | RTP4 | ENSG00000136514 | protein_coding | 0.214314177 | 0.067679003 | No | IFN_related_manual |
| Green | STX17 | ENSG00000136874 | protein_coding | 0.179198384 | 0.03449256 | No | IFN_related_manual |
| Green | CMTR1 | ENSG00000137200 | protein_coding | 0.219309191 | 0.072997808 | No | IFN_related_manual |
| Green | CPEB2 | ENSG00000137449 | protein_coding | 0.061330776 | 3.88E-07 | Yes | discarted_as_peripheral |
| Green | DDX60 | ENSG00000137628 | protein_coding | 0.220528231 | 0.084146418 | No | IFN_related_automatic |
| Green | CASP1 | ENSG00000137752 | protein_coding | 0.165824912 | 0.023463817 | No | IFN_related_automatic |
| Green | SQRDL | ENSG00000137767 | protein_coding | 0.134656915 | 0.010070354 | Yes | discarted_as_peripheral |
| Green | TMEM62 | ENSG00000137842 | protein_coding | 0.143139233 | 0.008596035 | Yes | discarted_as_peripheral |
| Green | IFI44L | ENSG00000137959 | protein_coding | 0.238753243 | 0.099521682 | No | IFN_related_automatic |
| Green | IFI44 | ENSG00000137965 | protein_coding | 0.229250417 | 0.078821331 | No | IFN_related_manual |
| Green | PNPT1 | ENSG00000138035 | protein_coding | 0.233480797 | 0.095808841 | No | IFN_related_manual |
| Green | PARP9 | ENSG00000138496 | protein_coding | 0.202535318 | 0.066845007 | No | IFN_related_automatic |
| Green | HERC6 | ENSG00000138642 | protein_coding | 0.219958229 | 0.086298741 | No | IFN_related_manual |
| Green | HERC5 | ENSG00000138646 | protein_coding | 0.240158584 | 0.097583786 | No | IFN_related_automatic |
| Green | ETV6 | ENSG00000139083 | protein_coding | 0.129016916 | 0.003777507 | Yes | discarted_as_peripheral |
| Green | NDUFA9 | ENSG00000139180 | protein_coding | 0.077330233 | 4.39E-06 | Yes | discarted_as_peripheral |
| Green | TAPBPL | ENSG00000139192 | protein_coding | 0.149584666 | 0.013778913 | Yes | discarted_as_peripheral |

| | | | | | | | |
|---|---|---|---|---|---|---|---|
| Green | PML | ENSG00000140464 | protein_coding | 0.217696025 | 0.071758598 | No | IFN_related_automatic |
| Green | HAPLN3 | ENSG00000140511 | protein_coding | 0.16174765 | 0.024511663 | No | IFN_related_manual |
| Green | MARVELD3 | ENSG00000140832 | protein_coding | 0.08927658 | 4.57E-05 | Yes | discarted_as_peripheral |
| Green | NLRC5 | ENSG00000140853 | protein_coding | 0.242869859 | 0.09512601 | No | IFN_related_automatic |
| Green | ZMYND15 | ENSG00000141497 | protein_coding | 0.058744574 | 6.18E-06 | Yes | discarted_as_peripheral |
| Green | ZCCHC2 | ENSG00000141664 | protein_coding | 0.128065703 | 0.001794408 | Yes | discarted_as_peripheral |
| Green | IFITM3 | ENSG00000142089 | protein_coding | 0.224102878 | 0.090613499 | No | IFN_related_automatic |
| Green | IL22RA1 | ENSG00000142677 | protein_coding | 0.215754859 | 0.065197036 | No | IFN_related_manual |
| Green | MOB3C | ENSG00000142961 | protein_coding | 0.209831279 | 0.066204831 | No | novel |
| Green | NME7 | ENSG00000143156 | protein_coding | 0.171139751 | 0.043839373 | No | novel |
| Green | RFX5 | ENSG00000143390 | protein_coding | 0.13639078 | 0.004374195 | Yes | discarted_as_peripheral |
| Green | DUSP10 | ENSG00000143507 | protein_coding | 0.104859395 | 0.004640053 | Yes | discarted_as_peripheral |
| Green | GRIP2 | ENSG00000144596 | protein_coding | 0.203919138 | 0.055311032 | No | novel |
| Green | NR1I2 | ENSG00000144852 | protein_coding | 0.13176008 | 0.004396167 | Yes | discarted_as_peripheral |
| Green | OSMR | ENSG00000145623 | protein_coding | 0.213967477 | 0.069045022 | No | IFN_related_automatic |
| Green | TMEM140 | ENSG00000146859 | protein_coding | 0.162925983 | 0.022699313 | No | IFN_related_manual |
| Green | SERPING1 | ENSG00000149131 | protein_coding | 0.17716389 | 0.025919719 | No | IFN_related_automatic |
| Green | GPR158 | ENSG00000151025 | protein_coding | 0.223492207 | 0.075149968 | No | novel |
| Green | SCLT1 | ENSG00000151466 | protein_coding | 0.12464683 | 0.003000153 | Yes | discarted_as_peripheral |
| Green | C4orf33 | ENSG00000151470 | protein_coding | 0.180819972 | 0.041506186 | No | IFN_related_manual |
| Green | RASGRP3 | ENSG00000152689 | protein_coding | 0.235491974 | 0.078874722 | No | IFN_related_manual |
| Green | IFIT5 | ENSG00000152778 | protein_coding | 0.212267406 | 0.077993117 | No | IFN_related_automatic |
| Green | TTC39B | ENSG00000155158 | protein_coding | 0.074434108 | 2.60E-04 | Yes | discarted_as_peripheral |
| Green | SLC25A28 | ENSG00000155287 | protein_coding | 0.199966428 | 0.054659013 | No | IFN_related_manual |
| Green | MOV10 | ENSG00000155363 | protein_coding | 0.217660837 | 0.062492717 | No | IFN_related_automatic |
| Green | PIK3AP1 | ENSG00000155629 | protein_coding | 0.226800742 | 0.071874857 | No | IFN_related_automatic |
| Green | WHAMM | ENSG00000156232 | protein_coding | 0.11216525 | 5.87E-04 | Yes | discarted_as_peripheral |
| Green | FAM122C | ENSG00000156500 | protein_coding | 0.118009965 | 0.005551021 | Yes | discarted_as_peripheral |
| Green | UBE2L6 | ENSG00000156587 | protein_coding | 0.245421679 | 0.102512263 | No | IFN_related_automatic |
| Green | B3GNT7 | ENSG00000156966 | protein_coding | 0.160941691 | 0.018883139 | Yes | discarted_as_peripheral |
| Green | SUSD3 | ENSG00000157303 | protein_coding | 0.106188771 | 0.001512491 | Yes | discarted_as_peripheral |
| Green | MX1 | ENSG00000157601 | protein_coding | 0.191612421 | 0.054332305 | No | IFN_related_automatic |
| Green | TNFRSF14 | ENSG00000157873 | protein_coding | 0.107436865 | 0.008314769 | Yes | discarted_as_peripheral |
| Green | C1R | ENSG00000159403 | protein_coding | 0.182107202 | 0.034685642 | No | IFN_related_automatic |
| Green | ADAR | ENSG00000160710 | protein_coding | 0.230742916 | 0.094332993 | No | IFN_related_automatic |
| Green | MFSD12 | ENSG00000161091 | protein_coding | 0.183547481 | 0.037931588 | No | novel |
| Green | GBP4 | ENSG00000162654 | protein_coding | 0.150053443 | 0.012084599 | Yes | discarted_as_peripheral |
| Green | KCNT2 | ENSG00000162687 | protein_coding | 0.17511694 | 0.026298627 | No | novel |
| Green | ATF3 | ENSG00000162772 | protein_coding | 0.140866482 | 0.011199238 | Yes | discarted_as_peripheral |
| Green | NEURL3 | ENSG00000163121 | protein_coding | 0.075068719 | 8.94E-05 | Yes | discarted_as_peripheral |
| Green | CTSS | ENSG00000163131 | protein_coding | 0.146658687 | 0.023192428 | No | IFN_related_automatic |
| Green | PPM1K | ENSG00000163644 | protein_coding | 0.211028469 | 0.069798402 | No | IFN_related_manual |
| Green | IL17RC | ENSG00000163702 | protein_coding | 0.063512755 | 2.49E-07 | Yes | discarted_as_peripheral |
| Green | CCR1 | ENSG00000163823 | protein_coding | 0.132307474 | 0.004064014 | Yes | discarted_as_peripheral |
| Green | DTX3L | ENSG00000163840 | protein_coding | 0.211369455 | 0.076331049 | No | IFN_related_manual |
| Green | IL15 | ENSG00000164136 | protein_coding | 0.215992743 | 0.071347535 | No | IFN_related_automatic |
| Green | ELOVL7 | ENSG00000164181 | protein_coding | 0.128598514 | 0.003724444 | Yes | discarted_as_peripheral |
| Green | STARD4 | ENSG00000164211 | protein_coding | 0.129389788 | 0.004657988 | Yes | discarted_as_peripheral |
| Green | ERAP1 | ENSG00000164307 | protein_coding | 0.240070887 | 0.094951354 | No | IFN_related_automatic |
| Green | ERAP2 | ENSG00000164308 | protein_coding | 0.235027716 | 0.08737836 | No | IFN_related_automatic |
| Green | TLR3 | ENSG00000164342 | protein_coding | 0.237475884 | 0.085464716 | No | IFN_related_automatic |
| Green | CASP7 | ENSG00000165806 | protein_coding | 0.252034842 | 0.099869221 | No | IFN_related_manual |
| Green | IFI27 | ENSG00000165949 | protein_coding | 0.228435976 | 0.075843796 | No | IFN_related_automatic |
| Green | C2 | ENSG00000166278 | protein_coding | 0.115101928 | 0.001324776 | Yes | discarted_as_peripheral |
| Green | PLEKHA7 | ENSG00000166689 | protein_coding | 0.181994942 | 0.028800027 | No | novel |
| Green | B2M | ENSG00000166710 | protein_coding | 0.254246346 | 0.104347132 | No | IFN_related_automatic |
| Green | RILP | ENSG00000167705 | protein_coding | 0.077613107 | 3.17E-05 | Yes | discarted_as_peripheral |
| Green | TRANK1 | ENSG00000168016 | protein_coding | 0.206920207 | 0.061357666 | No | IFN_related_manual |
| Green | BATF2 | ENSG00000168062 | protein_coding | 0.230154423 | 0.089705303 | No | IFN_related_manual |
| Green | PXK | ENSG00000168297 | protein_coding | 0.187607816 | 0.039327201 | No | IFN_related_manual |
| Green | IRF2 | ENSG00000168310 | protein_coding | 0.163168308 | 0.033039335 | No | IFN_related_automatic |
| Green | TAP1 | ENSG00000168394 | protein_coding | 0.244164696 | 0.104958609 | No | IFN_related_automatic |
| Green | MLKL | ENSG00000168404 | protein_coding | 0.190639503 | 0.041570615 | No | IFN_related_manual |

| | | | | | | | |
|---|---|---|---|---|---|---|---|
| Green | VAMP5 | ENSG00000168899 | protein_coding | 0.126530198 | 0.001755142 | Yes | discarted_as_peripheral |
| Green | CXCL10 | ENSG00000169245 | protein_coding | 0.225980679 | 0.066115357 | No | IFN_related_automatic |
| Green | CXCL11 | ENSG00000169248 | protein_coding | 0.224847746 | 0.068350129 | No | IFN_related_manual |
| Green | CXCL8 | ENSG00000169429 | protein_coding | 0.180503337 | 0.038263079 | No | IFN_related_automatic |
| Green | TRIM56 | ENSG00000169871 | protein_coding | 0.158939171 | 0.013484839 | Yes | discarted_as_peripheral |
| Green | MUC3A | ENSG00000169894 | protein_coding | 0.141970213 | 0.018019892 | Yes | discarted_as_peripheral |
| Green | STAT2 | ENSG00000170581 | protein_coding | 0.254507075 | 0.111166485 | No | IFN_related_automatic |
| Green | PRKCE | ENSG00000171132 | protein_coding | 0.119513241 | 0.003217673 | Yes | discarted_as_peripheral |
| Green | ISG20 | ENSG00000172183 | protein_coding | 0.230192959 | 0.07103809 | No | IFN_related_automatic |
| Green | MYD88 | ENSG00000172936 | protein_coding | 0.235515184 | 0.079775892 | No | IFN_related_automatic |
| Green | PARP14 | ENSG00000173193 | protein_coding | 0.231351844 | 0.098663017 | No | IFN_related_manual |
| Green | GLRX | ENSG00000173221 | protein_coding | 0.073629908 | 5.54E-05 | Yes | discarted_as_peripheral |
| Green | MUC13 | ENSG00000173702 | protein_coding | 0.14855824 | 0.005796627 | Yes | discarted_as_peripheral |
| Green | CNP | ENSG00000173786 | protein_coding | 0.243430043 | 0.090107851 | No | IFN_related_manual |
| Green | RNF213 | ENSG00000173821 | protein_coding | 0.245763028 | 0.094950333 | No | IFN_related_manual |
| Green | BTC | ENSG00000174808 | protein_coding | 0.124246639 | 0.020059366 | No | IFN_related_automatic |
| Green | SAMD9L | ENSG00000177409 | protein_coding | 0.247699484 | 0.102993734 | No | IFN_related_manual |
| Green | ANXA2R | ENSG00000177721 | protein_coding | 0.110534574 | 0.006735148 | Yes | discarted_as_peripheral |
| Green | ODF3B | ENSG00000177989 | protein_coding | 0.15850575 | 0.012157131 | Yes | discarted_as_peripheral |
| Green | PARP10 | ENSG00000178685 | protein_coding | 0.243570578 | 0.104663603 | No | IFN_related_automatic |
| Green | GRINA | ENSG00000178719 | protein_coding | 0.17061791 | 0.025306124 | No | novel |
| Green | ZBTB42 | ENSG00000179627 | protein_coding | 0.11904427 | 0.003459392 | Yes | discarted_as_peripheral |
| Green | PCGF5 | ENSG00000180628 | protein_coding | 0.146280175 | 0.013818626 | Yes | discarted_as_peripheral |
| Green | DDX60L | ENSG00000181381 | protein_coding | 0.229576082 | 0.089804649 | No | IFN_related_manual |
| Green | UBA7 | ENSG00000182179 | protein_coding | 0.247492498 | 0.093210287 | No | IFN_related_automatic |
| Green | C1S | ENSG00000182326 | protein_coding | 0.229905341 | 0.077775717 | No | IFN_related_automatic |
| Green | CSF1 | ENSG00000184371 | protein_coding | 0.127961231 | 0.003788824 | Yes | discarted_as_peripheral |
| Green | RBM43 | ENSG00000184898 | protein_coding | 0.207805949 | 0.060915463 | No | IFN_related_manual |
| Green | USP18 | ENSG00000184979 | protein_coding | 0.231144753 | 0.095446236 | No | IFN_related_automatic |
| Green | IFITM2 | ENSG00000185201 | protein_coding | 0.120251538 | 0.005220023 | Yes | discarted_as_peripheral |
| Green | SP140L | ENSG00000185404 | protein_coding | 0.244760085 | 0.089951858 | No | IFN_related_manual |
| Green | IFIT1 | ENSG00000185745 | protein_coding | 0.203589234 | 0.068312302 | No | IFN_related_automatic |
| Green | TRIM69 | ENSG00000185880 | protein_coding | 0.206630868 | 0.065944552 | No | IFN_related_manual |
| Green | IFITM1 | ENSG00000185885 | protein_coding | 0.224227802 | 0.077266037 | No | IFN_related_automatic |
| Green | BTN3A2 | ENSG00000186470 | protein_coding | 0.227060006 | 0.081003323 | No | IFN_related_manual |
| Green | ISG15 | ENSG00000187608 | protein_coding | 0.214715573 | 0.080647588 | No | IFN_related_automatic |
| Green | HES4 | ENSG00000188290 | protein_coding | 0.104429732 | 1.86E-04 | Yes | discarted_as_peripheral |
| Green | PLSCR1 | ENSG00000188313 | protein_coding | 0.221182654 | 0.088024355 | No | IFN_related_automatic |
| Green | TDRD7 | ENSG00000196116 | protein_coding | 0.223540063 | 0.075630113 | No | IFN_related_manual |
| Green | ZNF107 | ENSG00000196247 | protein_coding | 0.163147779 | 0.025969006 | No | novel |
| Green | CASP4 | ENSG00000196954 | protein_coding | 0.211185931 | 0.055913627 | No | IFN_related_automatic |
| Green | SERTAD1 | ENSG00000197019 | protein_coding | 0.073742454 | 1.01E-04 | Yes | discarted_as_peripheral |
| Green | ACSL5 | ENSG00000197142 | protein_coding | 0.084314162 | 7.94E-05 | Yes | discarted_as_peripheral |
| Green | C5orf56 | ENSG00000197536 | protein_coding | 0.18852751 | 0.045280699 | No | IFN_related_manual |
| Green | TGM2 | ENSG00000198959 | protein_coding | 0.196414396 | 0.049686186 | No | IFN_related_automatic |
| Green | PSMB8 | ENSG00000204264 | protein_coding | 0.22843768 | 0.080624864 | No | IFN_related_automatic |
| Green | TAP2 | ENSG00000204267 | protein_coding | 0.238860835 | 0.094357088 | No | IFN_related_automatic |
| Green | CARD16 | ENSG00000204397 | protein_coding | 0.140689683 | 0.009411601 | Yes | discarted_as_peripheral |
| Green | HLA-C | ENSG00000204525 | protein_coding | 0.200895397 | 0.051779605 | No | IFN_related_automatic |
| Green | HLA-E | ENSG00000204592 | protein_coding | 0.224749519 | 0.073800802 | No | IFN_related_automatic |
| Green | PSMB10 | ENSG00000205220 | protein_coding | 0.167105463 | 0.022798637 | No | IFN_related_automatic |
| Green | SAMD9 | ENSG00000205413 | protein_coding | 0.23935225 | 0.10259074 | No | IFN_related_manual |
| Green | HLA-H | ENSG00000206341 | unprocessed_pseudo | 0.177160522 | 0.03389293 | No | IFN_related_manual |
| Green | HLA-A | ENSG00000206503 | protein_coding | 0.133822812 | 0.004203309 | Yes | discarted_as_peripheral |
| Green | LAP3P2 | ENSG00000213500 | processed_pseudoge | 0.180753505 | 0.053195067 | No | novel |
| Green | IRF9 | ENSG00000213928 | protein_coding | 0.173969903 | 0.038680401 | No | IFN_related_automatic |
| Green | GANC | ENSG00000214013 | protein_coding | 0.070252959 | 6.57E-05 | Yes | discarted_as_peripheral |
| Green | C17orf67 | ENSG00000214226 | protein_coding | 0.119187477 | 0.002344829 | Yes | discarted_as_peripheral |
| Green | SMTNL1 | ENSG00000214872 | protein_coding | 0.191905032 | 0.053213816 | No | novel |
| Green | AC025171.1 | ENSG00000215068 | antisense | 0.068534027 | 8.07E-05 | Yes | discarted_as_peripheral |
| Green | APOL6 | ENSG00000221963 | protein_coding | 0.246034189 | 0.105711676 | No | IFN_related_manual |
| Green | AC009948.5 | ENSG00000223960 | antisense | 0.189141107 | 0.056868635 | No | novel |
| Green | PSME2P2 | ENSG00000225131 | processed_pseudoge | 0.199006257 | 0.052494377 | No | novel |

| | | | | | | | |
|---|---|---|---|---|---|---|---|
| Green | NRSN2-AS1 | ENSG00000225377 | antisense | 0.085426025 | 5.34E-04 | Yes | discarted_as_peripheral |
| Green | GBP1P1 | ENSG00000225492 | transcribed_unproce: | 0.123289726 | 0.007141426 | Yes | discarted_as_peripheral |
| Green | RP11-288L9.4 | ENSG00000225886 | antisense | 0.187706232 | 0.050173867 | No | IFN_related_manual |
| Green | AC009950.2 | ENSG00000225963 | antisense | 0.199429251 | 0.065825321 | No | novel |
| Green | AP001610.5 | ENSG00000228318 | antisense | 0.204791714 | 0.070108816 | No | novel |
| Green | PNPT1P1 | ENSG00000229241 | processed_pseudoge | 0.190170213 | 0.058537211 | No | novel |
| Green | PATL2 | ENSG00000229474 | protein_coding | 0.174053225 | 0.04639047 | No | novel |
| Green | NAMPTL | ENSG00000229644 | processed_pseudoge | 0.193361923 | 0.052664574 | No | novel |
| Green | RP11-274B21.4 | ENSG00000230715 | processed_pseudoge | 0.055135844 | 4.63E-06 | Yes | discarted_as_peripheral |
| Green | HLA-K | ENSG00000230795 | unprocessed_pseudo | 0.163235097 | 0.024861276 | No | novel |
| Green | TAPBP | ENSG00000231925 | protein_coding | 0.228586686 | 0.081228063 | No | IFN_related_automatic |
| Green | RP5-1011O1.2 | ENSG00000232498 | antisense | 0.080383875 | 1.99E-04 | Yes | discarted_as_peripheral |
| Green | TRIM26 | ENSG00000234127 | protein_coding | 0.207711222 | 0.053328767 | No | IFN_related_automatic |
| Green | HLA-B | ENSG00000234745 | protein_coding | 0.229662869 | 0.080813542 | No | IFN_related_automatic |
| Green | RP11-274E7.2 | ENSG00000238000 | processed_pseudoge | 0.211860234 | 0.061567478 | No | novel |
| Green | PSMB9 | ENSG00000240065 | protein_coding | 0.24366183 | 0.092806607 | No | IFN_related_automatic |
| Green | RP11-274B21.2 | ENSG00000243302 | processed_pseudoge | 0.057500115 | 2.60E-06 | Yes | discarted_as_peripheral |
| Green | CFB | ENSG00000243649 | protein_coding | 0.081785751 | 2.52E-04 | Yes | discarted_as_peripheral |
| Green | RN7SL834P | ENSG00000243650 | misc_RNA | 0.133006915 | 0.018869372 | Yes | discarted_as_peripheral |
| Green | SCAMP1-AS1 | ENSG00000245556 | lincRNA | 0.104106124 | 0.005351879 | Yes | discarted_as_peripheral |
| Green | AP001258.4 | ENSG00000245571 | lincRNA | 0.126054686 | 0.004397237 | Yes | discarted_as_peripheral |
| Green | RP11-468E2.4 | ENSG00000259529 | protein_coding | 0.15012136 | 0.024527613 | No | IFN_related_automatic |
| Green | RP11-973H7.1 | ENSG00000260302 | lincRNA | 0.096200381 | 0.001873162 | Yes | discarted_as_peripheral |
| Green | CTD-2047H16.4 | ENSG00000263069 | antisense | 0.091192264 | 0.002640277 | Yes | discarted_as_peripheral |
| Green | IKBKE | ENSG00000263528 | protein_coding | 0.142882951 | 0.004834677 | Yes | discarted_as_peripheral |
| Green | BAHCC1 | ENSG00000266074 | protein_coding | 0.13060528 | 0.001079202 | Yes | discarted_as_peripheral |
| Green | CTD-2521M24.9 | ENSG00000269640 | lincRNA | 0.240616473 | 0.092916161 | No | IFN_related_manual |
| Green | BNIP3P11 | ENSG00000271550 | processed_pseudoge | 0.087365356 | 3.39E-04 | Yes | discarted_as_peripheral |
| Green | RP11-54O7.17 | ENSG00000272512 | lincRNA | 0.147850379 | 0.028382566 | No | novel |
| Green | DCP1A | ENSG00000272886 | protein_coding | 0.136034284 | 0.006187527 | Yes | discarted_as_peripheral |
| Green | AC008079.10 | ENSG00000280007 | antisense | 0.069585032 | 9.70E-05 | Yes | discarted_as_peripheral |
| Tan | CFH | ENSG00000000971 | protein_coding | 0.208253108 | 0.288528055 | No | IFN_related_automatic |
| Tan | BIRC3 | ENSG00000023445 | protein_coding | 0.152159792 | 0.138114542 | No | IFN_related_automatic |
| Tan | STOML1 | ENSG00000067221 | protein_coding | 0.130676031 | 0.09908808 | No | novel |
| Tan | CXCL2 | ENSG00000081041 | protein_coding | 0.133377738 | 0.067603692 | No | IFN_related_automatic |
| Tan | VNN3 | ENSG00000093134 | protein_coding | 0.15462185 | 0.159602374 | No | IFN_related_manual |
| Tan | NANS | ENSG00000095380 | protein_coding | 0.132035409 | 0.086724579 | No | novel |
| Tan | PCK2 | ENSG00000100889 | protein_coding | 0.141503654 | 0.109922789 | No | IFN_related_manual |
| Tan | IL7 | ENSG00000104432 | protein_coding | 0.163914175 | 0.176232323 | No | IFN_related_manual |
| Tan | LIPA | ENSG00000107798 | protein_coding | 0.203175597 | 0.280185908 | No | IFN_related_automatic |
| Tan | SOD2 | ENSG00000112096 | protein_coding | 0.198212525 | 0.25683105 | No | IFN_related_manual |
| Tan | CCL20 | ENSG00000115009 | protein_coding | 0.187745604 | 0.22917369 | No | IFN_related_automatic |
| Tan | GCA | ENSG00000115271 | protein_coding | 0.158766491 | 0.176857919 | No | IFN_related_manual |
| Tan | ATP10B | ENSG00000118322 | protein_coding | 0.1262987 | 0.093177997 | No | novel |
| Tan | TNFAIP3 | ENSG00000118503 | protein_coding | 0.166023064 | 0.154663488 | No | IFN_related_automatic |
| Tan | TRIM6 | ENSG00000121236 | protein_coding | 0.184096148 | 0.235374123 | No | IFN_related_manual |
| Tan | INHBA | ENSG00000122641 | protein_coding | 0.102087152 | 0.050586876 | No | IFN_related_manual |
| Tan | ATP8A1 | ENSG00000124406 | protein_coding | 0.135206262 | 0.097911425 | No | IFN_related_manual |
| Tan | PRRG4 | ENSG00000135378 | protein_coding | 0.122164945 | 0.067685507 | No | novel |
| Tan | NIPSNAP3A | ENSG00000136783 | protein_coding | 0.169441056 | 0.190861166 | No | novel |
| Tan | SPTBN5 | ENSG00000137877 | protein_coding | 0.15483781 | 0.155907214 | No | IFN_related_manual |
| Tan | TNFRSF11A | ENSG00000141655 | protein_coding | 0.151460045 | 0.127883583 | No | IFN_related_automatic |
| Tan | PMAIP1 | ENSG00000141682 | protein_coding | 0.203018025 | 0.270569861 | No | IFN_related_manual |
| Tan | ATHL1 | ENSG00000142102 | protein_coding | 0.150161292 | 0.133258983 | No | novel |
| Tan | NFKBIZ | ENSG00000144802 | protein_coding | 0.172202013 | 0.15745303 | No | IFN_related_automatic |
| Tan | GPR128 | ENSG00000144820 | protein_coding | 0.115496227 | 0.0728563 | No | novel |
| Tan | TMCO4 | ENSG00000162542 | protein_coding | 0.125966053 | 0.082562499 | No | novel |
| Tan | GBP2 | ENSG00000162645 | protein_coding | 0.166696601 | 0.175455019 | No | IFN_related_manual |
| Tan | CXCL1 | ENSG00000163739 | protein_coding | 0.140776671 | 0.085937761 | No | IFN_related_automatic |
| Tan | AGBL2 | ENSG00000165923 | protein_coding | 0.104189999 | 0.065924302 | No | novel |
| Tan | ABTB2 | ENSG00000166016 | protein_coding | 0.130669226 | 0.073767707 | No | IFN_related_manual |
| Tan | FAM111A | ENSG00000166801 | protein_coding | 0.203205851 | 0.281017154 | No | IFN_related_manual |
| Tan | SDPR | ENSG00000168497 | protein_coding | 0.153764232 | 0.152868837 | No | IFN_related_manual |

| | | | | | | | |
|---|---|---|---|---|---|---|---|
| Tan | SLFN12 | ENSG00000172123 | protein_coding | 0.141521843 | 0.122315012 | No | IFN_related_manual |
| Tan | SLCO4C1 | ENSG00000173930 | protein_coding | 0.120313216 | 0.062255494 | No | IFN_related_manual |
| Tan | TNFAIP2 | ENSG00000185215 | protein_coding | 0.151119575 | 0.133611843 | No | IFN_related_manual |
| Tan | PLEKHN1 | ENSG00000187583 | protein_coding | 0.092999834 | 0.046961054 | No | novel |
| Tan | DPYD | ENSG00000188641 | protein_coding | 0.153642166 | 0.1299005 | No | IFN_related_manual |
| Tan | TLR7 | ENSG00000196664 | protein_coding | 0.132739569 | 0.100283638 | No | IFN_related_automatic |
| Tan | CD47 | ENSG00000196776 | protein_coding | 0.154378804 | 0.135411993 | No | IFN_related_automatic |
| Tan | AC009299.3 | ENSG00000227403 | lincRNA | 0.109916834 | 0.059511922 | No | novel |
| Tan | RNF144A-AS1 | ENSG00000228203 | processed_transcript | 0.111994798 | 0.072721019 | No | novel |
| Tan | RP11-395B7.7 | ENSG00000260336 | sense_overlapping | 0.14801089 | 0.140429541 | No | novel |
| Tan | CD74 | ENSG00000019582 | protein_coding | 0.071491216 | 0.005177548 | Yes | discarted_as_peripheral |
| Tan | IL2RG | ENSG00000147168 | protein_coding | 0.091325963 | 0.018062755 | Yes | discarted_as_peripheral |
| Tan | TNFRSF11B | ENSG00000164761 | protein_coding | 0.108318728 | 0.020469727 | No | IFN_related_manual |
| Tan | CYP2B6 | ENSG00000197408 | protein_coding | 0.099967504 | 0.021095663 | No | IFN_related_manual |
| Tan | CADPS2 | ENSG00000081803 | protein_coding | 0.122082847 | 0.021566868 | No | novel |
| Tan | CXCL3 | ENSG00000163734 | protein_coding | 0.112225 | 0.036597222 | No | IFN_related_automatic |
| Tan | TMEM51 | ENSG00000171729 | protein_coding | 0.127903008 | 0.043761492 | No | IFN_related_manual |
| Tan | MAFF | ENSG00000185022 | protein_coding | 0.098585769 | 0.012566932 | Yes | discarted_as_peripheral |
| Tan | CTSO | ENSG00000256043 | protein_coding | 0.100041319 | 0.028340813 | No | IFN_related_manual |
| Tan | KLF4 | ENSG00000136826 | protein_coding | 0.079025606 | 0.003240423 | Yes | discarted_as_peripheral |
| Tan | RP11-492E3.2 | ENSG00000227619 | antisense | 0.076053571 | 0.015939755 | Yes | discarted_as_peripheral |
| Tan | UBD | ENSG00000213886 | protein_coding | 0.072127145 | 0.00368002 | Yes | discarted_as_peripheral |
