## Supplemental Figures and Tables for "Novel antiviral interferon sensitive genes unveiled by correlation-driven gene selection and integrated systems biology approaches"

Cheroni *et al.*

**Supplementary Table 1.** Published datasets of human samples treated with IFN or HCV infected, used to validate the ISG modules signatures. Dataset title and GEO/NCBI accession number of cognate publication are reported.

| **Reference** | **Title** | **GEO ID** |
|---|---|---|
| Bolen et al. 2014 | Dynamic expression profiling of type I and type III interferon-stimulated hepatocytes. | GSE48400 |
| Carnero et al. 2014 | Type I interferon regulates the expression of long non-coding RNAs | GSE64796 |
| Blackham et al. 2010 | The Effect of Hepatitis C Virus Infection on Host Gene Expression | GSE20948 |
| Grünvogel et al. 2015 | Comparison of Huh7 cells under IFN alpha and gamma treatment | GSE68927 |

### Supplementary Figures Legends

**Supplementary Figure 1. Results of differential expression analysis. a.** PCA analysis performed on the 43 samples of the dataset. Samples are clustered according to treatment and cell identity; no outliers are detected. Volcano plots represent the outcome of differential expression analysis for (**b**) Huh7 and (**c**) Huh7.5 cells treated with IFNα, IFNβ and IFNλ (from left to right panel). Results obtained by comparing IFN-treated versus mock-treated cells are reported for the pool of tested genes as –log10FDR and log2FC. Genes identified as significantly modulated (FDR < 5% and absolute log2FC > 1) are shown in red, while those respecting only the FDR threshold are depicted in grey and not significant genes (FDR > 5%) are coloured in white.

**Supplementary Figure 2. Functional Enrichment differences between cell-line specific DE genes. a.** Results of overall functional enrichment screening for ontologies (GO), pathways (KEGG, Reactome, WP), motifs (Transfac, miRtarBase), protein (Human protein Atlas, Corum) and human phenotype (HP). performed with g:Profiler on mutually exclusive gene subsets as shown in the Venn diagram for the IFN alpha perturbed cell lines. **b.** Same results for the IFN beta perturbed cell lines.

**Supplementary Figure 3. Multi-dimensional scaling and quantitative traits measurements a.** Multidimensional scaling depicting similarities across the data set in a tri-dimensional space. Each dot represents a gene and is colored according to module assignment; genes belonging to the same module are clustered together (T: treated samples; M: mock samples) **b-c.** Quantification of IFIT2 and MX1 transcripts by real-time PCR. Bar plots representing real-time PCR transcript measurement of IFIT2 (**b**) and MX1 (**c**) in the 43 samples of the dataset. For each treatment, gene expression is quantified as -ΔΔCt after normalization for the endogenous control (Gapdh) and for the mean ΔCt value in respective mock-treated cells. Values relative to each biological sample are reported and have been employed for the correlation analysis with module eigengenes in order to identify modules related to the interferon response (results presented in Figure 2b).

**Supplementary Figure 4. Modules Gene expression. a, c**. Heat maps illustrating the expression profiles of the genes belonging to the Green and Tan modules, respectively. **b, d.** Dendrograms of sample clustering according to the gene expression profiles (variance stabilizing transformed values) of the Green (**b**) or the Tan module (**d**). Each sample is identified by a name reporting the cell line, the treatment (T: treated samples; M: mock samples) and a number representative of the biological replicate.

**Supplementary Figure 5: Regulatory relationships of motif clusters. a**. Circular plot showing regulatory relationship between the Green module generated motif clusters (Green M1 and M2) and their putative target genes. 200 out of 282 genes of the green sub-network are putative targets of at least one motif family, with 163 regulated by both, as indicated by the blue arrowed line (14 only M1; 23 only M2). **b**. Same relationship plot for motif clusters Tan M1 and Tan M2. A total of 30 putative to genes were identified, with 9 genes predicted to be the target of both transcription factors, as indicated by the blue arrowed line (19 only M1, 2 only M2).

**Supplementary Figure 6. Network structure and node pruning for the Green module sub-network. a**. Sub-network generated in Cytoscape for the Green module. The network has 282 nodes, and more than 18000 edges quantifying the strengths of co-modulation of these genes. Size of the nodes represents the eigenvector value and the colour the closeness, with yellow nodes showing the highest closeness values. The metrics are quite aligned and identify a group of more relevant players and a group of more peripheral ones. **b**. With an eigenvector threshold of 0.02 most peripheral players are eliminated, and 186 genes are left. **c**. The remaining 28 genes after the subtractive gene ontology approach, among which 11 long-non coding RNAs, as putative novel players in interferon response.

**Supplementary Figure 7. Network structure and node pruning for the Tan module sub-network. a**. With an eigenvector threshold of 0.02 most peripheral nodes in the Tan module are eliminated, and 48 non-peripheral genes are shown. **b**. The remaining 14 genes after the subtractive gene ontology approach, among which 3 long-non coding RNAs, as putative novel players in interferon response.

**Supplementary Figure 8. IFN-alpha induction from GSE48400.** Data from Bolen et al 2014, analyzed by GSEA method using custom genesets with genes from the Green module, the tan module and combination of the two. The panels show the enrichment running sums for datasets of five times after induction

**Supplementary Figure 9. IFN-beta induction from GSE48400.** Data from Bolen et al 2014, analyzed by GSEA method using custom genesets with genes from the Green module, the tan module and combination of the two. The panels show the enrichment running sums for datasets of four times after induction

**Supplementary Figure 10. IFN-lambda induction from GSE48400.** Data from Bolen et al 2014, analyzed by GSEA method using custom genesets with genes from the Green module, the tan module and combination of the two. The panels show the enrichment running sums for datasets of four times after induction

**Supplementary Figure 11. IFN-alpha2 induction from GSE64796.** Data from Carnero et al 2014, analyzed by GSEA method using custom genesets with genes from the Green module, the tan module and combination of the two. Geneset from the Tan module alone did not show any enrichment

**Supplementary Figure 12. IFN-lambda1 induction from GSE20948.** Data from Blackham et al 2010, analyzed by GSEA method using custom genesets with genes from the Green module, the tan module and combination of the two. The panels show the enrichment running sums for datasets of three times after induction. Geneset from the Tan module alone did not show any enrichment

**Supplementary Figure 13. IFN-alpha induction from GSE68927.** Data from Grünvogel et al 2015, analyzed by GSEA method using custom genesets with genes from the Green module, the tan module and combination of the two. Geneset from the Tan module alone and from the combination of the two did not show any significative enrichment

### Supplementary files description

**Supplementary File 1.** Text file listing all differentially expressed genes at the set cut-off values for each cell line (Huh7 or Huh7.5) and perturbation with IFN type (alpha, beta, and lamda).

**Supplementary File 2.** Text file listing all genes (ensembl gene ids) from green (365 genes) and tan (65 genes) modules.

**Supplementary File 3.** Text file listing hub genes (subnetworks of highly connected genes) from green and tan modules, with their calculated network weights (closeness, eigenvector)

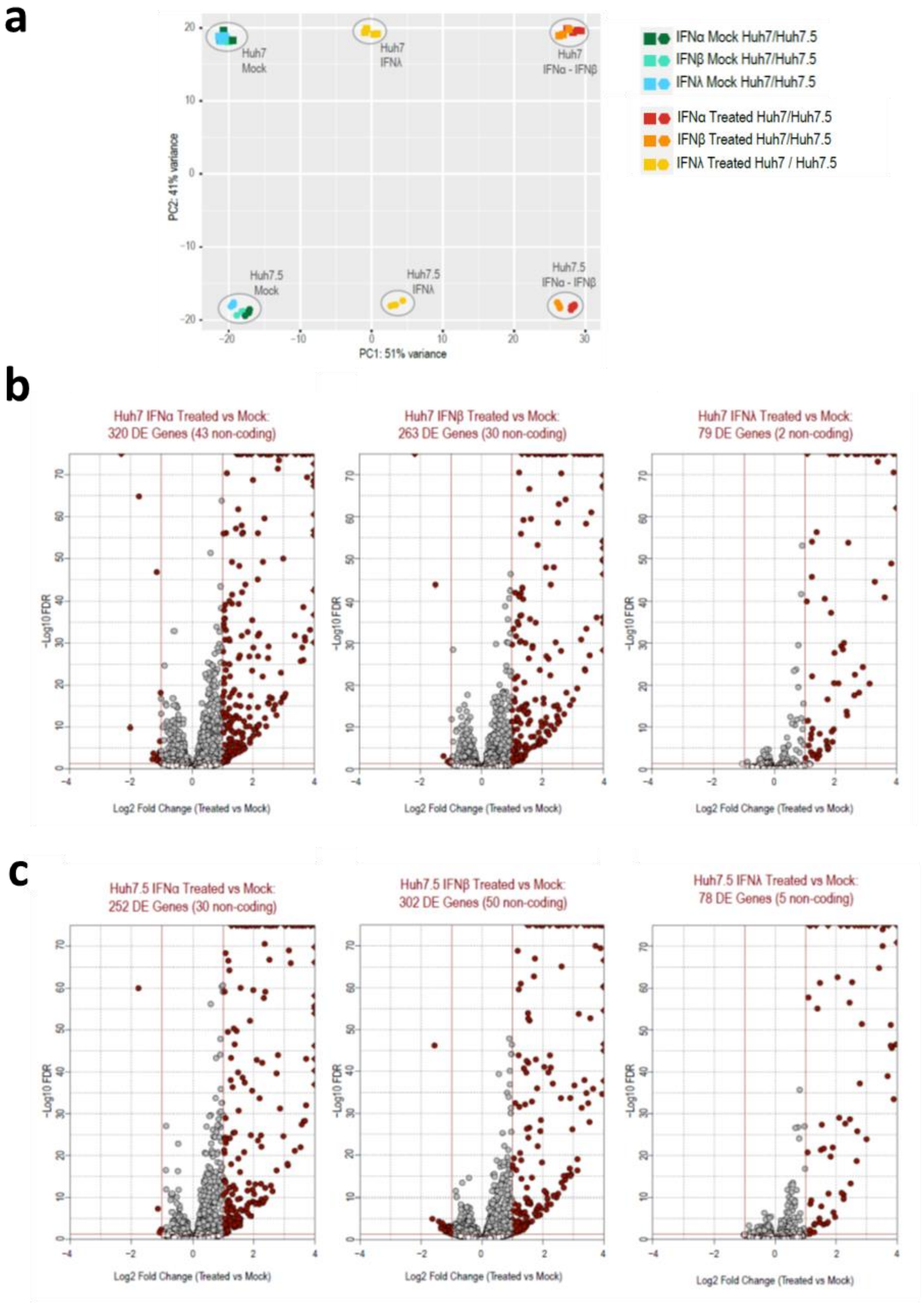


**Supplementary Figure 1**

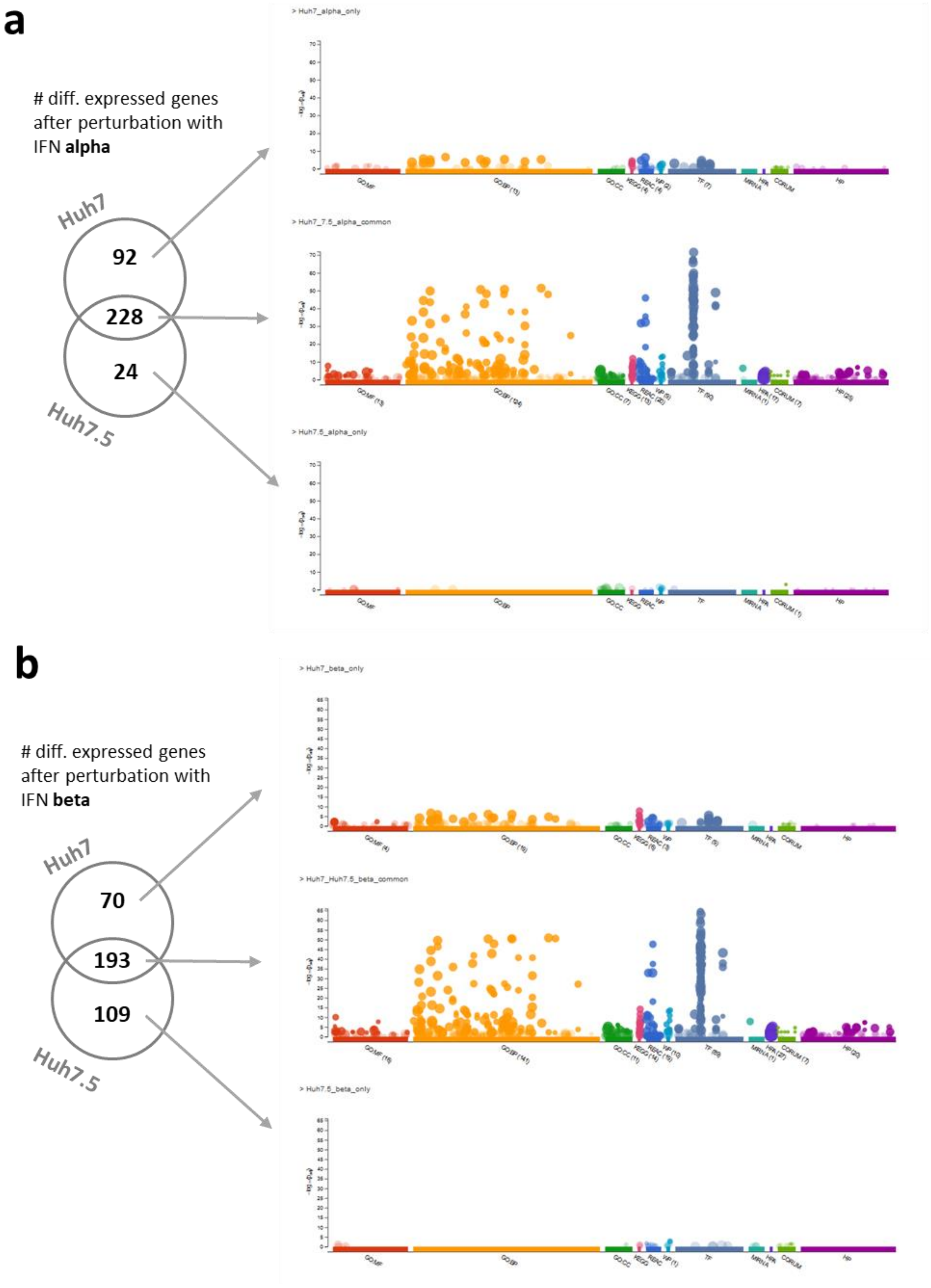


**Supplementary Figure 2**

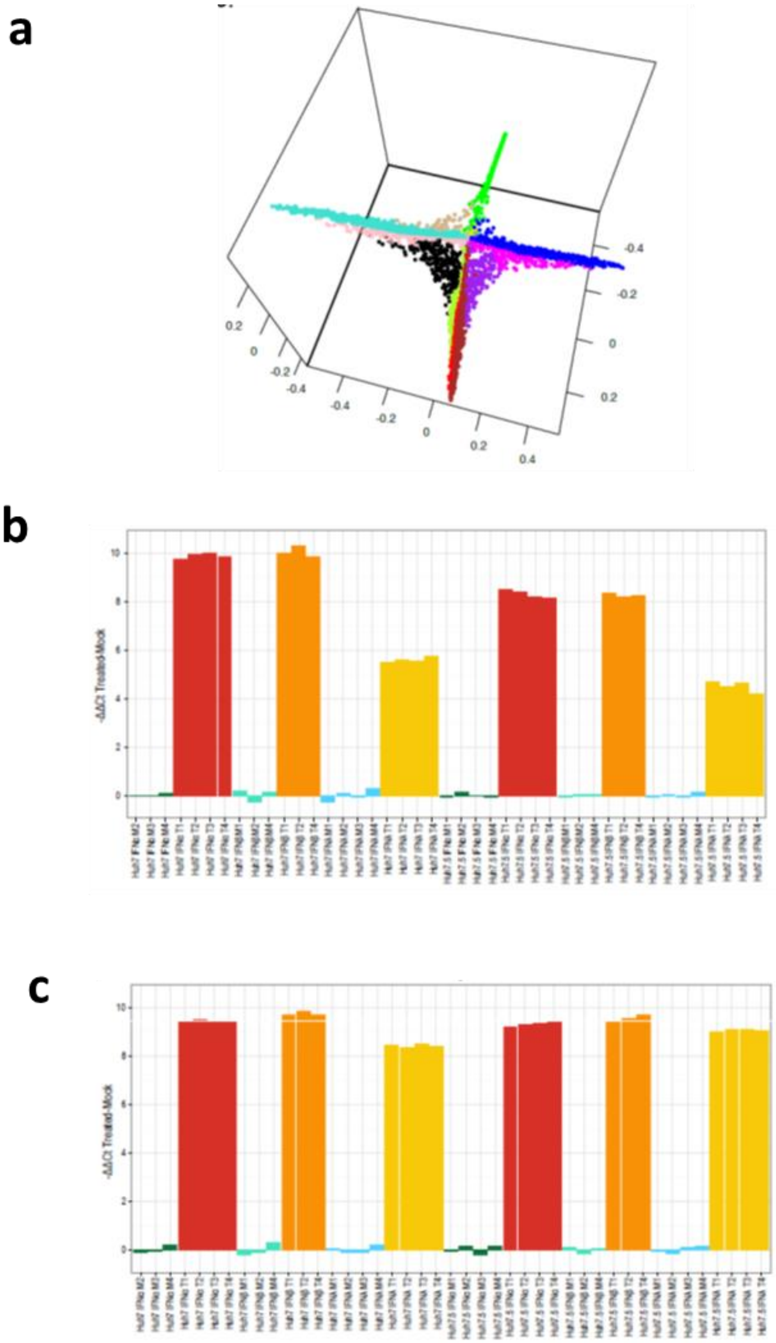


**Supplementary Figure 3**

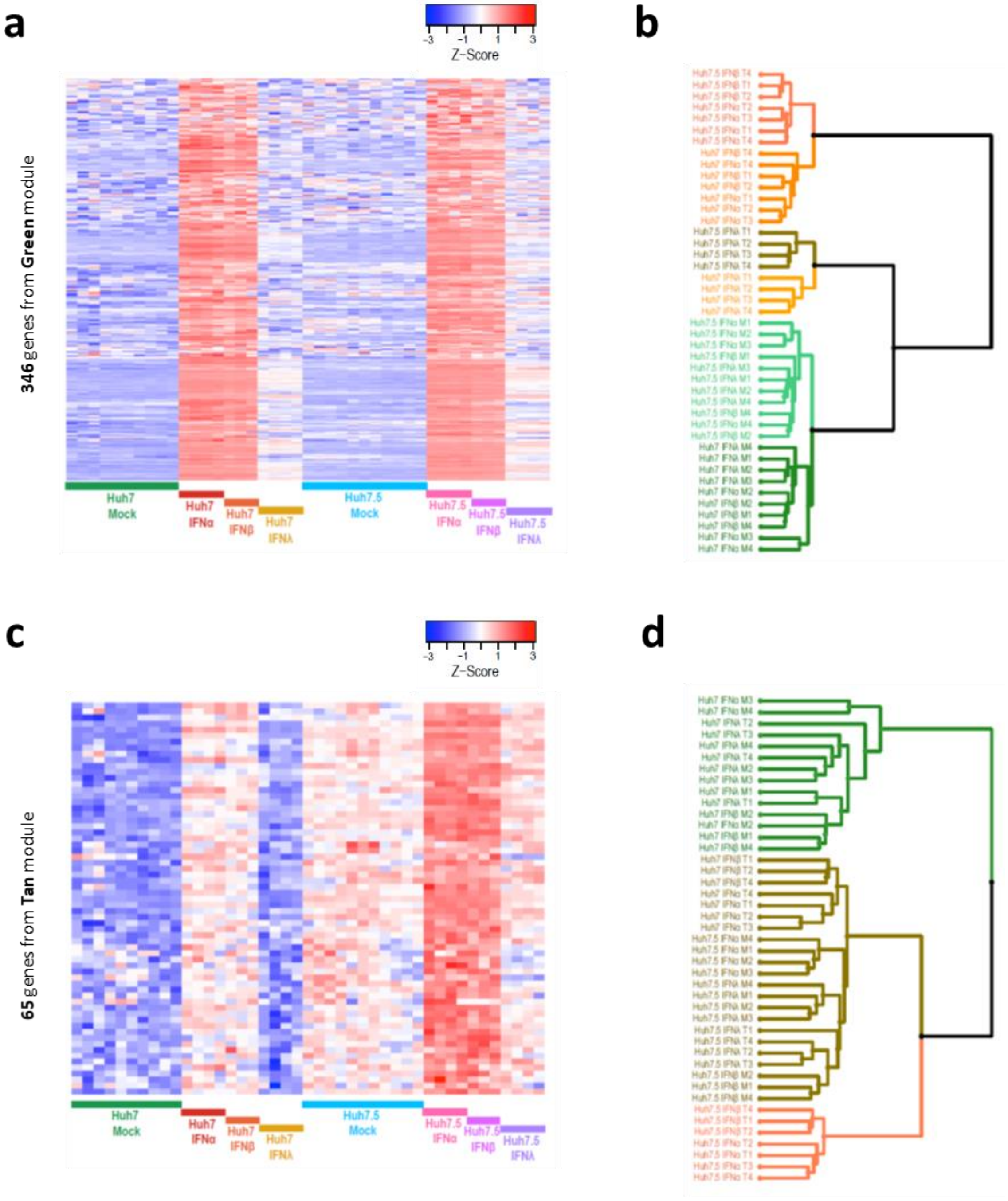


### Supplementary Figure 4

a

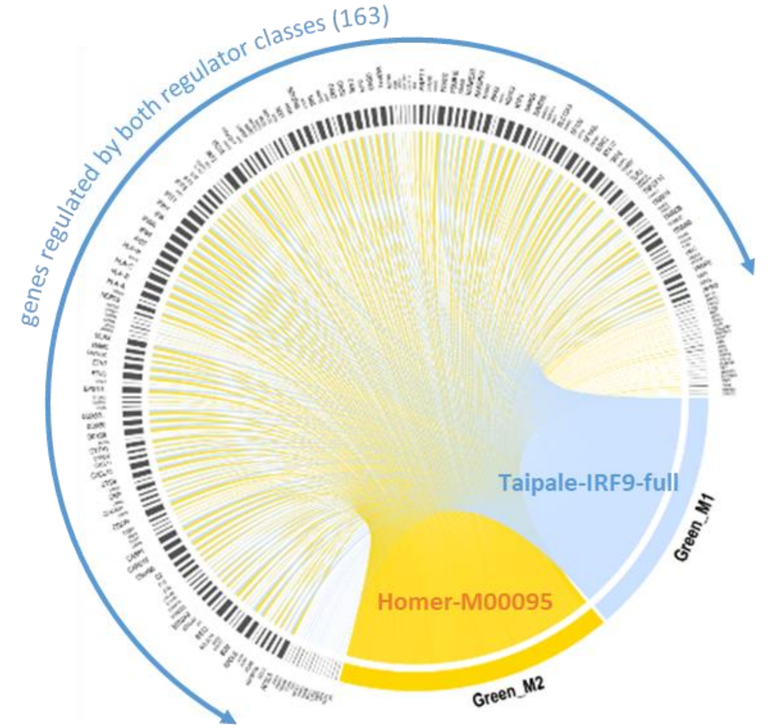


b

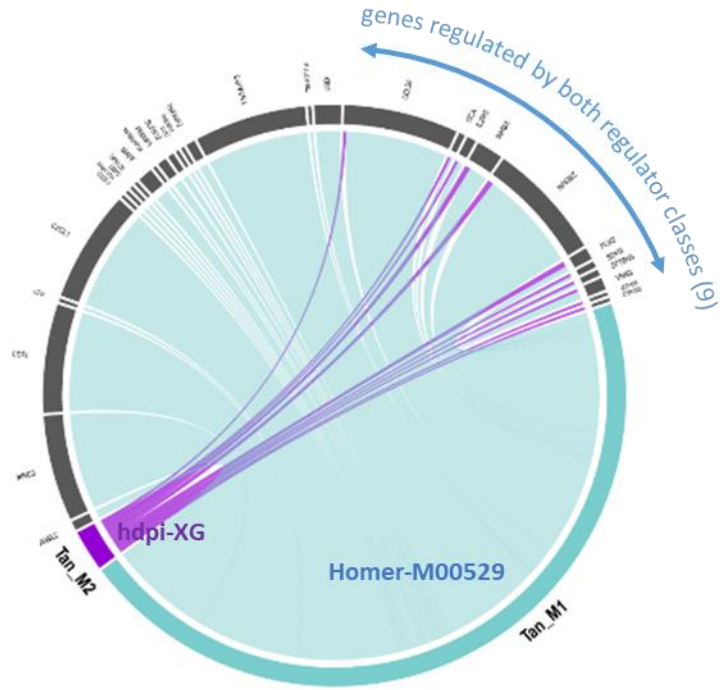


**Supplementary Figure 5**

a

b

c

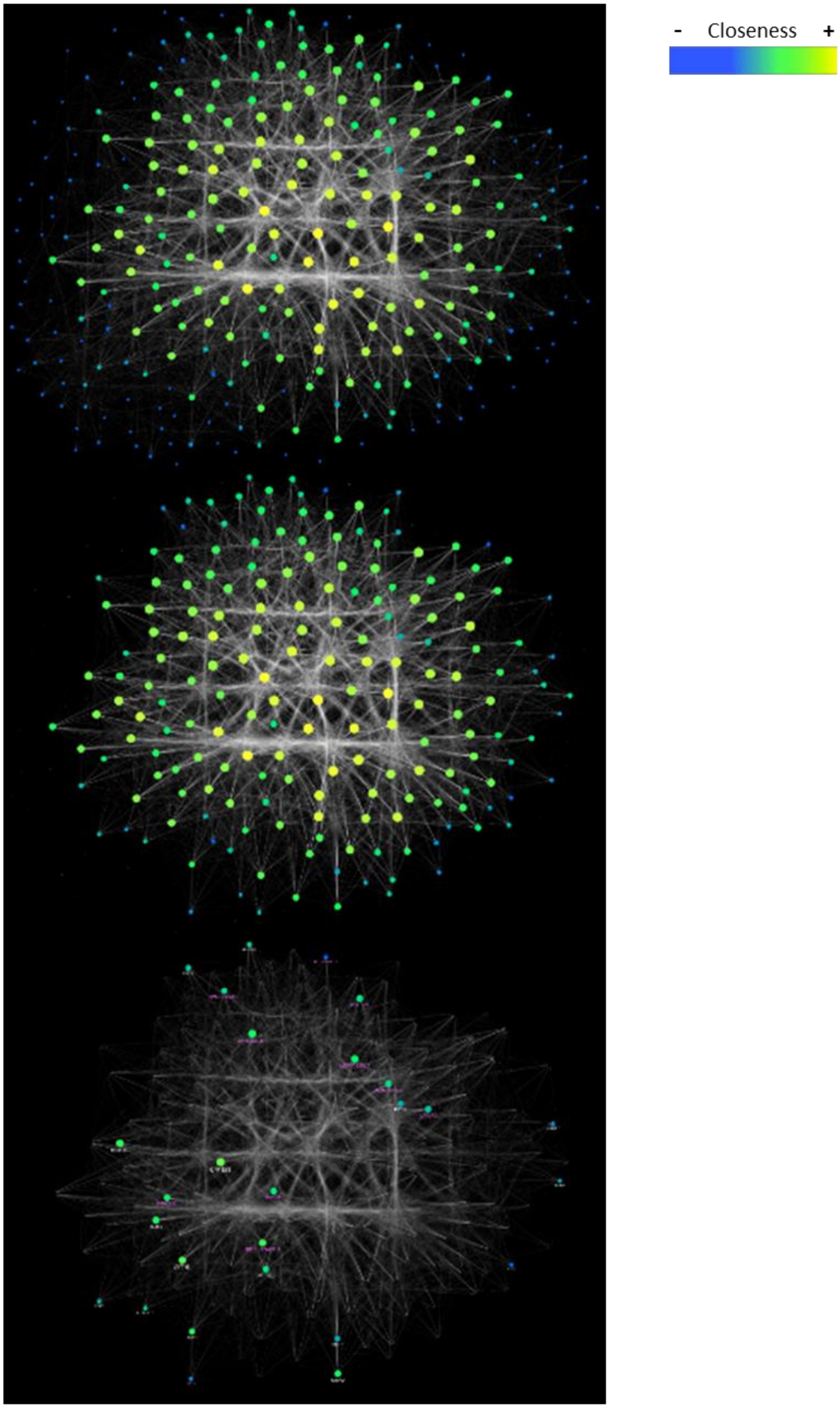


**Supplementary Figure 6**

a

b

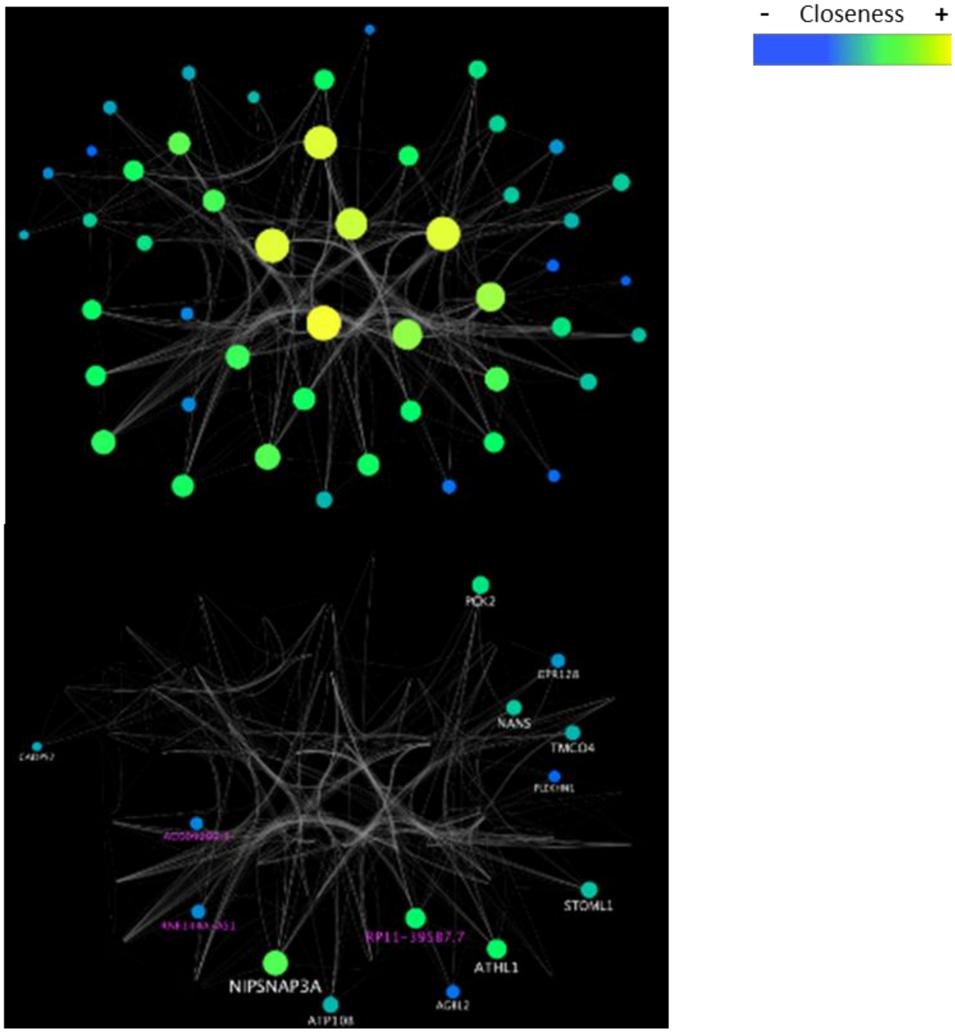


**Supplementary Figure 7**

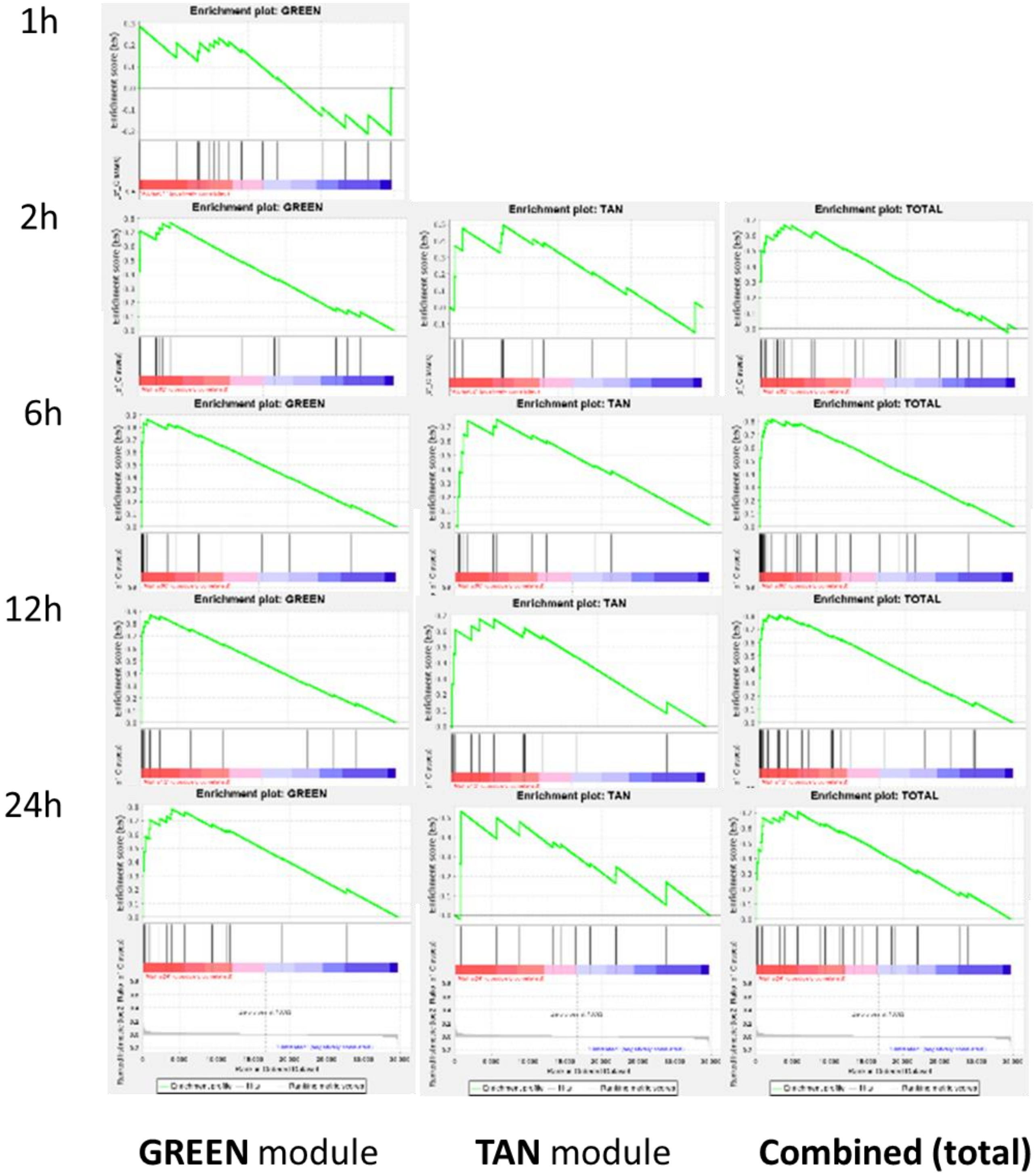


**Supplementary Figure 8**

1h

6h

12h

24h

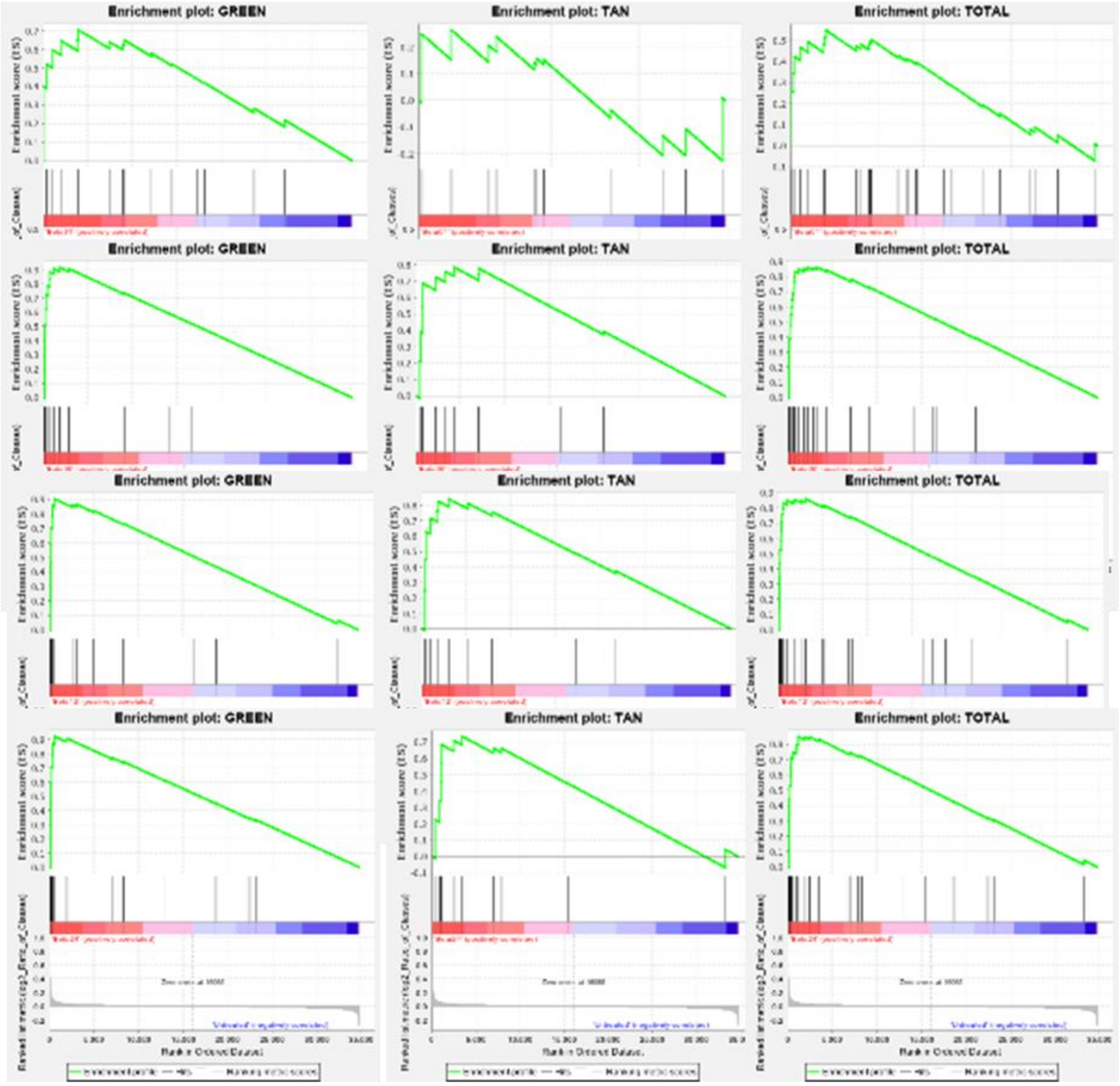


**GREEN** module **TAN** module **Combined (total)**

**Supplementary Figure 9**

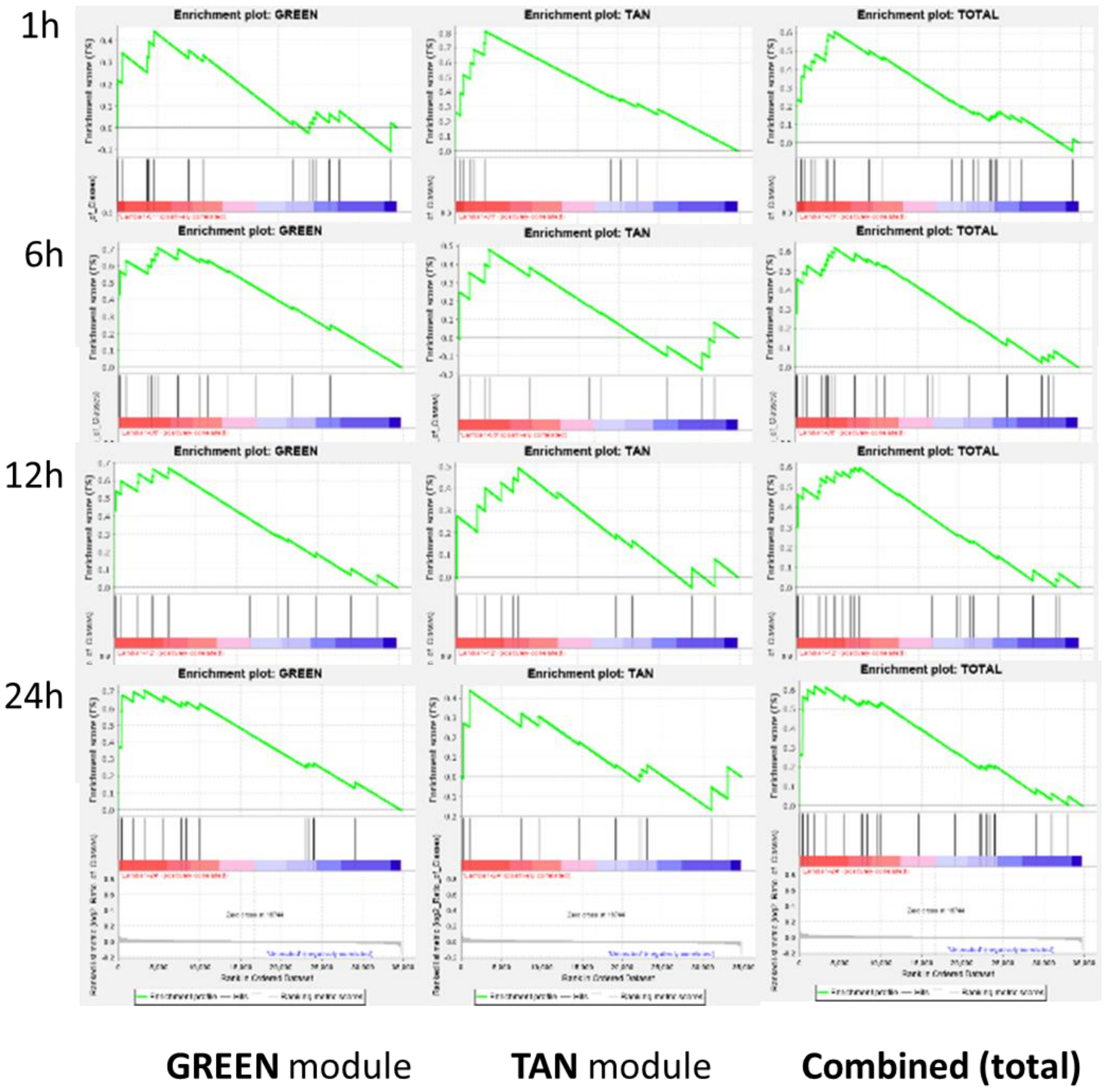


**Supplementary Figure 10**

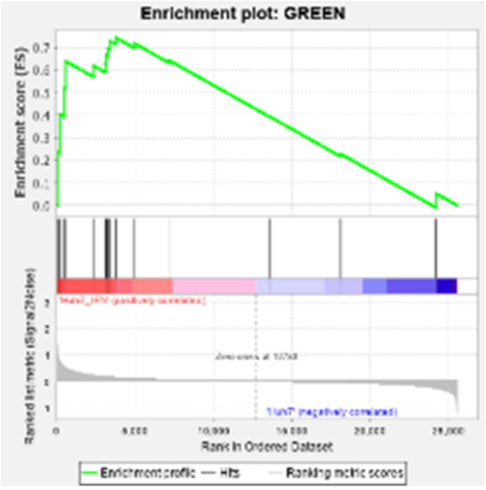


**GREEN** module

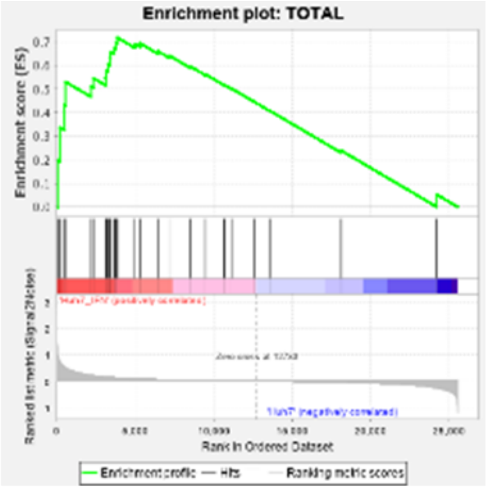


**Combined (total)**

**Supplementary Figure 11**

12h

24h

48h

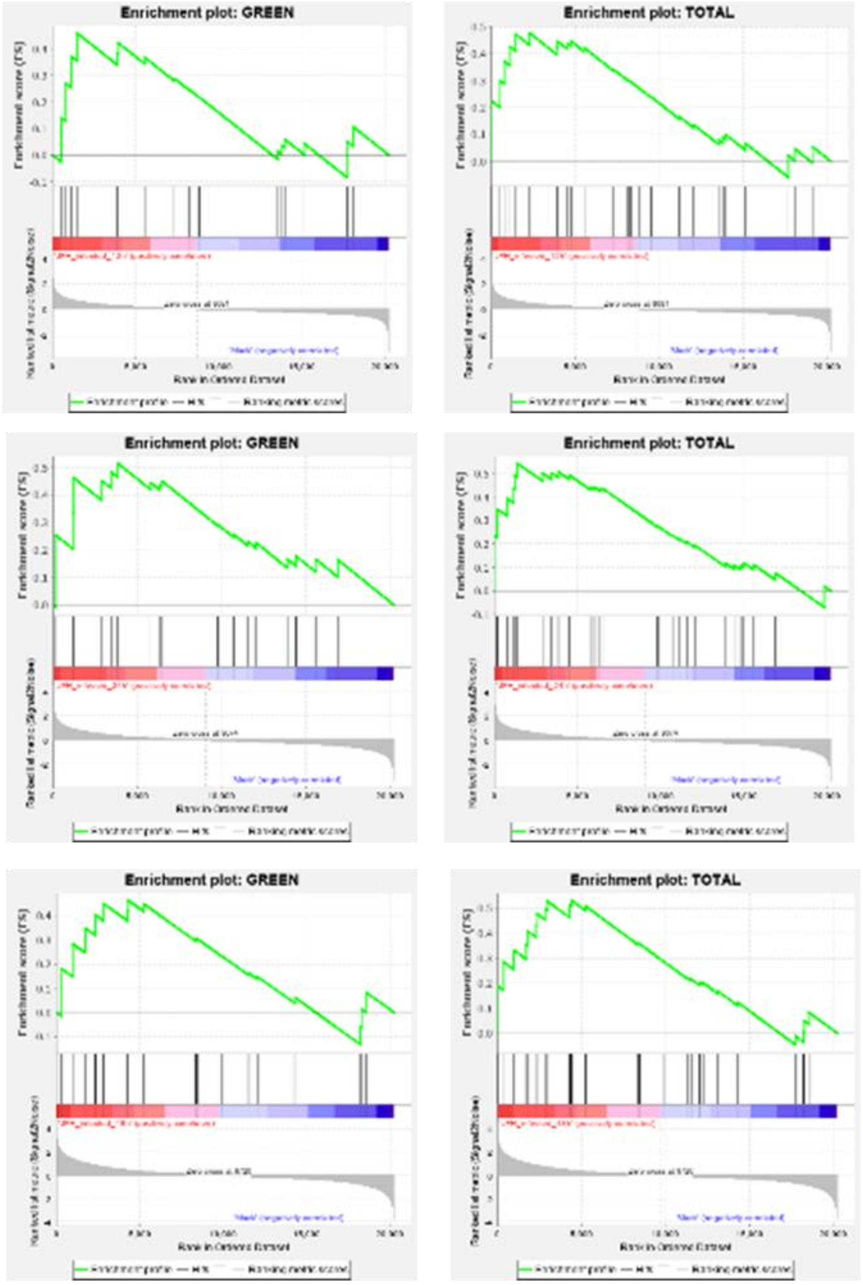


**GREEN** module **Combined (total)**

**Supplementary Figure 12**

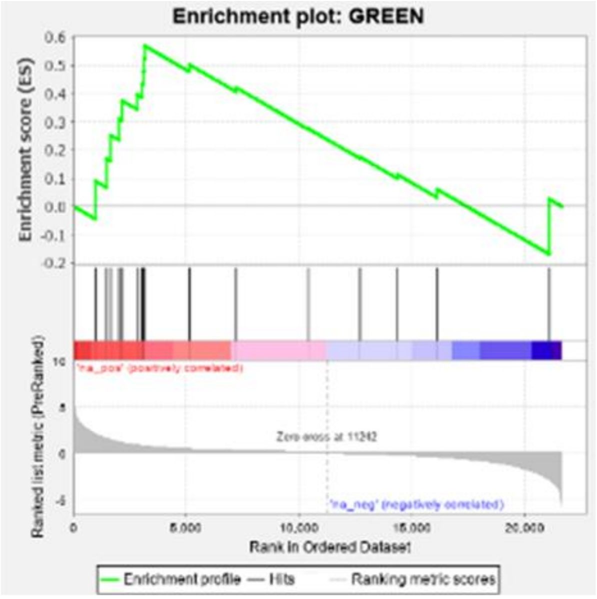


**GREEN** module

**Supplementary Figure 13**
